# Silent reservoirs are shaping the emergence of Usutu virus

**DOI:** 10.1101/2024.12.17.628855

**Authors:** Mariken M. de Wit, Gaël Beaunée, Martha Dellar, Emmanuelle Münger, Louie Krol, Nnomzie Atama, Jurrian van Irsel, Henk van der Jeugd, Judith M.A. van den Brand, Chantal B.E.M. Reusken, Marion Koopmans, Mart C.M. de Jong, Reina S. Sikkema, Quirine A. ten Bosch

**Affiliations:** Infectious Disease Epidemiology, Department of Animal Sciences, Wageningen University and Research, Wageningen, The Netherlands; Oniris, INRAE, BIOEPAR, 44300, Nantes, France; Institute of Environmental Sciences, University of Leiden, Leiden, The Netherlands; Deltares, Utrecht, The Netherlands; Viroscience, Pandemic and Disaster Preparedness Research Centre, Erasmus MC, Rotterdam, Netherlands; Vogeltrekstation-Dutch Centre for Avian Migration and Demography, Netherlands Institute of Ecology NIOO-KNAW, Wageningen, The Netherlands; Radboud Institute for Biology and Environmental Sciences, Radboud University, Nijmegen, The Netherlands; Dutch Wildlife Health Centre, Utrecht University, Utrecht, The Netherlands; Centre for infectious Disease Control, European and National reference laboratory for vector-borne viruses, National Institute for Public Health and the Environment, Bilthoven, the Netherlands; Division of Infectious Diseases and Immunology, Department of Biomolecular Health Sciences, Faculty of Veterinary Medicine, Utrecht University, Utrecht, the Netherlands

**Keywords:** mosquito-borne diseases, mathematical modelling, Usutu virus, multi-host disease

## Abstract

Disentangling contributions of different hosts to disease transmission is highly complex, but critical for improving predictions, surveillance, and response. This is particularly challenging in wildlife, with pathogens often infecting multiple species and data collection being difficult. Using the emergence of Usutu virus (USUV) in the Netherlands as a case study, we demonstrate the use of an Approximate Bayesian Computation framework on diverse data sources to uncover drivers of spatio-temporal wildlife disease emergence. We calibrated single- and multi-host mechanistic transmission models to five types of wildlife surveillance and research data, describing molecular and serological evidence of USUV in birds. Although Eurasian blackbirds, the primary target species for surveillance, were most severely affected, our models indicated that USUV could not persist in blackbirds alone. Our framework provided statistical support for additional, unobserved bird species to have contributed to transmission. This population of bird species is characterised by limited infection mortality, a longer lifespan, and likely further dispersal than blackbirds. Immunity in this population appears to have protected blackbirds from further USUV-related population decline. Our results underscore the importance of considering multiple host populations to understand outbreak dynamics. Neglecting the multi-host context of transmission can impact the reliability of predictions and projected impact of interventions.

## Introduction

The spread of many infectious diseases occurs through complex ecological interactions between humans, animals, and ecosystems [1,2]. Distinguishing the contributions of different host types to observed outbreak dynamics is crucial to inform predictions, early detection, preparedness, and outbreak response. However, disentangling contributions to disease transmission is notoriously complex. This is especially the case for zoonotic diseases where pathogens are capable of infecting multiple species. In zoonotic wildlife diseases, our ability to unravel these contributions is often hampered by the lack of high-quality information on population distributions, patterns of disease occurrence, and the lack of computational tools to integrate a wide range of data sources. Particularly during the emergence phase of a novel zoonotic pathogen, detailed outbreak data may be scarce and biased towards humans.

Contributions of different host species to transmission have been studied for several diseases and settings, such as minks and humans for SARS-CoV-2 [3], different bird species for avian influenza [4], cattle and wildlife for bovine tuberculosis [5] and wildlife reservoir hosts for plague [6]. Such analyses improve our understanding of the disease system and could also help inform and target surveillance strategies [7,8] and interventions, including vaccines where available [9–11]. Depending on each species’ contribution to transmission, interventions may need to target multiple host species as e.g., for brucellosis [12] and rabies [13]. A better understanding of host contributions can also inform risk of geographical expansion of emerging pathogens and future changes in disease burden of established pathogens.

A wide range of approaches for disentangling host contributions have been used, including the use of genetic sequencing [3,5] and combining mechanistic models with outbreak data [4,6]. However, these studies generally focus on the contributions of species that are known to contribute to transmission and use data specifically collected from these species, while it is also possible that relevant hosts are not (yet) identified. Such unidentified host species that influence disease dynamics constitute the so-called ‘epidemiological dark matter’ [14,15]. Developing approaches capable of assessing the role of this dark matter in shaping transmission has been described as one of the main challenges for understanding multi-host disease dynamics [14]. Moreover, transmission models are often calibrated to one or two specific outbreak characteristics such as prevalence over time or mortality patterns. Characterisation of unobserved reservoir hosts is more robust when models are calibrated to several outbreak patterns.

A prime example of a virus where multiple, unidentified, host species may contribute to transmission is Usutu virus (USUV). USUV is a mosquito-borne virus, closely related to West Nile virus, that emerged in the Netherlands in the past decade [16]. It was first detected in Europe in 2001 following an observed increase in bird deaths [17] and has since spread to several European countries [18]. The first detection of USUV virus in the Netherlands in April 2016 was followed by a substantial increase of dead Eurasian blackbirds (*Turdus merula*, hereafter: blackbird) [16,19]. Multi-year surveillance conducted in the Netherlands revealed striking emergence patterns (Figure 1) [20,21]. Firstly, USUV appeared to have spread from South to North between 2016-2018. Secondly, this period was followed by several years with fewer detections, after which the incidence increased again in 2022. USUV is transmitted in a cycle between *Cx. pipiens* mosquitoes and birds [22]. The virus has been detected in many bird species [20,22] and serological evidence of past exposure has been found in a wide range of animals [18,20]. Across Europe, the most affected bird orders are Passeriformes, Accipitriformes, Strigiformes, and Columbiformes, with infections occurring in both wild and captive birds [18]. The virus is also able to infect humans but does generally not lead to symptoms [18]. Nine countries in the EU have USUV surveillance schemes for animals, but target species vary between countries and available epidemiological and experimental data does not provide sufficient information on the role of bird species or mammals in the USUV life cycle [18].

**Figure 1.**
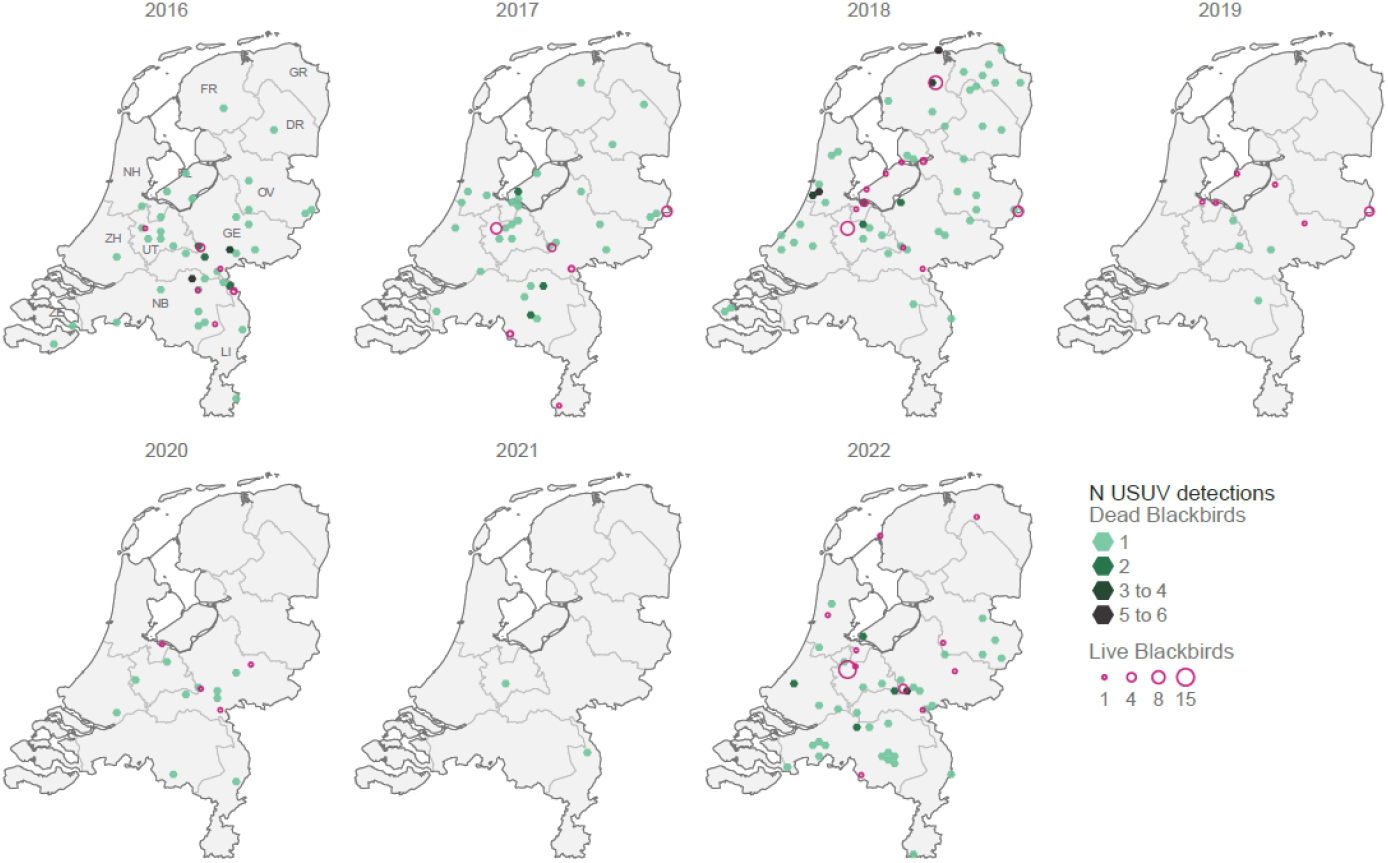
Spatio-temporal emergence pattern of USUV in the Netherlands, 2016-2022. Pink circles indicate detections in live birds, with sizes proportional to the number of detections. Green hexagons (size: 30 km^2^) indicate detections in dead birds, with shading intensity reflecting the number of detections in dead free-ranging birds in that cell. Provincial borders are shown and are labelled with two-letter abbreviation in the first map. The figure is adapted from a related figure in Munger et al. 2024 [20].

Blackbirds were the most commonly sampled species in the Dutch surveillance data between 2016-2022, showed the highest USUV prevalence among live birds, and showed the highest number of USUV-related deaths [20]. High viral RNA loads detected in multiple organs suggests blackbirds also contribute to onwards transmission [23,24]. However, several other animal species can experience infections with USUV. In this study, we aimed to explore whether, and to what extent, non-blackbird, reservoir hosts played a role in shaping spatio-temporal patterns of USUV emergence and spread. Secondly, we aimed to reconstruct the emergence and spread of USUV in the Netherlands and quantify the contributions of different host populations to this. To characterize the role of host populations other than blackbirds in transmission dynamics, we developed a Bayesian inference framework in which mechanistic USUV transmission models were fitted to several sources of blackbird surveillance data. Through comparing these models, we disentangled the characteristics of a potential unobserved reservoir population. Using the best-fitting model we reconstructed the emergence over time and space and quantified each population’s contribution to transmission over time.

## Results

### Trends in epidemiological surveillance data

Five sources of blackbird surveillance data were used to study USUV emergence between 2016-2022 [20,21]. Observations were filtered to align with the modelled transmission season (April to October). Surveillance data included infection prevalence (based on RT-PCR; n=2499) and seroprevalence in live blackbirds (n=1091), infection prevalence in dead blackbirds (n=284), blackbird population size, and reported dead blackbirds (n=3505). Infection prevalence in live blackbirds varied between years with the lowest observed prevalence in 2021 (0 positive cases) and highest in 2018 (0.09 (n=25/291) 95%CI 0.05-0.12). Prevalence in dead blackbirds was much higher and ranged from 0.10 (n=2/21, 95%CI 0-0.22) in 2021 to 0.87 (n=61/70, 95%CI 0.79-0.95) in 2018. Seroprevalence peaked in 2018 (0.23 (n=44/194) 95%CI 0.17-0.29), which coincided with the lowest blackbird population size being observed in the following spring. Most dead birds were reported in 2016 (29% (n=1014/3505) of the total across 2016-2022) (Supplementary figure 9).

### Characterising hidden reservoir populations

To explore the potential role of host populations other than blackbirds in the emergence of USUV we developed four compartmental transmission models with different reservoir population characteristics (Table 1) based on empirical data of mosquito- and bird abundance [25,26], bird movement, and local temperature [27], using a 5×5km grid cell structure. These models were fitted to the five surveillance datasets using Approximate Bayesian Computation by aggregating the grid cells into three regions (North, Middle, South). Models were compared based on the (normalised) distances between model predictions and observed data across the five datasets. Models that included an reservoir population (mean normalised distances 0.52-0.80) consistently outperformed the blackbird-only model (model A, mean normalised distances 0.97) (Figure 2E). Comparison of different reservoir population characteristics revealed that the observed patterns in blackbirds were best described by the presence of a reservoir population that does not die from infection and disperses further than blackbirds (model D). This model estimated high seroprevalence levels in the reservoir population (Supplementary figure 12) and was able to capture both the South-to-North patterns and the reduced transmission in later years (reflected in a stabilisation of blackbird population size, Figure 2A-D). It showed the best fit to the data across four out of five datasets (Figure 2E, mean normalised distances = 0.52, compared to 0.60 for B, and 0.80 for C).

**Table 1:**
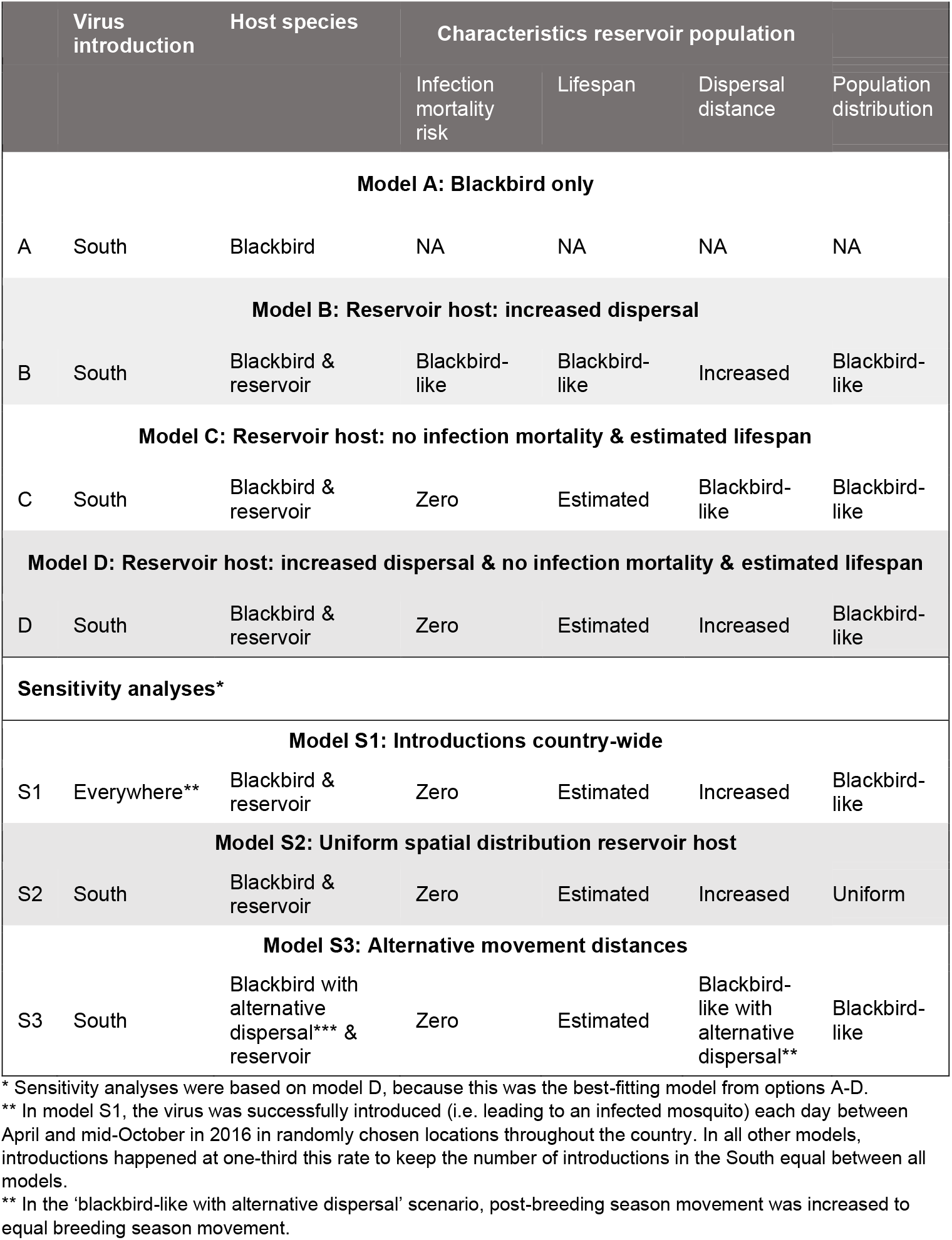
Specification of model versions.

**Figure 2.**
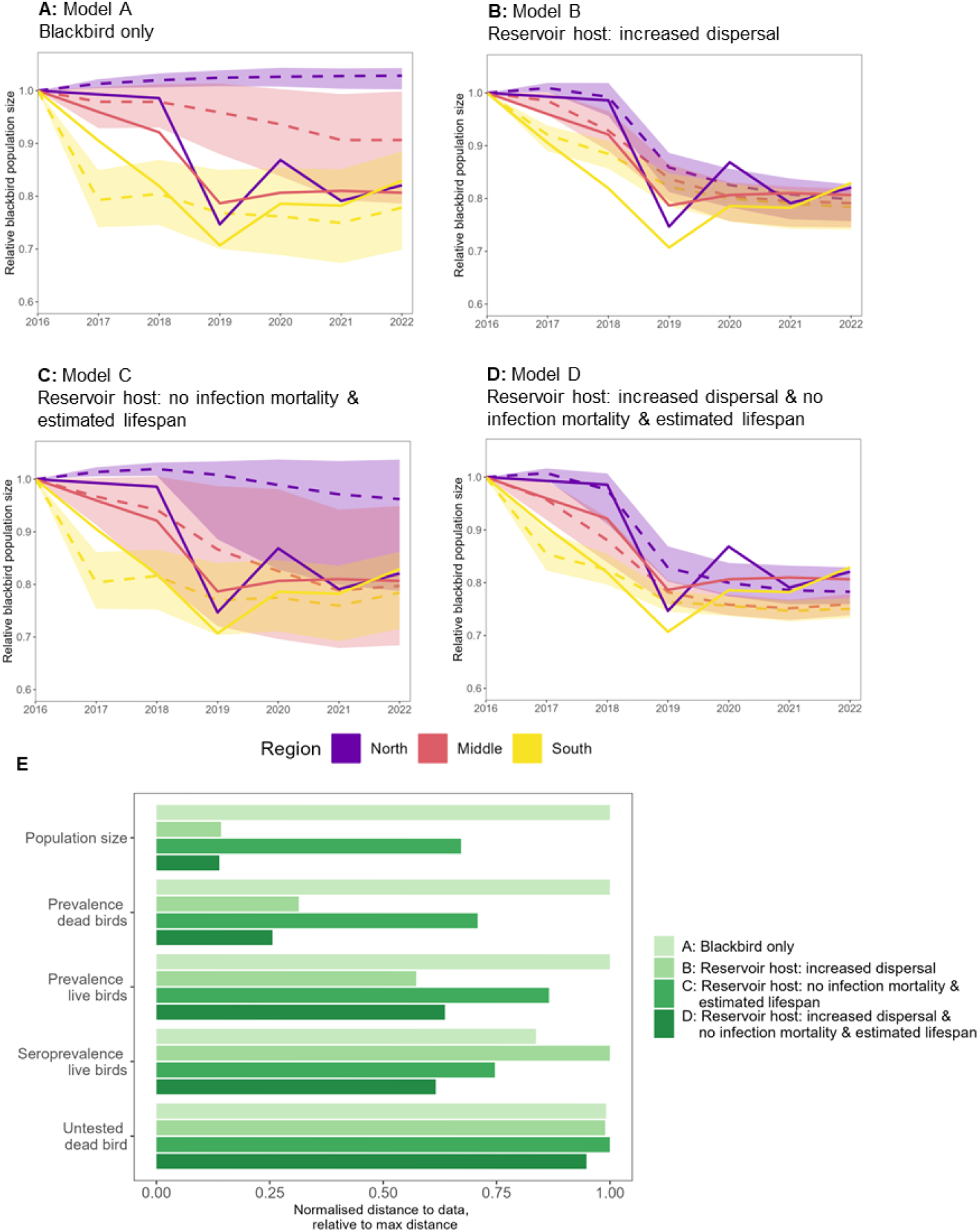
Comparison of model versions. A-D) Visual fit of simulated data (dashed lines) to observed data (solid lines) for the blackbird population size. Simulated data are presented as mean values with 95% prediction intervals. Results are shown for the four main models A to D. E) Comparison of model fits between main model versions. Comparisons were based on distances between observed and simulated data for each data set. For each data set, distances were normalised by dividing each model’s distance by the maximum across all model versions. A normalised distance of 1 thus indicates that this model showed the largest distance between observed and simulated data (i.e. worst fit). Lower normalised distances indicate a better fit to the data.

Sensitivity analyses showed that results of the best-fitting model (model D) were robust to assumptions regarding the location of virus introductions (mean normalised distance 0.53) and the spatial distribution of the reservoir population (mean normalised distance 0.54, Supplementary figure 13). Sensitivity analyses exploring the impact of uncertainty in blackbird movement distances (i.e., equal movement in breeding and post-breeding seasons) suggested that the increased dispersal required to explain the data falls within the uncertainty range of our blackbird movement estimates (mean normalised distance 0.55). Leaving out PCR data during model calibration resulted in posterior estimates that were similar to those obtained from the best-fitting model using all data (Supplementary Material I). All further analyses are based on the best fitting main model: model D. Inspection of model fits showed that this model was able to capture the trends in population size and immunity levels in blackbirds, but underestimated prevalence in 2017 and 2022 and did not capture trends in reported dead blackbirds (Supplementary figure 14). Model fits to within-year trends are shown in Supplementary figure 15, model fits of all other models are shown in Supplementary Figure 16-21, and further inference diagnostics can be found in Supplementary Material G.

Posterior estimates showed that the average adult lifespan of the modelled reservoir population was 5.8 years (95% CI 1.1 – 12.7). This population received 42 times (95%CI 32 – 50) more bites than the blackbird population per day, due to differences in population size and/or mosquito biting preference. Posterior distributions were largely robust across model versions showing high degree of overlap in 95% highest density intervals, especially across models D and S1-3 (Supplementary table 8).

### Reconstruction of USUV emergence

In addition to the role of reservoir populations, we also examined other determinants of the USUV outbreak. The best-fitting model showed an estimated infection mortality risk for blackbirds of 74% (95%CI 66-78). While the first RT-PCR-positive bird was found in 2016, undetected transmission prior to 2016 cannot be ruled out. We therefore aimed to estimate the level of immunity in 2016, but found that evidence of transmission prior to 2016 using our model was inconclusive (historical FOI 0.14, 95%CI 0.01 – 0.28). Additionally, we quantified the annual USUV re-emergence rate, representing the possibility of local overwintering and/or re-introductions from migratory birds. This rate was estimated at 18.0% (95%CI 6.5-29.6), indicating that USUV prevalence in mosquitoes at the start of the new season (April) is around 80% lower than at the end of the previous season (mid-September). Posterior distributions for all parameters and model versions are available in Supplementary table 8.

We used our best-fitting model to reconstruct the emergence of USUV into the Netherlands. Prevalence in live and dead blackbirds generally peaked in August and September, matching the trends in observed data (Figure 3A & B). In our model, highest monthly average prevalence was estimated in August 2018 at 3.0% (95%CI 2.3-3.7) in live birds and 67.1% (95%CI 60.6-72.5) in dead blackbirds. While the mean prevalence in live blackbirds tended to be highest in juveniles (0.7%, 95%CI 0.5-0.9, adults: 0.5%, 95%CI 0.3-0.7), prevalence in dead blackbirds was highest in adults (30.4%, 95%CI 23.4-37.6, juveniles: 17.1%, 95%CI 13.5-20.6). Seroprevalence generally peaked in October and was on average 5.1 times higher in the modelled reservoir population (68.1%, 95%CI 56.1-74.8) compared to seroprevalence in blackbirds (13.4%, 95%CI 9.7-17.9) (Supplementary figure 12), highlighting the role of reservoir population immunity in shaping transmission patterns.

**Figure 3.**
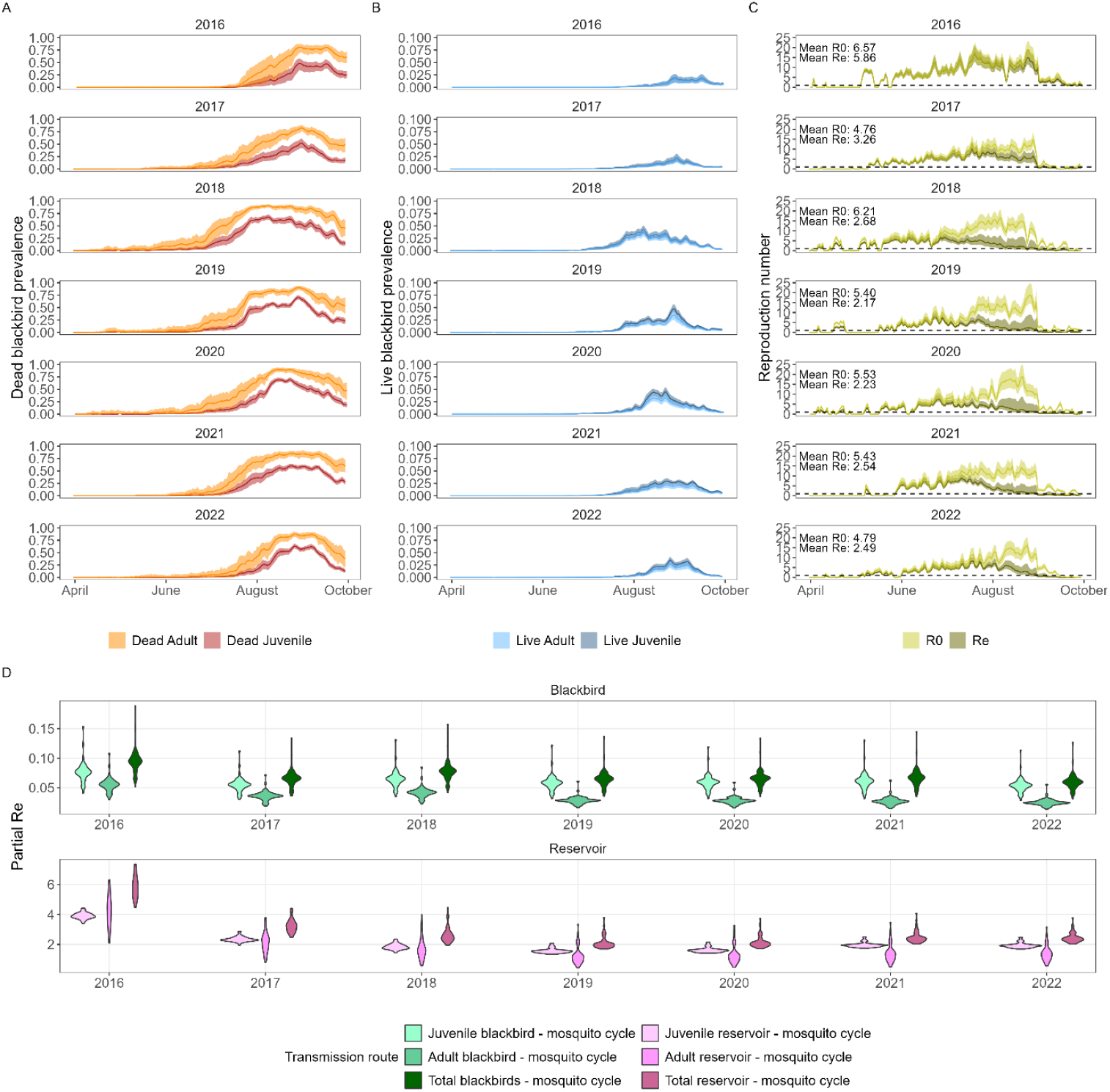
Model outputs generated from 100 simulations using random draws from parameter sets from posterior distributions. A) Infection prevalence during the transmission season in dead blackbirds split by age group. B) Infection prevalence during the transmission season in live blackbirds split by age group. C) R_0_ and R_e_ during the transmission season. D) Partial effective reproduction numbers per transmission route averaged over the transmission season. Results are shown per transmission season from April to October while averaging across all locations. In plots A-C, solid lines indicate mean values and shaded areas show 95% prediction intervals.

We calculated basic reproduction numbers (R_0_), indicating the number of secondary infections resulting from one infectious individual in a fully susceptible population, and effective reproduction numbers (R_e_), indicating the number of secondary infections when accounting for immunity in the population. The annual mean R_e_ was highest in 2016, when immunity was still limited, and lowest in 2019 (Figure 3C). The R_e_ tended to get above 1 at the end of May, peaked in July and dropped below 1 in early September, when mosquitoes started entering diapause. Differences in R_e_ between regions were small (R_e_: North 3.4 (95%CI 2.6-4.8), Middle 3.0 (95%CI 2.5-4.9), South 2.9 (95%CI 2.4-4.7)) (Supplementary figure 26-27).

### Contributions of different host populations to transmission

To disentangle contributions to transmission from the different host populations, we calculated reproduction numbers for each transmission cycle, so-called partial reproduction numbers (Supplementary figure 28). We found that the reservoir-mosquito cycle contributed more to overall transmission than the blackbird-mosquito cycle. The R_e_ of the blackbird-mosquito cycle during the transmission season was 0.07 (95%CI 0.05-0.11), meaning that USUV could not persist in blackbirds alone. With an R_e_ of 3.0 (95%CI 2.3-4.3), USUV could persist in the reservoir-mosquito cycle (Figure 3D). Juvenile blackbirds contributed about 1.5 times more to transmission than adult blackbirds (juvenile R_e_ 0.06 (95%CI 0.04-0.10), adult R_e_ 0.04 (95%CI 0.02-0.05) (Figure 3D).

## Discussion

In this study, we integrated five sources of surveillance data with data-driven transmission models in a Bayesian framework to assess drivers of spatio-temporal wildlife disease emergence. Using USUV as a case study, we showed that ignoring the multi-host context of transmission strongly impacts estimated outbreak dynamics. Specifically, failing to account for the build-up of immunity in reservoir populations hampers the model’s ability to accurately capture multi-annual trends. We found that while blackbirds were strongly impacted by the virus, they could not by themselves contribute sufficiently to the population-level build-up of immunity, nor to the spread needed to explain the spatial and multi-annual patterns of emergence. We used the inference framework to characterise the unobserved reservoir population as having a longer lifespan, low infection mortality, and likely a larger home range than blackbirds.

### Contributions from different host populations

We found that, during the first recorded outbreak of USUV in the Netherlands, one or more host species in addition to blackbirds played a role in transmission. Previous studies highlighted evidence of exposure in a wide range of animal hosts, most commonly in birds but also in mammals such as horses, dogs, and wolves [18,20,22]. Several papers have explored contributions of and interactions between host stages and/or species for the closely related West Nile virus from a theoretical perspective, highlighting the large impact that such interactions can have on outbreak dynamics [28–33]. However, this impact has not been demonstrated from real-life West Nile virus outbreak data. We found that the reproduction number for the blackbird-mosquito cycle was below 1 (maximum value 0.28 (95%CI 0.19-0.42)) during the whole transmission season, suggesting that USUV persisted due to the host competence (i.e. ability to transmit USUV onwards) and abundance of other populations. An important driver of the difference in contribution between the two host groups was the estimated distribution of mosquito bites. The proportion of bites on each host is a consequence of both host preference and relative abundance, which we could not disentangle in our analysis. Our estimated distribution of bites (1-2% on blackbirds) is consistent with observations in the field (0-9% of bites on blackbirds) [34]. However, this model-based estimate is sensitive to assumptions around transmission probabilities, which we assumed to be the same for the reservoir population as for blackbirds. It is important to note that these findings do not imply that USUV transmission can only be sustained in the presence of other competent bird species but that this depends on how many mosquito bites blackbirds receive, which is a function of the composition of the total population mosquitoes feed on [34,35].

### Immunity in the reservoir population

High level of immunity (reflected by high seroprevalence levels) in the reservoir population was found to be an important explanation of observed transmission dynamics in blackbirds. While the basic reproduction number remained high each year, the effective reproduction number, which accounts for immunity in the host population, reduced as the outbreak spread across the country. The strong decline in the effective reproduction number in the (adult) reservoir population in the first three years indicates that high immunity in this population reduced USUV transmission in later years and thereby protected the blackbird population from further decline. This is an example of the ecological concept of population immunity [36], that is, for instance, used in measles control where high population immunity is achieved through vaccination to protect vulnerable populations, such as those unable to develop protective immunity [37]. Seroprevalence could reach high levels in the reservoir population due to their longer lifespan. In accordance with life-history theory [38], longer lifespans were modelled to be associated with a lower annual birth rate (for example through smaller clutch sizes) and therefore a smaller number of naïve individuals that enter the population each year. Following ecological theory, if only a limited number of susceptible individuals enter the population each year, the resulting increase in effective reproduction number is limited. An Austrian study reported an USUV seroprevalence of 8.5% across several wild bird species in 2003-2004, but seroprevalence had increased to 54% even before the transmission seasons of 2005-2006 [39]. In Spain, mean annual WNV seropositivity levels ranged from 5 to 69% [40], in line with end-of-season seroprevalence levels estimated in Italy (7.8% −89.1%) [41]. These studies show that heterogeneity in reported seroprevalence levels is large, with estimates varying both between and within years. Observational studies showed large variation in USUV antibody waning rates [42,43]. Lifespan of the reservoir population also affects immune dynamics. Seroprevalence and reservoir population lifespan estimates (which showed large uncertainty in this study) could be improved as longer time series of surveillance data become available.

### Identifying the reservoir population

Existing literature can be combined with surveillance data available on other bird species to start identifying which species are represented in the reservoir population. Results indicated that high seroprevalence levels was an important characteristic of the reservoir population. Of the four species with more than ten samples in 2005 in an Austrian study [39], all showed USUV seroprevalence estimates of around 50% or higher (blackbirds, Eurasian blackcaps (*Sylvia atricapilla)*, Ural owls *(Strix uralensis)*, and European robins *(Erithacus rubecula)*). However, only a subset of these birds were live captured wild birds and a significant proportion were captive birds and/or sick, injured, or dead. In the Netherlands, annual seroprevalence levels of that order of magnitude were not observed in any species, but species-specific sample sizes were often limited and estimates are sensitive to the timing of sampling. Highest seroprevalence levels among commonly sampled species in 2016-2022 were found in Carrion crows *(Corvus corone)* (13.7%, n=51, some of which were sick), Eurasian magpies *(Pica pica)* (14.1%, n=64), and Eurasian collared doves *(Streptopelia decaoto)* (11.1%, n=36) [20], of which the first two are relatively highly abundant in North-Western Europe [44]. Another important characteristic of the reservoir population was limited infection mortality and further dispersal. While the impact of USUV infection is unknown for most of these species, studies indicated that magpies do not show clinical symptoms after USUV infection [45]. Magpies also live somewhat longer than blackbirds, typically five years after reaching breeding age [46]. Other species found positive for USUV in the Netherlands disperse further, such as crows and doves [47,48]. Finally, to act as a reservoir host, species should be able to transmit the virus to mosquitoes and contribute to persistence. Experimental studies have shown that house sparrows can transmit USUV to mosquitoes and domestic canaries develop high viral RNA loads after infection, suggesting these species could act as reservoir hosts provided their transmission cycle is sufficiently efficient [49,50]. In conclusion, enhancing sampling in species with high seropositivity levels, experimental infections to assess competence and infection response, supplemented by demographic characteristics, could shed more light on the identity of species constituting the reservoir population.

### Temporal trends in transmission

We were able to reproduce the large outbreaks observed during the first three years of emergence, after which transmission was sustained at a lower intensity. Our model captured the observations that the virus spread from South to North in a period of three years. This spatio-temporal pattern was a result of the location of virus introduction, dispersal distances, and duration of the transmission season. We estimated that the effective reproduction number generally dropped below one after early September. Variation in basic reproduction numbers within and between years were mainly driven by mosquito population size and mean daily temperature, and to a smaller extent by bird population size. Spatial heterogeneity in reproduction numbers was limited and we did not identify clear transmission hotspots. Reproduction numbers were estimated to be highest in 2016 and 2018, coinciding with the highest estimated mosquito abundance in 2016 and highest spring and summer temperatures in 2018. In 2016, a large USUV outbreak was observed in Germany during which the virus spread towards North Rhine-Westphalia [51,52], a region to the southeast of the Netherlands. The high reproduction numbers in 2018 coincided with the largest observed WNV outbreak to that date in Europe [53]. Transmission season was shortest in 2021, where the reproduction number was not consistently above one until June, coinciding with the coldest spring of the study period. While some of the surveillance data sources showed increased transmission in 2022 compared to the previous years, this was not captured by the model, mostly because predicted mosquito abundance was lowest during this year. Limitations that may have impacted the predicted annual variation in transmission include the simplifying assumption of a constant number of newborn birds each year, while variation might occur due to environmental conditions [54], the assumption of a constant virus re-emergence rate each year, and remaining biases in surveillance data. We estimated the re-emergence rate (around 20%) as the proportion of the USUV prevalence in mosquitoes that persists to the next spring, representing the possibility of local overwintering and/or re-introductions from migratory birds. While we estimated a constant rate, this may be affected by winter temperatures [55] and variation in bird migration patterns, among others. Quality of model fit varied between datasets. While this could reflect epidemiological or ecological processes that were not captured in the model, such as those mentioned above, this was likely largely driven by imperfect observation processes such as non-constant sampling intensity and varying detection probabilities, especially because the observation process was sometimes linked to circulation levels.

### Implications for surveillance and predictions

Our findings indicate that blackbirds are useful as a target species for USUV surveillance because of their high infection mortality rate, which makes it possible to detect signals indicating large outbreaks in the absence of live bird surveillance schemes. The high prevalence in dead blackbirds makes sampling of dead blackbirds an efficient surveillance strategy to detect circulation. However, to better identify which other species can contribute to transmission, frequent sampling of a wider range of species is necessary. Longitudinal serological surveillance would be the most appropriate starting point to identify reservoir species, as they might not display increased mortality. However, to distinguish reservoir species (i.e. those capable of transmitting the virus onwards) from dead-end host species (i.e. those unable to further transmit), infectiousness would have to be confirmed for example using proxies such as viral RNA loads. Such information could be useful when making projections about future transmission risks and when predicting the suitability for USUV circulation. The large contribution of non-blackbird host species implies that also regions with low blackbird density may well be suitable for USUV transmission, but this might remain undetected if surveillance is focused on dead bird reporting, as is the case in many other European countries. Additionally, the model could be employed to evaluate current surveillance strategies by adding these into the model and provide recommendations regarding sampling intensity, timing, and location for different surveillance systems.

### Importance of multiple data sources

Using multiple sources of data for inference is one of the key challenges in epidemiological modelling [56]. We addressed several obstacles related to synthesising diverse datasets. A common challenge in the use of multiple datasets for inference is weighing observations that are not on the same scale. We addressed this by defining acceptance thresholds for each dataset individually thereby placing equal weight on each dataset and only accepting parameters that simultaneously met the threshold for all datasets. This allowed for the model to be fitted to multiple aspects of an outbreak, representing different characteristics of the disease system, including population impact, infection levels in both live and dead hosts, and immunity build-up. Additionally, these data sources represented varying levels of temporal aggregation of the system. By simultaneously calibrating to multiple datasets, the impact of bias in one of the datasets and the risk of overfitting is reduced. This is illustrated by the observation that removing PCR data from the model inference had little impact on parameter estimates, likely because parameters were sufficiently informed by patterns in other datasets, although the effect of removing datasets likely varies between dataset. As a consequence of calibrating to diverse datasets simultaneously, the resulting model could not capture all individual observations when these were not consistent across all datasets. The use of summary statistics, in contrast to likelihood functions, increased flexibility in the definition of optimisation function and these could therefore be designed such that they are more robust to imperfect sampling compared to the full dataset. The inference validation analysis showed that our framework was able to correctly estimate unknown parameters from simulated data. These approaches enhance the usability of datasets, that would be insufficient on their own to address specific epidemiological questions. While more complete and larger datasets would be recommended for such analyses, this is unrealistic in the context of wildlife surveillance. With the increasing availability of novel surveillance data streams, possibilities to collect multiple types of data on the same outbreak are expanding. This highlights the importance of developing and evaluating computational approaches capable of integrating different data types.

Our results emphasise the importance of considering multiple host populations to understand outbreak dynamics. Ignoring the multi-host context of disease transmission can have implications for the projected effectiveness of intervention strategies or accuracy of future predictions. Advances in novel data streams and modelling techniques present highly promising developments in our ability to unravel transmission processes and characterise ‘epidemiological dark matter’.

## Methods

### Surveillance data

Five sources of USUV surveillance data were collected on blackbirds: 1,2) PCR and seroprevalence in live blackbirds, 3) reported dead blackbirds, 4) PCR prevalence in a subset of these dead blackbirds, 5) annual blackbird population size. We used these to calibrate mechanistic transmission models to better understand the emergence and spread of USUV in the Netherlands. Further details of these data sources can be found in supplementary material C and in [20,21].

### Model overview

We created a stochastic compartmental metapopulation model to simulate transmission of USUV between *Culex pipiens* mosquitoes and blackbirds in the Netherlands from 2016 to 2022 (Figure 4). Both the host and vector populations were divided into compartments based on their infection status: susceptible, exposed, infected, recovered, and dead compartments for birds, and susceptible, exposed, and infected compartments for mosquitoes. Blackbirds were chosen as the main host species for this study, because they showed highest USUV prevalence among live birds [20] and showed high mortality following USUV transmission [16,19]. To explore the potential role of reservoir hosts, we also created model variants that included an additional host population, representing unidentified USUV-competent species (i.e. species that can transmit USUV onwards). Birds can get infected when they get bitten by an infectious mosquito. Mosquitoes can get infected when they bite an infectious bird. We assumed host frequency-dependent transmission. This follows from the assumption that the vector biting rate is independent of host density, and the biting rate experienced by hosts increases with vector density [57,58]. The mosquito population dynamics, extrinsic incubation period, and biting rate were temperature-dependent. The model’s starting conditions were determined by the historical FOI (see ‘Estimated parameters’).The full set of equations and parameter values are described in Supplemental Material A.

**Figure 4.**
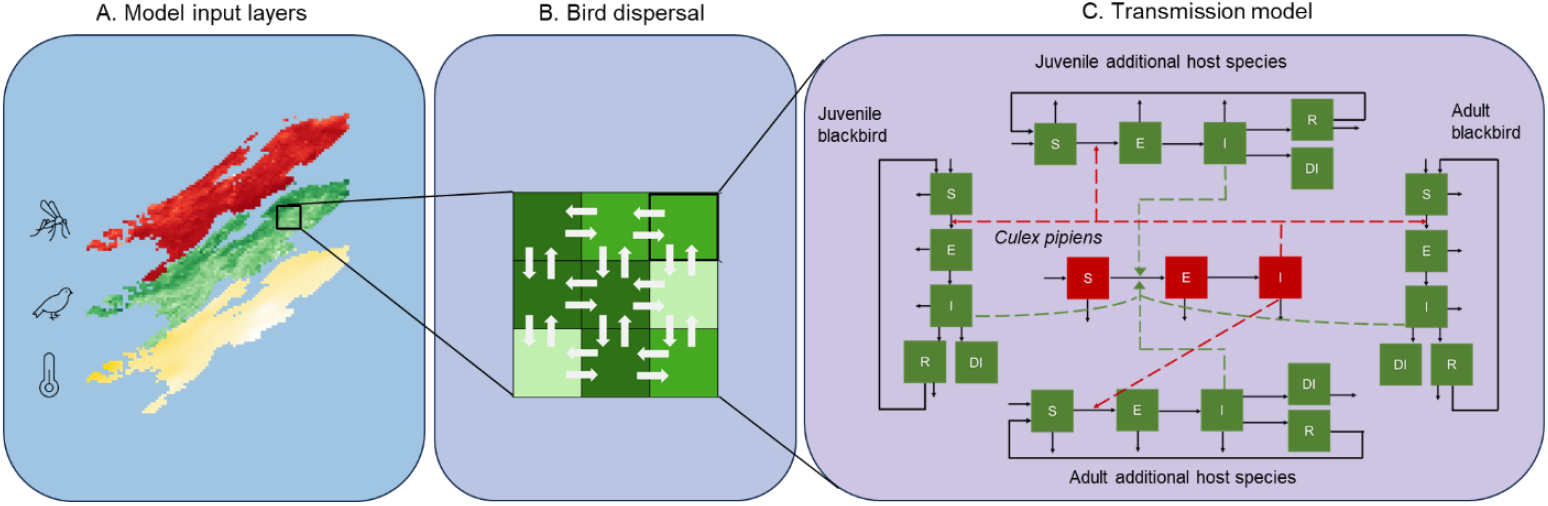
**Model overview**. The country is divided into equally sized 5×5km grid cells, which vary with respect to bird and mosquito abundance as well as temperature. Birds disperse between cells. Transmission occurs within a grid cell. In the SEIR-model, dark arrows represent transitions between infection states, while dashed arrows represent the link to the source of transmission. S = susceptible, E = exposed, I = infectious, R = recovered, DI = dead from infection. Non-connected outward pointing arrows represent natural death of the animal. Non-connected inward pointing arrows represent births.

The temporal resolution of the model was one day. We used a 5 by 5 km grid structure with bird and mosquito abundance varying between grid cells (see ‘Input layers’). Virus transmission occurred within a grid cell and cells were connected through bird movements. The transmission season was set from April to mid-October, with no transmission occurring outside this period. We distinguished daily foraging movement patterns from seasonal natal and breeding dispersal [59,60]. To allow for different movement patterns across life stages and seasons, we divided the transmission year into breeding season (April-July) and post-breeding (July-October) and the bird populations into juveniles (from fledging until the next breeding season) and adults (birds older than one year). Only bird movement during the virus transmission season was included, as movement outside this season was assumed not to affect transmission. Similarly, blackbird spring and autumn migration was not included explicitly because this largely occurs outside the transmission season and Dutch blackbirds have almost completely stopped migrating over the past decades [61]. Movement of mosquitoes between cells was ignored due to a lack of high-quality evidence available for parameterisation [62]. The model was run continuously from April 2016 to the end of 2022, with the virus re-emerging in the mosquito population each year, representing the possibility of local overwintering and/or re-introductions from migratory birds. All analyses were performed in R 4.2.1.

### Input layers

#### Bird and mosquito abundance

Relative blackbird abundance was obtained for each grid cell from Dellar et al. [26]. In this paper, Random Forest models were fitted to blackbird point count data (n=183,123 observations) from the Netherlands (Meetnet Urbane Soorten [63], Meetnet Agrarische Soorten [64] and Common Bird Census [65] from Sovon Vogelonderzoek Nederland (Dutch Centre for Field Ornithology)) and France (Common Bird Monitoring Scheme [66]) in the period 2001-2016, using a set of environmental and climatic predictors. Relative *Culex pipiens* abundance was obtained for each day and each grid cell from Krol et al. [25]. Here, data on female mosquito trap counts was obtained from the National Mosquito Survey 2010-2013 [67] and the MODIRISK project [68]. Random Forest models were fitted to female mosquito count data including a set of land cover, environmental, and climatic predictors. Estimates for both blackbird and mosquitoes reflect relative abundance and were scaled proportional to each other using a scaling parameter (see ‘Estimation parameters’).

#### Bird movements

Two movement patterns were distinguished in the model, representing daily movements and seasonal dispersal. Daily movements include foraging-type movements often within the bird’s home range, i.e. the area the bird regularly traverses around in its territory. In the model, this is reflected as a contribution to the FOI experienced in other grid cells. In addition to daily movements, we included seasonal dispersal that represents the search for a new nest location. Seasonal dispersal takes place during the change from breeding to post-breeding season in early July, whereby birds can move to a different grid cell. We quantified both the daily and seasonal blackbird movement patterns by fitting dispersal kernels to ringing and recovery data of blackbirds ringed and recovered in the Netherlands collected by Vogeltrekstation NIOO-KNAW. This resulted in a median daily movement distance of 143 m (inter-quartile range 19 - 692) during the breeding season and 29 m (inter-quartile range 4 – 129) during the post-breeding season. Seasonal dispersal distances were larger and estimated at a median distance of 377 m (inter-quartile range 95 - 1114) for juveniles and 180 m (inter-quartile range 19 – 1062) for adults. We constructed movement matrices by sampling from these dispersal kernels assuming rotational symmetry (Supplementary Material B). For the additional reservoir population we explored an additional movement scenario with a median distance of 1,016 m for breeding season and 323 m for post-breeding season (Supplementary Material B), seasonal movement was not included.

#### Temperature data

Average daily temperature data in the period 2016-2022 were obtained from the Royal Meteorological Institute (KNMI), with a 1×1 km spatial resolution [27].

Abundance and temperature data were resampled into the model grid using bilinear interpolation. A visual overview of bird and mosquito abundance as well as monthly temperature values can be found in Supplementary Figure 1.

### Inference

#### Approach

We used an Approximate Bayesian Computation approach using a Sequential Monte Carlo sampler (ABC-SMC) to calibrate our model to surveillance data [69]. This approach allows for complex transmission models to be fitted to data by using summary statistics, thereby avoiding having to explicitly define likelihood functions [70,71]. In ABC-SMC, estimation of the posterior distribution is achieved sequentially by constructing intermediate distributions in each generation, converging towards the final posterior distribution. After each model iteration using proposed parameters, model output is compared to the data using summary statistics. Summary statistics each represent different epidemiological aspects of the outbreak. The parameter set is accepted if the distances between the model and data summary statistics are below their respective thresholds, where the thresholds become progressively stricter each generation based on results from the previous generation. For each model, we ran three chains of 200 particles and stopped when the acceptance rate reached 5% or less.

We used five summary statistics, one for each type of surveillance data: seroprevalence in live blackbirds, PCR prevalence in live blackbirds, and PCR prevalence in dead blackbirds for each month and region (North, Middle, South (Supplementary figure 8)) as well as the annual blackbird population size relative to 2016 and the relative number of reported dead blackbirds per year within each region (Supplementary figure 9). In the model, regions were created by aggregating the grid cells. More details on the inference approach can be found in Supplementary Material D.

#### Estimated parameters

The following parameters were estimated: 1) abundance scaling parameter, this converts the host and vector abundance to be on the same scale, meaning that their ratio reflects the vector-to-host ratio, 2) re-emergence rate, the fraction of the USUV prevalence in mosquitoes in September that persists to the subsequent spring, 3) infection mortality rate in blackbirds (ν), 4) transmission probability bird-to-mosquito (p_mb_), 5) historical FOI, to quantify transmission prior to first detection leading to non-zero immunity levels in 2016. For the model versions including a reservoir population, also this population’s natural mortality rate (μ_AR_) and the distribution of mosquito bites between blackbirds and the reservoir population were estimated (ω). For a more detailed description of the parameters, including the priors, see Supplementary Material D.

### Validation of inference approach

To validate the ability and accuracy of our inference algorithm, we simulated data mimicking the surveillance data available with known parameter values using the blackbird-only model. We evaluated the ability of our model framework to estimate these parameter values using five simulated datasets (Supplementary Material E). We found good agreement between true values and posterior medians (concordance coefficients 0.55-0.97) for the re-emergence rate, abundance scaling parameter, transmission probability bird-to-mosquito, and infection mortality rate. Identifiability of historical force of infection (FOI) was poor (concordance coefficient <0.1). This parameter was still kept to ensure that the uncertainty around historical transmission is reflected in model results.

### Model comparisons and analyses

Four models (A-D) were created, which differed with respect to the presence and characteristics of an reservoir population (Table 1). Because there is no general consensus on the definition of reservoirs, we use a broad definition encompassing a population of animal species capable of (indirectly) transmitting infections to a target species (here: blackbirds) and enabling persistence of the virus in the ecosystem. We assumed the virus was introduced in the South as that appeared to be the first region where transmission occurred. Additionally, we developed three sensitivity analyses (models S1-S3) based on the best-fitting model to explore sensitivity to the location of initial virus introductions (S1), to the spatial distribution of the reservoir populations (S2) and to uncertainty in our blackbird dispersal estimates (S3). In model S1, the virus was successfully introduced (i.e. leading to an infected mosquito) each day between April and mid-October in 2016 in randomly chosen locations throughout the country. In all other models, introductions happened at one-third this rate to keep the number of introductions in the South equal between all models. We also explored the impact of removing the PCR data from the surveillance data and only fitting to the remaining three datasets (Supplementary Material I). Models were compared based on their distances between observed and simulated summary statistics calculated in the last generation of the ABC algorithm, where accepted particles represent the best approximation of the posterior distribution. To ensure all summary statistics’ distances were on the same scale, for each summary statistic separately we normalised the summed distances by dividing these by the maximum summed distances across all model versions.

Using the best fitting model, we ran 100 simulations using sampled parameter sets from the posterior distributions and calculated prevalence, seroprevalence, and (partial) basic (R_0_) and effective (R_e_) reproduction numbers for each day and grid cell. Reproduction numbers were calculated using the Next Generation Matrix approach [72] (Supplementary Material H).

## Statements

### Data availability statement

Data on dead reported birds is available upon request at https://www.sovon.nl/tellen/telprojecten/dode-vogel-melden. Population trends for the Dutch blackbird population were derived from a national breeding bird monitoring scheme [73]. The datasets on live and dead free-ranging birds analysed in this study are available in the BioStudies database (http://www.ebi.ac.uk/biostudies) under accession codes S-BSST1522, S-BSST1523, S-BSST1867 and can also be accessed via the Pathogens Portal Netherlands: https://www.pathogensportal.nl/arboviruses.html.

### Code availability statement

All code is available in a public GitHub repository at https://github.com/Mariken95/VBD-ABC/.

## Acknowledgements

We would like to thank all volunteers at the bird ringing stations across the Netherlands for their efforts in catching and sampling birds. We would also like to thank all people involved in the coordination of these surveillance schemes as well as those involved in the laboratory analyses. We are grateful to the members of the One Health PACT work package 3 and 4 for their helpful discussions and ideas. Lastly, we are grateful to the organising committees of the Society for Veterinary Epidemiology and Preventive Medicine (SVEPM) conference, Epidemics conference, Modelling in Animal Health (ModAH) conference, and Ecology and Evolution of Infectious Diseases (EEID) conference for the opportunity to present earlier versions of this work.

## Funding

This publication is part of the project ‘Preparing for Vector-Borne Virus Outbreaks in a Changing World: a One Health Approach’ (NWA.1160.18.210), which is (partly) financed by the Dutch Research Council (NWO)). This work used the Dutch national e-infrastructure with the support of the SURF Cooperative using grant no. EINF-9367. The project has received funding from the European Union’s Horizon 2020 research and innovation programme under grant agreement No. 874735 (VEO) NWA.1160.18.210: MdW, MD, EM, LK, NA, JvI. EINF-9367: MdW. Horizon 2020 No. 874735: RS, MK.

## Author contributions

**MdW**: conceptualisation, methodology, software, formal analysis, visualisation, data curation, writing – original draft, writing – reviewing and editing. **GB**: methodology, software, writing - reviewing and editing. **MD**: formal analysis, writing – reviewing and editing. **EM**: data curation, writing - reviewing and editing. **LK**: formal analysis, writing – reviewing and editing. **NA**: data curation. **JvI**: formal analysis, data curation, writing – reviewing and editing. **HvdJ**: conceptualisation, data curation, writing – reviewing and editing. **JvdB**: data collection, data curation, writing – reviewing and editing. **CR**: data collection, data curation, writing – reviewing and editing. **MK**: funding acquisition, conceptualisation, writing – reviewing and editing. **MdJ**: conceptualisation, methodology, supervision, writing – reviewing and editing. **RS**: conceptualisation, data curation, writing – reviewing and editing. **QtB**: conceptualisation, methodology, supervision, writing – original draft, writing – reviewing and editing.

## Competing interests statement

The authors declare no competing interests.

## Supplementary materials

- Supplementary Material A: Model formulation and parameterisation
- Supplementary Material B: Bird dispersal
- Supplementary Material C: Surveillance data
- Supplementary Material D: Inference approach
- Supplementary Material E: Validation of inference approach
- Supplementary Material F: Additional results
- Supplementary Material G: Inference diagnostics
- Supplementary Material H: Reproduction numbers
- Supplementary Material I: Model calibration without PCR datasets

## Supplementary Material A: Model formulation and parameterisation

The equations for the modelled system (Eq 1-23) describe susceptible (Sm), exposed (Em), and infectious (Im) mosquitoes, where the total adult mosquito population is Nm = Sm + Em + Im, and susceptible (S), exposed (E), infectious (I), recovered (R) and dead (D) birds. The bird population is split into two groups: reservoir birds and blackbirds, which are further split into two age classes: those under 1 year old (juvenile blackbirds (*JB)* and juvenile reservoir birds (*JR*)) and those above 1 year of age (adult blackbirds (*AB)* and adult reservoir birds (*AR*)). For each group, the total live bird population in grid cell *i* is N*i* = S*i* + E*i* + I*i* + R*i*.

Mosquito population:

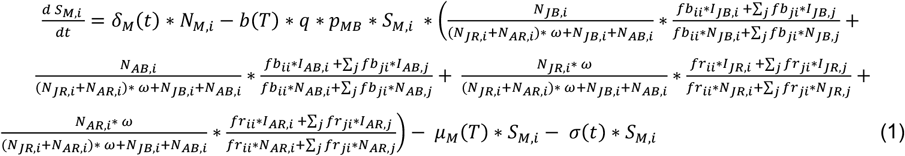

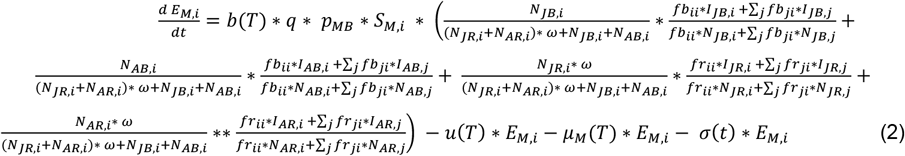

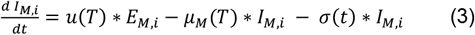

Juvenile blackbird population:

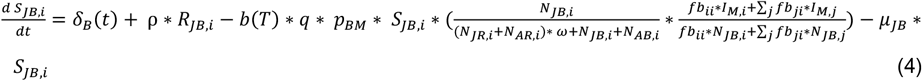

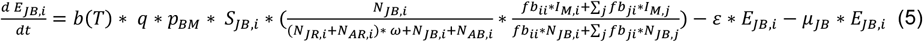

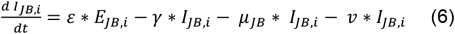

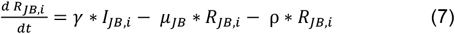

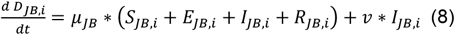

Adult blackbird population:

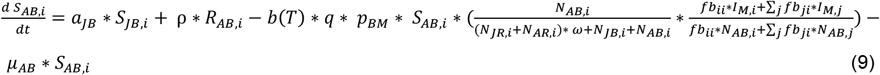

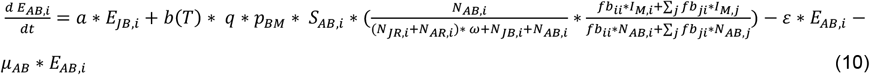

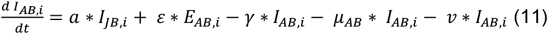

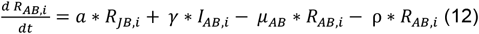

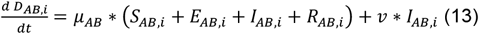

Juvenile reservoir population:

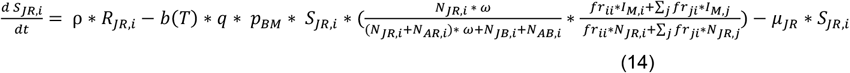

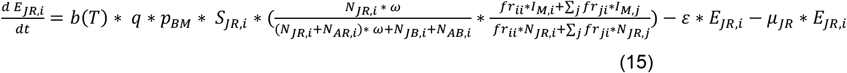

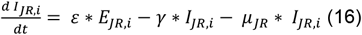

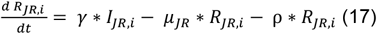

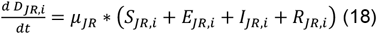

Adult reservoir population:

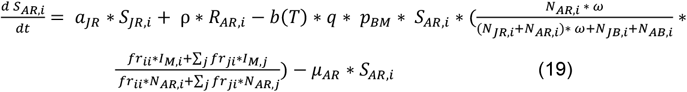

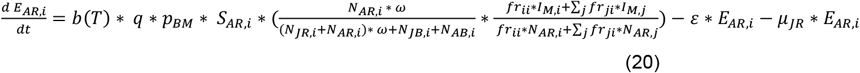

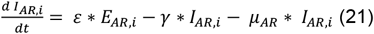

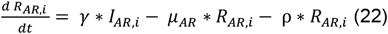

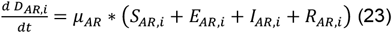

### Model parameters

Parameter values are described in Supplementary Table 1. These can be time-dependent (t) or temperature-dependent (T). Most model parameters were directly obtained from literature, however some were based on calculations.

#### Proportion of bites on competent hosts

As *Culex pipiens/torrentium* mosquitoes bite a wide range of hosts [34], we accounted for the presence of non-competent hosts. For this, we used a parameter that determines the proportion of mosquito bites that are on competent hosts and assumed that the modelled bird populations represent all competent hosts. We also assumed that the total number of (competent and non-competent) hosts is the same in each location. The proportion of bites that are on competent hosts was set equal to the proportion of the total bird population that are competent. This proportion was calculated each day and location, by setting it to 100% in the location where the bird population density was largest. In all other locations this proportion was equivalent to the relative density compared to the cell where this was largest.

#### Bird birth rate

The bird death rate was assumed to be constant over space and time, while births occurred seasonally. These births were distributed uniformly across the period where newborns start leaving their nests assuming a hatching period of 14 days and a further 14 days until they fledge the nest [74]. Ageing from juvenile to adult bird occurred instantaneously on the last day of April, the day before births start.

We used a Leslie matrix to calculate the annual number of births assuming the population remains constant in absence of infection. The following Leslie matrix was developed for the blackbird population, where F stands for the annual fertility rate and S for the annual survival rate of juvenile (j) or adult (a) birds:

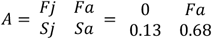

We calculated the fertility rate for adults for which the eigenvalue of the matrix equals 1. The resulting fertility rate was 2.5 births per year, using juvenile survival estimates from [75] and adult survival estimates from [76]. While the blackbird population declined during the first three years of USUV circulation, the population recovered in subsequent years. In the model, we included the possibility of population recovery by assuming a fixed number of annual births (i.e. independent of the population size.) Therefore we calculated the daily number of births during the breeding period based on the population size at the beginning of 2016 (2.5 births / 45 days = 0.056 births per bird per day) and used this fixed value for all years.

Annual survival probabilities were transformed to daily mortality rates to be used in the simulation model based on the following equation: *annual survival probability* = (1 − *daily mortality rate*)*^*365.

The same approach was used for the modelled reservoir population. When the adult lifespan was a parameter to be estimated (in models C, D, S1, S2, S3), the calculations were performed for each simulation run.

#### Diapause

The proportion of active adult mosquitoes (*ϑ*(*t*)) is dependent on daylength *φ*(*t*):

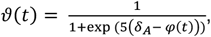

where *δ*_*A*_ is the autumn threshold when the mosquito population starts entering diapause [77]. While estimates of the autumn threshold value are lacking for the Netherlands, this value was estimated to be 13 hours of daylight in the United Kingdom [77]. Using this threshold with local estimates of daylength suggests that that mosquitoes start entering diapause around the beginning of September and all mosquitoes are in diapause towards the end of September. Diapause induction is not only dependent on day length, but also on temperature [78]. To adjust for somewhat warmer autumn temperatures in the Netherlands compared to the United Kingdom [79,80] and to match observations from local mosquito trapping data, we assumed that the average time to entering diapause (since 1^st^ September) was 20 days, leading to an estimated rate of 0.05 at which mosquitoes enter diapause. In our model, mosquitoes entered diapause in the period between September to mid-October at this fixed daily rate. Any remaining active mosquitoes at the end of this period were forced to enter diapause in mid-October. Mosquitoes re-emerged from diapause on the first of April, following estimates from the abundance model.

**Supplementary Table 1:** Parameters and their values used in the model. T indicates temperature, t indicates time.

| Parameter | Description | Value | Source |
| --- | --- | --- | --- |
| **Blackbirds** |  |  |  |
| $\delta_b(t)$ | Daily number of births | 1 May – 15 June:<br>0.056 per bird | Value: calculated, see above. |
|  |  | Else: 0 | Timing: [75] |
| $\mu_{JB}$ | Mortality rate, juvenile | 0.0059 / day | [75] |
| $\mu_{AB}$ | Mortality rate, adult | 0.0011 / day | [76] |
| $p_{bm}$ | Transmission probability, mosquito to blackbird | 0.88 | [57] |
| $\varepsilon$ | Intrinsic incubation rate | 0.67 / day | [49,82] |
| $\gamma$ | Recovery rate | 0.25 / day | [83] |
| $v$ | Infection mortality rate | NA | To be estimated |
| $\rho$ | Rate of immunity loss | 0.0014 / day | [84] |
| $fb_{ji}$ | Fraction of blackbirds in cell $j$ dispersing to $i$ | Variable | Calculated, see Supplementary Material B |
| $\alpha_{JB}$ | Ageing rate from juvenile to adult | On 30<sup>th</sup> April: 1.<br>Else: 0 | [75] |
| **Additional reservoir population** |  |  |  |
| $\delta_r(t)$ | Daily number of births | 1 May – 15 June: To be estimated<br>Else: 0 | Value: to be estimated. See above.<br>Timing: [75] |
| $\mu_{JR}$ | Mortality rate, juvenile | 0.0059 / day | Set equal to blackbird juvenile |
| $\mu_{AR}$ | Mortality rate, adult | NA | To be estimated |
| $p_{bm}$ | Transmission probability, mosquito to reservoir host | 0.88 | [57] |
| $\varepsilon$ | Intrinsic incubation rate | 0.67 / day | [49,82] |
| $\gamma$ | Recovery rate | 0.25 / day | [83] |
| $\rho$ | Rate of immunity loss | 0.0014 / day | [84] |
| $fr_{ji}$ | Fraction of reservoir birds in cell $j$ dispersing to $i$ | Variable | Calculated, see Supplementary Material B |
| $\alpha_{JR}$ | Ageing rate from juvenile to adult | On 30<sup>th</sup> April: 1.<br>Else: 0 | Set equal to blackbird |
| $\omega$ | Relative biting frequency | NA | To be estimated |
| ***Culex pipiens*** |  |  |  |
| $\delta_m(t)$ | Emergence rate | Variable | Estimated from abundance data, see above |
| $\mu_m(T)$ | Mortality rate [daily] | For $T < 33$ °C:<br>$1 / (-4.86 \cdot T + 169.8)$,<br>else 0.17 | [85] fitted to data from [86–88] |
| $b(T)$ | Biting rate [daily] | $0.344 / (1 + 1.231 \cdot \exp(-0.184 \cdot (T - 20)))$ | [82] fitted to data from [89] |
| $q$ | Proportion of bites that are on competent host | Variable | Calculated, see above |
| $p_{mb}$ | Transmission probability, blackbird & reservoir population to mosquito | NA | To be estimated |
| $u(T)$ | Extrinsic incubation rate [daily] | For $T > 13$ °C:<br>$7.38 \cdot 10^{-5} \cdot T \cdot (T - 11.4)(45.2 - T)^{1/2}$,<br>else 0 | [85] fitted to data from [90,91] |
| $\sigma(t)$ | Rate of entering diapause | 1 Sept – 15 Oct: 0.05.<br>Else: 0 | [92] |

#### Virus re-emergence

We allowed for infectious mosquitoes to re-emerge from diapause thereby carrying the infection forward to the next transmission season as well as for infections to be re-introduced from migratory birds. This re-emergence was included in the model through reseeding infections in mosquitoes on the second of April each year (the day after mosquitoes re-emerged from diapause). This was implemented as an infection rate moving mosquitoes from the susceptible to exposed state. This rate was determined for each location separately by multiplying the prevalence in mosquitoes at the end of the previous season in each location (in mid-September) by the re-emergence rate parameter. The value of this parameter was estimated during model calibration and was the same across all locations and years. Re-emergence of USUV in spring represents the possibility of local overwintering and/or re-introductions from migratory birds.

#### Host-vector abundance scaling parameter

Bird and mosquito abundance estimates were on a relative scale, because the total population size is not known for these populations. We therefore introduced a host-vector abundance scaling parameter, which converts the host and vector abundance to be on the same scale, meaning that their ratio reflects the vector-to-host ratio. This parameter was used to determine the initial conditions of the model and was therefore not included in the differential equations.

#### Mosquito emergence and death rate

Mosquito abundance estimates were used to estimate vital dynamics in the mosquito population. Mosquito death rate was assumed temperature-dependent (Supplementary Table 1) and therefore differed over space and time. Similar to Perkins et al. [81], the number of emerging mosquitoes was estimated for each location and day such that, together with the mortality rates, the resulting population size follows estimates from the abundance model.

### Model design and implementation

The model was implemented using the SimInf package [93]. The model consisted of both stochastic and deterministic processes. The majority of processes (including transitions between compartments and daily dispersal) are modelled as continuous-time discrete variable Markov chains using the direct Gillespie stochastic simulation algorithm. Bird birth events were processed deterministically (and are therefore not shown in the model equations) to ensure an equal number of births each year, independent of changes in the population size due to virus transmission. This allowed for the recovery of the host population size after increased death rates due to infection. Mosquito emergence and death rates were also processed deterministically, to ensure the mosquito population sizes always follow the predicted abundance (see ‘Input layers’). Seasonal movements of birds were also included deterministically. These deterministic processes are executed at the end of each time step for which they were planned.

### Overview of data inputs

Information on blackbird abundance, *Cx. pipiens* abundance and temperature was obtained from literature, see main text. Supplementary Figure 1 provides an overview of these input data.

**Supplementary Figure 1:**
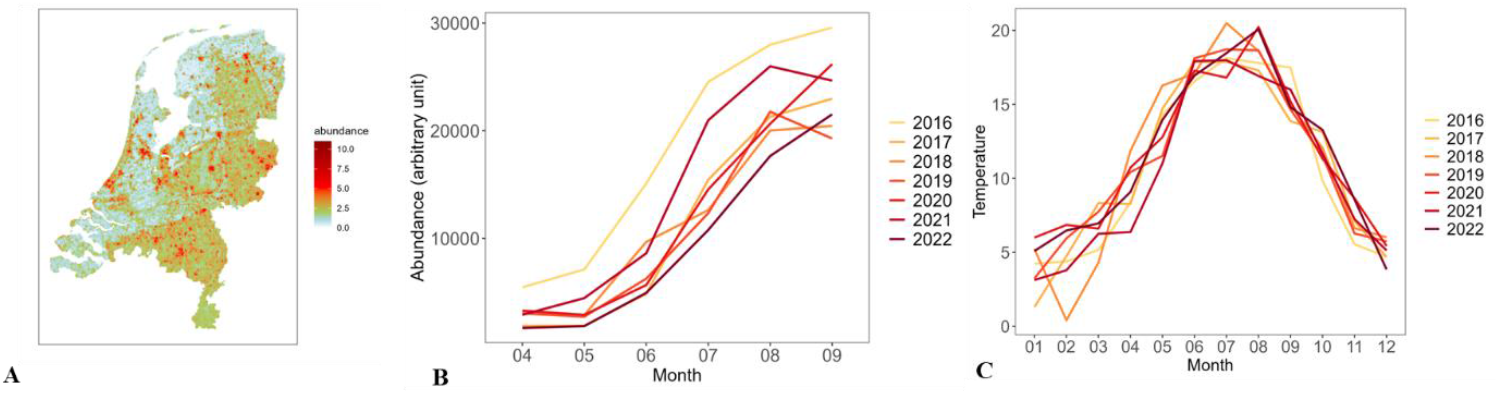
A. Relative blackbird abundance map. B. Monthly estimates of the relative abundance of Culex pipiens mosquitoes in the Netherlands for each year. C. Mean monthly temperature during 2016-2022.

Blackbird and *Cx pipiens* abundance estimates were obtained from previously published studies [25,26]. Both studies used a machine learning approach (Random Forest models), which is an important topic in current abundance modelling research [94]. A stacked machine learning model, that combines estimates from multiple models, predicting *Ae albopictus* egg populations showed that the Random Forest model was the most important model [94]. This study also identified lagged temperature as an important predictor, similar to the model used here.

#### Blackbird abundance

Estimates of blackbird abundance were obtained from Dellar et al [26]. Random Forest models were fitted to blackbird point count data (n=183,123 observations) from the Netherlands (Meetnet Urbane Soorten [63], Meetnet Agrarische Soorten [64] and Common Bird Census [65] from Sovon Vogelonderzoek Nederland (Dutch Centre for Field Ornithology)) and France (Common Bird Monitoring Scheme [66]) in the period 2001-2016, using a set of environmental and climatic predictors. Several measures were calculated to assess model performance (Supplementary table 2), showing a good overall performance. Land use-related variables were the most important predictor variables.

**Supplementary table 2:** Model performance statistics blackbird abundance model. Adapted from Dellar et al [26]. Results were scaled to have a maximum of 1.

| Mean absolute error | Mean Forecast Error | Root Mean | Correlation predicted vs | Intercept | Slope | Explained Variance |
| --- | --- | --- | --- | --- | --- | --- |
|  |  | **Squared Error** | **observed abundances** |  |  |  |
| 0.325 | -0.012 | 0.544 | 0.954 | -0.231 | 1.127 | 69.7% |

#### Mosquito abundance

Estimates of *Cx pipiens* abundance were obtained from Krol et al [25]. Data on female mosquito trap counts (n=1,447 observations) was obtained from the National Mosquito Survey 2010-2013 [67] and the MODIRISK project [68]. Random Forest models were fitted to female mosquito count data including a set of land cover, environmental, and climatic predictors. Several measures were calculated to assess model performance (Supplementary table 3), showing a good overall performance. The importance of climate-related variables was similar to land use/cover-related variables, see [25].

**Supplementary table 3:** Model performance statistics Cx pipiens abundance model. Adapted from Krol et al [25]. Results were scaled to have a maximum of 1.

| **Mean absolute error** | **Mean Forecast Error** | **Root Mean Squared Error** | **Correlation predicted vs observed abundances** | **R<sup>2</sup>** | **Intercept** | **Slope** | **Explained Variance** |
| --- | --- | --- | --- | --- | --- | --- | --- |
| 0.399 | -0.005 | 0.544 | 0.922 | 0.819 | -0.394 | 1.252 | 27.9% |

## Supplementary Material B: Bird dispersal

The Vogeltrekstation NIOO-KNAW dataset containing Dutch ringing recovery information was used to estimate bird movement distances. We created a dataset (n=333) using the following criteria:

- Blackbirds
- Observations from 1970 onwards
- Two observations per bird available: ringing and dead recovery (at least one day later)
- Ringed and recovered within the Netherlands
- High accuracy in location, age and date information
- Ringed in breeding season (April, May, June) or post-breeding season (July, August, September)
- Juvenile or adult stage (nestlings were excluded, because they cannot move)

The final dataset included observations from all provinces, although most observations came from the North and Middle regions (Supplementary figure 2).

Two movement patterns were distinguished in the model, representing daily movements and seasonal dispersal. Daily movements include foraging-type movements often within the bird’s home range, i.e. the area the bird regularly traverses around in its territory. In addition to daily movements, we included seasonal dispersal that represents the search for a new nest location. Seasonal dispersal takes place during the change from breeding to post-breeding season in early July.

To parameterize these processes, we compared distance between ringing and recovery location within seasons (daily movements) and between breeding and subsequent seasons (seasonal dispersal). Only bird movement during the transmission season was included, which was divided into two periods: breeding season (April to July) and post-breeding season (July to October). The bird population was divided into two age groups: juvenile (from fledging until the next breeding season) and adult (birds older than one year).

**Supplementary figure 2:**
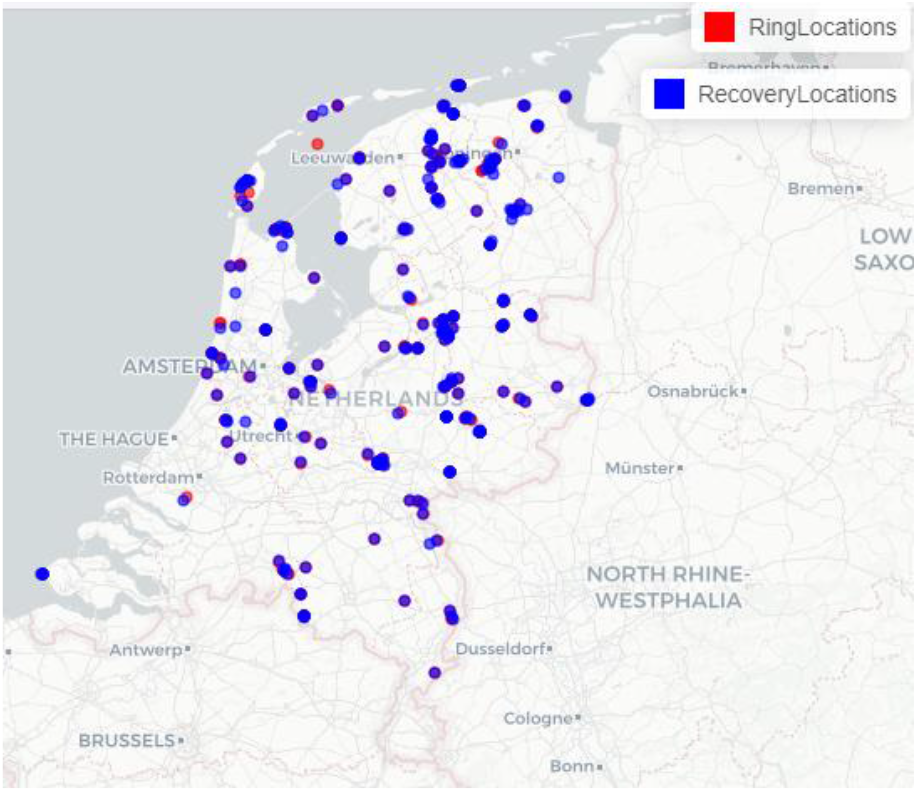
Locations of ringed and recovered birds. Image created using ‘mapview’ in R.

### Daily movements

We found no evidence of an association between distance and days since ringing (p=0.30) (Supplementary figure 3). For blackbirds that were ringed and recovered in the same season of the same year, we therefore assumed that the distance between ringing and recovery location represents the daily movement distance around the home base.

No difference was found in daily movement distance between juvenile and adult blackbirds (Wilcoxon rank sum test breeding season: p=0.51 & post-breeding season: p=0.52). However, there was evidence for a difference in movement distances between breeding and post-breeding season (Wilcoxon rank sum test p=0.03, Supplementary figure 4). We therefore estimated daily movement for each season, combining both age groups.

**Supplementary figure 3:**
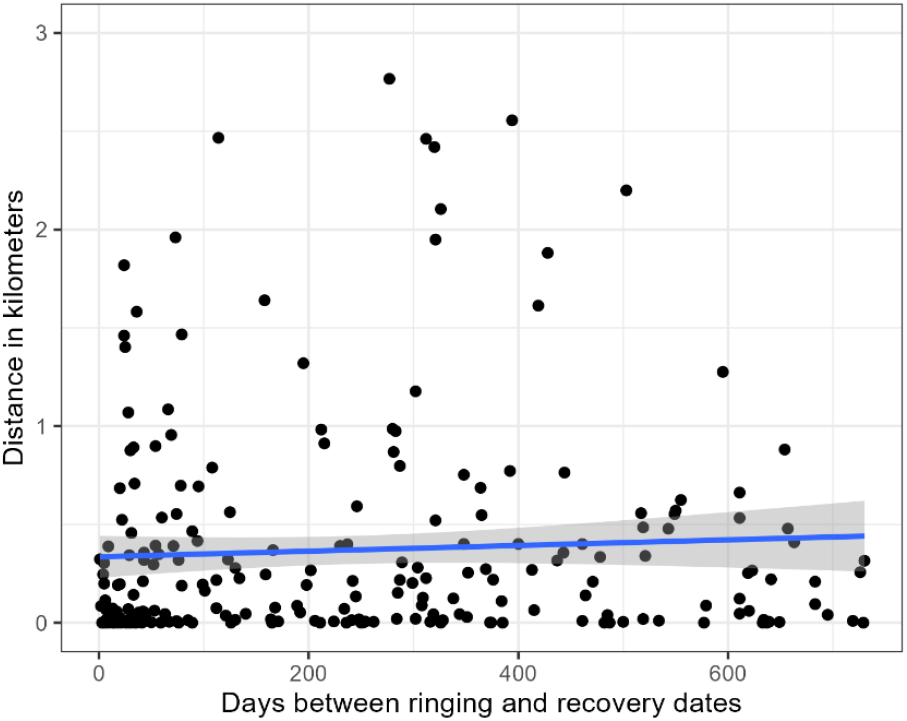
Relationship between distance and days since ringing.

**Supplementary figure 4:**
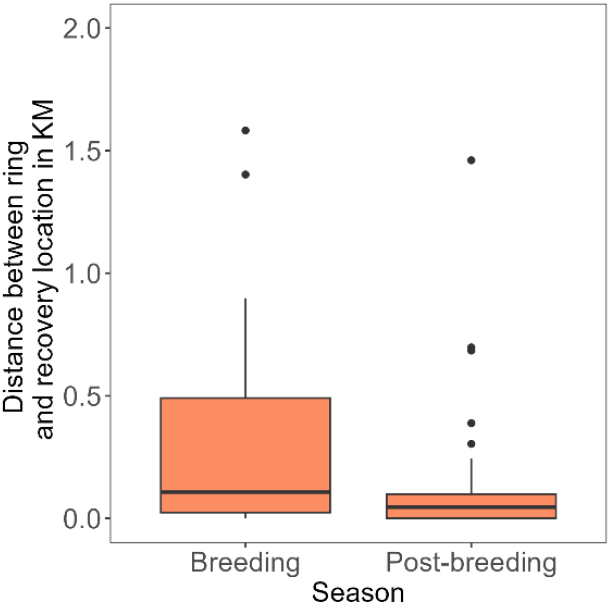
Distance between ring and recovery location of blackbirds by season. Only blackbirds ringed and recovered within the same season were included.

We fitted Weibull, gamma, and exponential distributions to each dataset and compared fits based on AIC (Supplementary table 4). The fitting of an exponential distribution was not possible for breeding season due to extreme outliers. As the Weibull distribution was the best fitting distribution in both cases, we used parameters from this distribution to calculate daily movement matrices. This resulted in a median daily movement distance of 143 m (inter-quartile range 19 - 692) during breeding season and 29 m (inter-quartile range 4 – 129) during post-breeding season.

**Supplementary table 4:** AIC values of three different distributions fitted to within-season dispersal distance and parameter estimates of the fitted Weibull distributions.

|  | Weibull | Gamma | Exponential | Weibull parameters |  |
| --- | --- | --- | --- | --- | --- |
|  |  |  |  | Scale | Shape |
| **Breeding (n=23)** | 324 | 330 | NA | 329.4 | 0.44 |
| **Post-breeding (n=31)** | 331 | 333 | 375 | 63.4 | 0.46 |

### Seasonal dispersal

The same approach was used to quantify between-season dispersal. To estimate the distance blackbirds move during seasonal dispersal, we selected birds that were ringed in breeding season and recovered after this season. There was evidence for a difference in seasonal dispersal distances between adult and juvenile birds (Wilcoxon rank sum test p=0.04), see Supplementary figure 5. In the transmission simulation model, seasonal dispersal took place once a year on the change from breeding to post-breeding season (1^st^ of July).

**Supplementary figure 5:**
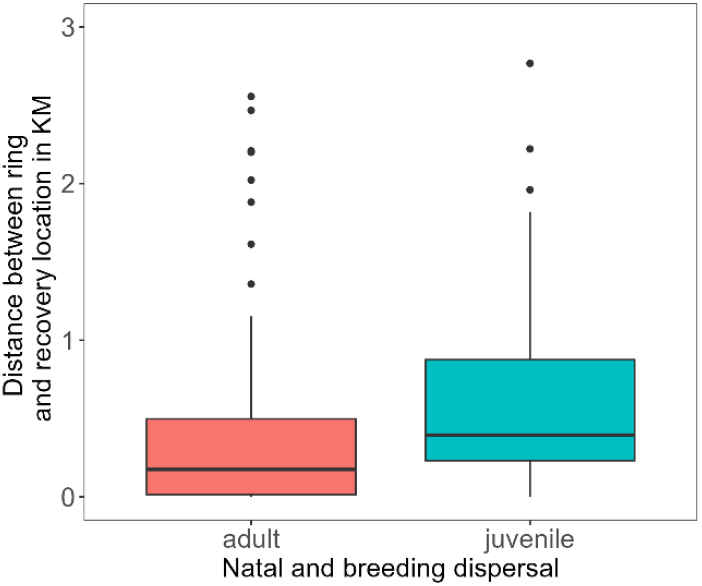
Distance between ringing and recovery location of blackbirds by age for birds ringed during breeding season and recovered after this season.

Again, we fitted Weibull, gamma, and exponential distributions to each dataset and compared fits based on AIC (Supplementary table 5). The fitting of an exponential distribution was not possible for both datasets due to extreme outliers. As the Weibull distribution was the best fitting distribution on average, we used parameters from this distribution (Supplementary table 5) to calculate seasonal dispersal matrices. This resulted in a median seasonal dispersal distance of 377 m (inter-quartile range 95 - 1114) for juveniles and 180 m (inter-quartile range 19 – 1062) for adults.

**Supplementary table 5:** AIC values of three different distributions fitted to between-season dispersal distance.

|  | Weibull | Gamma | Exponential | Weibull parameters |  |
| --- | --- | --- | --- | --- | --- |
|  |  |  |  | Scale | Shape |
| **Breeding season to post-breeding season** |  |  |  |  |  |
| **Juvenile (n=40)** | 612 | 610 | NA | 668.2 | 0.64 |
| **Adult (n=77)** | 1123 | 1149 | NA | 459.4 | 0.39 |

### Dispersal matrices

To link the dispersal kernels to the transmission simulation model, we estimated the probability to disperse to any location from the origin location, using an individual-based simulation analysis. We generated 1,000,000 random points within a grid cell, representing the origin location of birds. The grid size corresponded to the grid size used for the transmission simulation model. The distance that each bird moved was sampled from the dispersal kernel. The direction of the movement was generated from a uniform distribution. The destination grid cell was recorded for each bird. The proportion of birds that ended up in each grid cell was calculated to construct a movement matrix, see Supplementary figure 6 for a visualisation of the daily dispersal matrices. This matrix was applied to each location each day, where movements outside of the country borders were not allowed.

**Supplementary figure 6:**
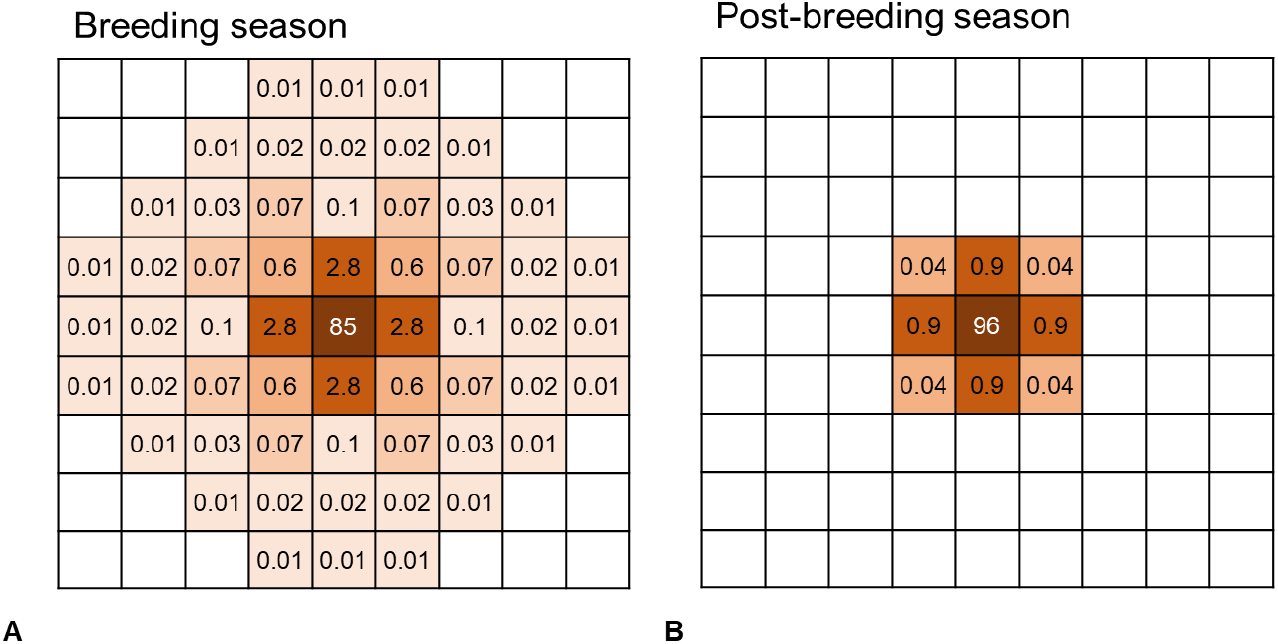
Visualisation of daily movement matrices for a) breeding season, b) post-breeding season. Values indicate the proportion of birds moving to that respective cell using the middle cell as origin location. Values lower than 0.01% are not shown.

### Daily movement scenarios for extra reservoir host

To explore the movement behaviour of a potential additional reservoir host, we created two movement scenarios: increased movement and movement within uncertainty range (sensitivity analysis S3).

#### Increased movement scenario

As this reservoir population represents several species, the ‘increased movement’ scenario was created by mixing dispersal kernels of two bird species. We obtained estimates of movement distances for mallards from literature, because these have been found positive for USUV infection in the Netherlands and show movement behaviour distinct from blackbirds. Median movement distance for mallards outside the breeding season was estimated at 1.5 km [95]. Using roughly the same ratio between breeding and post-breeding movement as for blackbirds (daily movement in breeding season being around 3x further than in post-breeding season), we assumed daily movement in breeding season to be 4.5 km. We calculated the corresponding scale parameters (keeping the shape parameter constant) that resulted in a median movement distance of 4.5 km in breeding season and 1.5 km in post-breeding season (Supplementary table 6). These parameters were then combined with the blackbird movement parameters to create the ‘increased movement’ scenario: reservoir host movement behaviour consists of 80% blackbird and 20% mallard dispersal.

#### Movement within uncertainty range (sensitivity analysis)

We found blackbird movement distances in post-breeding season to be lower than in breeding season (Supplementary figure 4). To explore the implications of this finding we created a sensitivity analysis where movement in both seasons was parameterised as breeding season movement. This was used for both the blackbird and the reservoir population.

Resulting dispersal kernels are shown in Supplementary figure 7 and corresponding parameters are shown in Supplementary table 6.

**Supplementary figure 7:**
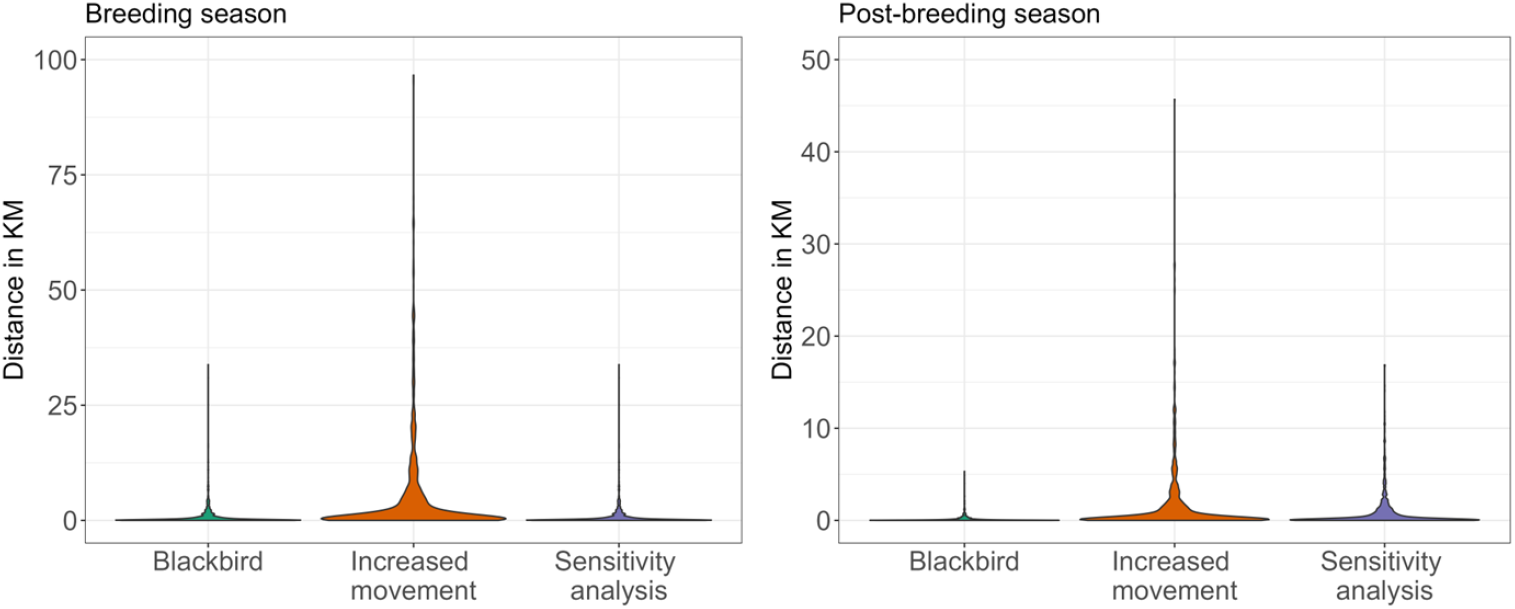
Daily movement kernels for blackbirds and the two scenarios.

**Supplementary table 6:**
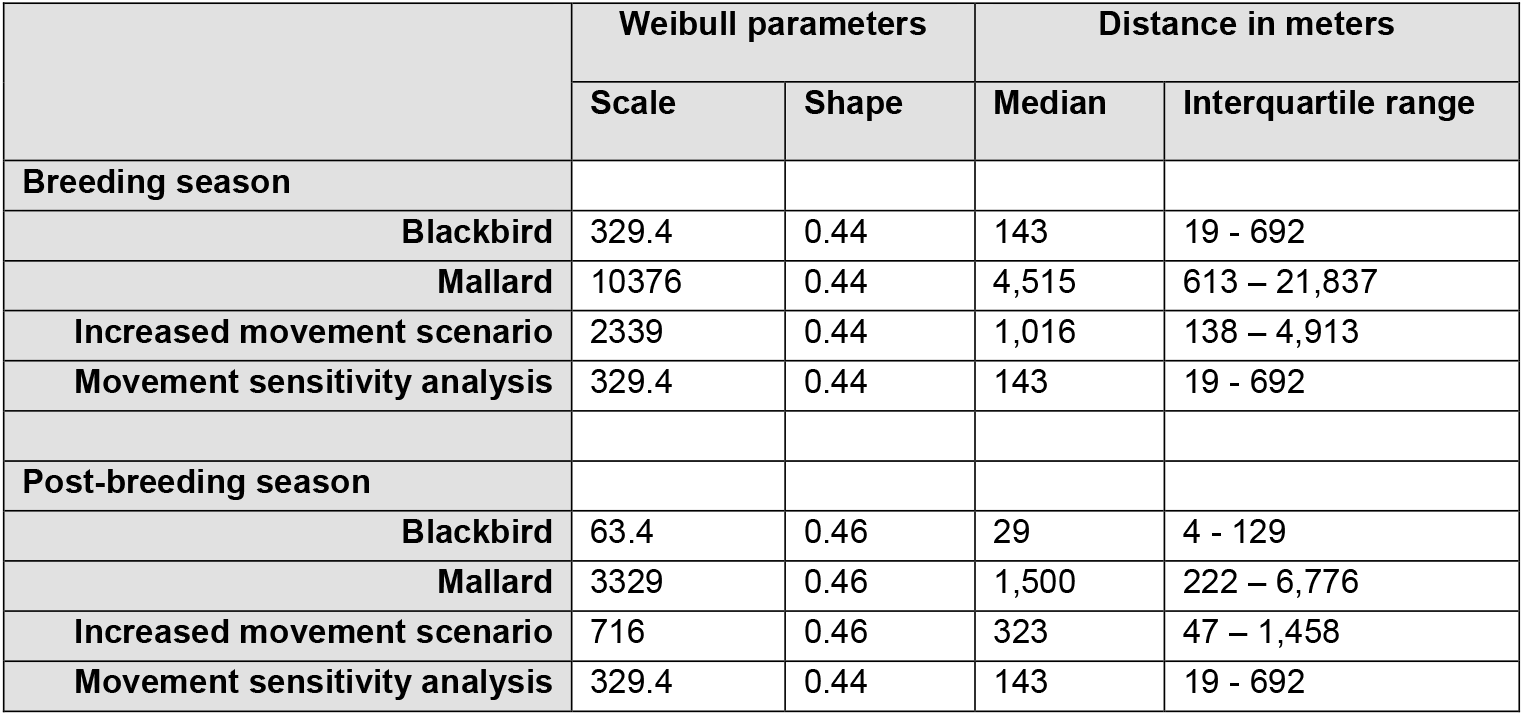
Daily movement parameters for blackbirds, mallards and the three mixture scenarios applied to the additional reservoir population.

|  | Weibull parameters |  | Distance in meters |  |
| --- | --- | --- | --- | --- |
|  | Scale | Shape | Median | Interquartile range |
| **Breeding season** |  |  |  |  |
| **Blackbird** | 329.4 | 0.44 | 143 | 19 - 692 |
| **Mallard** | 10376 | 0.44 | 4,515 | 613 – 21,837 |
| **Increased movement scenario** | 2339 | 0.44 | 1,016 | 138 – 4,913 |
| **Movement sensitivity analysis** | 329.4 | 0.44 | 143 | 19 - 692 |
| **Post-breeding season** |  |  |  |  |
| **Blackbird** | 63.4 | 0.46 | 29 | 4 - 129 |
| **Mallard** | 3329 | 0.46 | 1,500 | 222 – 6,776 |
| **Increased movement scenario** | 716 | 0.46 | 323 | 47 – 1,458 |
| **Movement sensitivity analysis** | 329.4 | 0.44 | 143 | 19 - 692 |

## Supplementary Material C: Surveillance data

A more detailed description of the surveillance data can be found in Münger et al. [20].

### PCR and serology testing in live birds (datasets 1 & 2)

Live free-ranging birds were sampled by trained volunteer ornithologists at different locations throughout the Netherlands between March 2016 and December 2022. Sampling was performed under ethical permits AVD801002015342 and AVD80100202114410 issued to NIOO-KNAW. Birds were captured using mist nets and ringed, after which a throat swab, a cloacal swab, and a blood sample were collected. Viral diagnostics was done by RT-PCR on throat and cloacal swabs. Bird sera were screened for USUV and WNV antibodies in a protein micro-array, where positive samples were subsequently tested for viral neutralization using Virus Neutralization Tests (VNT) and Focus Reduction Neutralization Tests (FRNT) as previously described [96,97].

### Reporting and PCR testing of dead birds (datasets 3 & 4)

Dead free-ranging birds were reported through a citizen science-based alerting system run jointly by Sovon and the Dutch Wildlife Health Centre (DWHC) between March 2016 and December 2022 [98]. A selection of the reported dead birds, based on the freshness of the carcasses (death within ca. 24h) and the likelihood of disease being the cause of death was collected, necropsied, and samples were taken. From 2016 to 2018, dead blackbirds were tested for USUV by RT-PCR if necropsy indicated USUV as a possible cause of death; from 2019, all submitted birds were tested. In 2016, dead blackbirds submitted for post-mortem examination were selectively chosen by the DWHC to determine the expansion of the USUV outbreak in 2016. So, the analysed dead birds were mainly selected from areas where USUV had not been detected previously but within several kilometres from where it had been. From 2017 onwards, the selection of fresh dead birds for post-mortem examination was random over the Netherlands.

### Blackbird population trends (dataset 5)

Population trends for the Dutch blackbird population were derived from a national breeding bird monitoring scheme [73] and further analysed in [21]. This scheme consists of standardised counts in thousands of sample sites (plots) by voluntary observers. Each year, multiple visits (5-10) within the breeding season (March-July) were used to derive the number of territories. These territories were determined based on species-specific interpretation criteria, including behaviour observed, detection probabilities, and timing of observation.

## Supplementary Material D: Inference approach

### The ABC-SMC algorithm

The ABC-SMC algorithm used here enhances the basic ABC algorithm by incorporating two main steps: weighted resampling of simulated particles and a gradual reduction in tolerance. Our specific implementation of the algorithm improves upon [99] original algorithm in three ways: (1) an adaptive threshold schedule selection based on quantiles of distances between simulated and observed data [100,101] (2) an adaptive perturbation kernel width during the sampling step, dependent on the previous intermediate posterior distribution [71,102], and (3) the capability to use multiple criteria simultaneously. The algorithm proceeds as follows:

#### Step 1: Definition of summary statistics, distances, and model

- **Summary statistics:** Define the summary statistics *S* that adequately capture the essential features of the observed data. We used five summary statistics (Supplementary figure 9): (i) seroprevalence in live blackbirds for each month and region (North, Middle, South), (ii) PCR prevalence in live blackbirds for each month and region, (iii) PCR prevalence in dead blackbirds for each month and region, (iv) annual relative blackbird population size for each region and (v) the relative number of reported dead blackbirds per year within each region. Regions were created by dividing the country into three equally sized regions (Supplementary figure 8).
- **Distances:** Define distance functions *D*_1_,*D*_2_,…,*D*_5_. For the three first summary statistics the negative log-likelihood was used as distance measure:

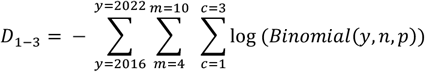

where y is the number of positive samples in the data, n is the total number of samples in the data, and p is the (sero)prevalence in the model simulation for seroprevalence in live blackbirds (D1), PCR prevalence in live blackbirds (D2), and PCR prevalence in dead blackbirds (D3). Likelihood values were summed across years (y, 2016 to 2022), months (m, April to October), and regions (c, North, Middle, and South). For the last two summary statistics, distance was calculated as the sum of squared differences between observed and simulated datapoints for each year (y, 2016 to 2022) and region (c, North, Middle, and South):

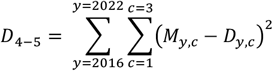

where *Dy,c* and *My,c* are the relative number of dead blackbirds (D4) or the relative blackbird population size (D5) per year and region in the data and model respectively.
- **Model definition:** Define the model *M*(*θ*) which takes a parameter vector *θ* and generates simulated data from which summary statistics are calculated. Each model replicate was simulated 3 times using the same parameter set to reduce the impact of stochasticity. Distances were calculated for each simulation, but the acceptance decision was based on the average of these three distances.

#### Step 2: Particle generation

- **First iteration** (*t* = 1):
  – Generate *N* particles *θ*_*i*_ by drawing *θ*_*i*_ ∼ *π*(*θ*), where *π*(*θ*) is the prior distribution.
- **Subsequent iterations** (*t >* 1):
  – Draw *θ*_*i*_ from the set of particles accepted at iteration *t* − 1 with probability proportional to their weights *w*^(*t*−1)^.
  – **Particle perturbation:** New particles were generated by perturbing each parameter *θ*_*i,k*_ (where *k* indexes the parameters of *θ*) using a Gaussian kernel with a standard deviation 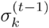 of 0.5 times the empirical standard deviation of the accepted particles for that parameter in the previous iteration:

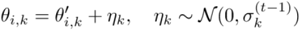

where 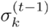 is the standard deviation of parameter *k* in the accepted particles at iteration *t* − 1.
  – **Check perturbed particle:** If any perturbed parameter *θ*_*i,k*_ is outside the prior distribution *π*(*θ*_*i,k*_), reject the perturbed particle and repeat the particle generation step until *θ*_*i,k*_ lies within *π*(*θ*).

#### Step 3: Running the model and calculating summary statistics and distances

For each particle *θ*_*i*_:

– Run the model *M*(*θ*_*i*_) to obtain simulated data.
– Calculate summary statistics *S*_model_(*θ*_*i*_) for the simulated data.
– Compute the distances *D*_1_,*D*_2_,…,*D*_5_ between the summary statistics *S*_model_(*θ*_*i*_) and the summary statistics *S*_obs_ of the observed data:

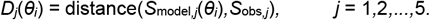

#### Step 4: Particle acceptance

- **First iteration** (*t* = 1): Accept all *N* particles as the initial sample from the prior distribution.
- **Subsequent iterations** (*t >* 1): Accept particle *θ*_*i*_ if and only 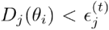 if for all distances *j* = 1,2,…,5. This means that a particle was only accepted when the distances of all summary statistics were below their respective threshold values.

#### >Step 5: Calculation of associated weight

- **First iteration** (*t* = 1): Set weight for each accepted particle *θ*_*i*_ as *w*_*i*_ = 1.
- **Subsequent Iterations** (*t >* 1): Compute the weight for each accepted particle *θ*_*i*_ as:

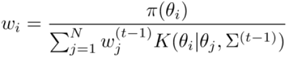

where *K*(*θ*_*i*_|*θ*_*j*_,Σ^(*t*−1)^) is the perturbation kernel centered at *θ*_*j*_

#### Step 6: Repeat steps 2 to 5 until *N* particles are accepted

#### Step 7: Normalise the weights

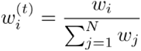

#### Step 8: Calculation of new thresholds from the 75th percentile values of *D*_1_,*D*_2_,…,*D*_5_

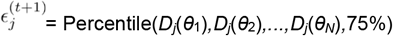

#### Step 9: Increment the iteration number

- Set *t* = *t* + 1.
- Go back to step 2 and repeat the process until convergence criteria are met.

The algorithm was run until the acceptance rate (number of particles accepted over the total number of particles tested during an iteration) reached 5% or lower. Convergence of the posterior distributions was confirmed by visual inspection of intermediate distributions of consecutive generations and by visual inspection of the threshold value of consecutive generations. Minor reductions in threshold values across generations suggests the algorithm has converged.

The algorithm was run using 3 independent chains, where each chain generated 200 accepted parameter sets for each generation. The last posterior distribution therefore consisted of 600 accepted particles.

### Design of summary statistics

The observation process, when known, was accounted for in three ways. When no data were available for a specific month-region combination, the distance for that data point was not calculated. Secondly, in the model, prevalence in dead birds was calculated among birds that died the previous day to reflect the data collection process where only recently-died birds were submitted for testing. Thirdly, model-predicted population size was calculated by averaging the period April to July, similar to when counts were conducted to estimate population size [73]. Distances between the observed and simulated summary statistics were calculated for each summary statistic separately, with each summary statistic having its own threshold value for acceptance. For the three (sero)prevalence-based summary statistics the likelihood of the observed prevalence was used as distance measure. For the summary statistics related to population size and dead blackbird count, the distance was calculated as the sum of squared differences between observed and simulated datapoints.

**Supplementary figure 8:**
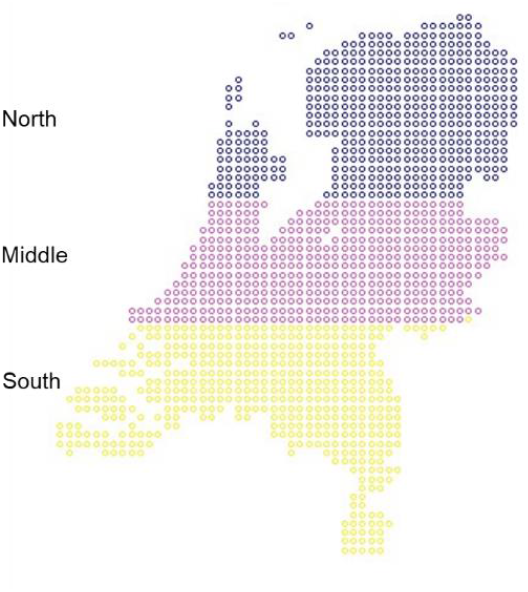
Division of the Netherlands into three equally sized regions.

**Supplementary figure 9:**
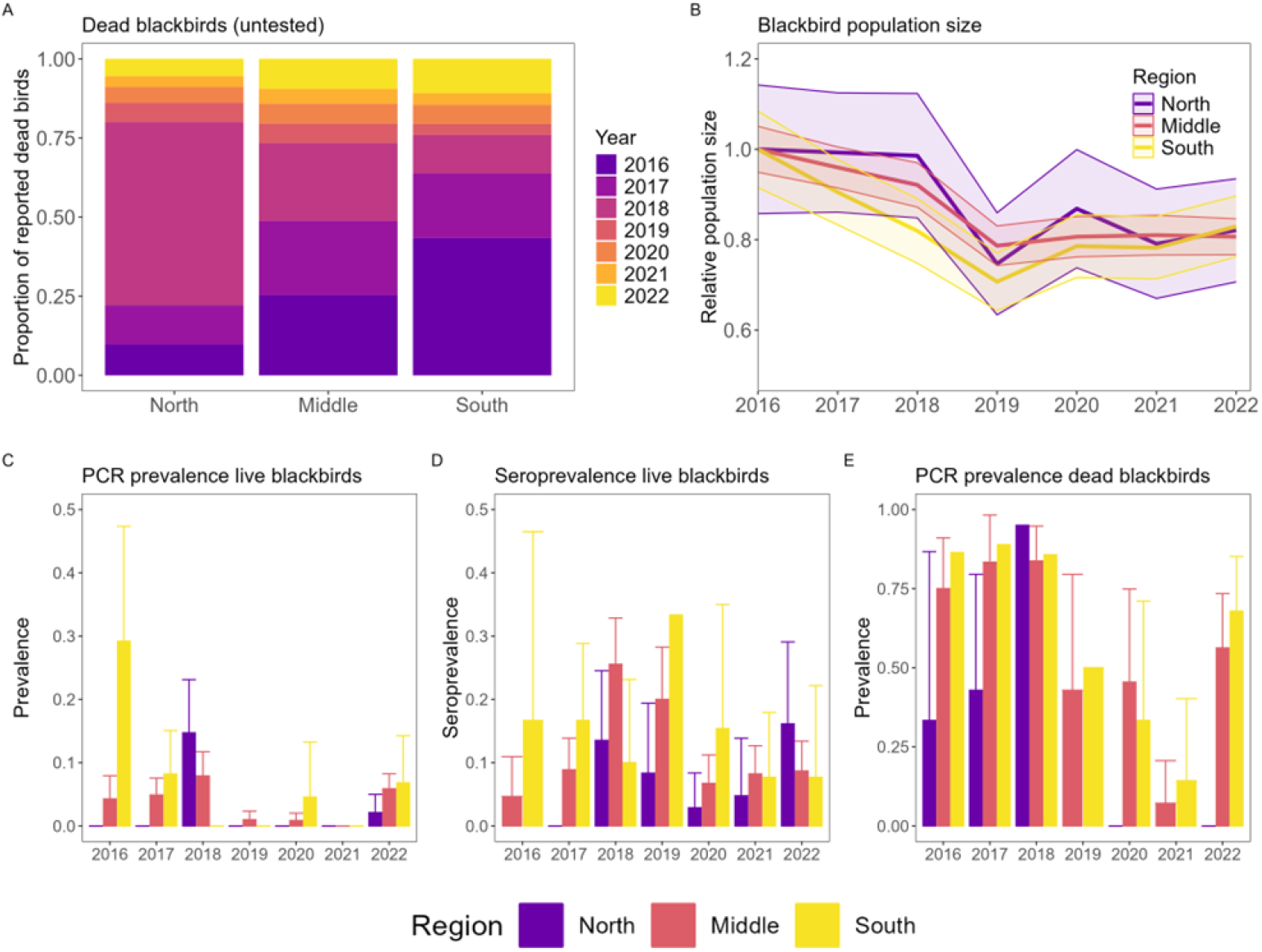
Overview of surveillance data by year and region. One summary statistic was constructed for each of the datasets. Data are presented as mean (solid line in B, bars in C-E) with 95% confidence intervals (shaded area in B, error bars in C-E).

### Prior distributions

All prior distributions were uniformly distributed. We discuss each parameter’s definition, chosen prior and reasoning behind our choices.

#### Re-emergence rate

Definition Fraction of the prevalence in mosquitoes in September that persists to the subsequent spring

Prior 0 to 0.3

Justification It is largely unknown which overwintering processes play a role in USUV re-emergence. One possible route is through vertical transmission in diapausing mosquitoes. Estimates of the efficiency of this route are lacking for USUV in *Cx. pipiens* mosquitoes. For other viruses and mosquitoes estimates are available such as for dengue where studies found vertical infection rates were often very low, but could get up to 13% [103]. Due to a lack of estimates for *Cx pipiens* and USUV and to allow for alternative routes such as maintenance in birds or reintroductions, we assumed a maximum re-emergence rate of 0.3.

#### Transmission probability bird to mosquito

Definition Probability that a mosquito gets infected upon a bite on an infectious bird

Prior 0 to 1

Justification Variation in values used in other models and in experimental studies is large, e.g. a value of 12.5% was used in Rubel et al.’s USUV model [82], while Turell et al. found 81% of mosquitoes infected after oral exposure to WNV [104]. We therefore did not restrict the range of possible values and allowed for values between 0 and 1.

#### Abundance scaling parameter

Definition Multiplication factor for host population size. This converts the host and vector abundance to be on the same scale, such that their ratio reflects the vector-to-host ratio

Prior 0.03 to 2

Justification This includes a wide range of vector-to-host ratios, ranging from around 1043:1 to 7:1 measured on August 15^th^ (the period where mosquito size peaks). Exact estimates vary depending on other parameters as the extent of transmission and the infection mortality rate affect the bird population size. The lower bound of 0.03 ensures that the mean starting abundance is at least 26 blackbirds per grid cell, as otherwise the population size becomes unreasonably and impractically small.

#### Infection mortality rate in birds

Definition Rate at which infectious birds die from infection Prior 0.1 to 1

Justification Significant mortality has been observed and reported following USUV infection, with first estimates of about 30% of infected blackbirds dying [82]. We therefore set the prior distribution between 1 and 10 days until death.

#### Historical annual force of infection (FOI)

Definition Rate at which susceptible hosts became infected per year prior to 2016. This determines the level of immunity present in birds at the start of 2016.

Prior 0 to 0.3

Justification There is no evidence of PCR-positive birds before 2016 in the Netherlands. The lower bound of the prior distribution is therefore set to 0. The seroprevalence in autumn 2016 was estimated at 4.7% (95%CI 0-11%) in the middle region and at 17% (95%CI 0-46%) in the southern region. The estimates were all based on samples are from September and October, so after transmission was detected. No significant changes in the blackbird population size were observed prior to 2016. We therefore assume that the maximum amount of transmission prior to 2016 was the seroprevalence in 2016 + 1*SE and use this as the upper bound of the historical FOI. This corresponds to a historical FOI of 0.30.

#### Relative biting rate on additional reservoir host compared to blackbirds

Definition The relative frequency at which the additional reservoir host receives bites compared to blackbirds. This is a consequence of both biting preference and differences in bird density.

Prior 0 to 50

Justification A large prior distribution was chosen here, because it is difficult to estimate as the distribution of mosquito bites is a consequence of biting preference and relative host abundance. The reservoir host represents several possible (unknown) host species, making it difficult to estimate its relative abundance and attractiveness to mosquitoes. We therefore chose a wide prior and ensured that the posterior distribution was fully informed by the data and not restricted by the prior distribution.

#### Reservoir population’s lifespan

Definition Mean lifespan of the additional reservoir population. The inverse of this value represents the daily mortality rate.

Prior 365 to 7305 days (1 to 20 years)

Justification As we wanted to characterise this population from the epidemiological data, we selected a wide prior that contains the vast majority of plausible values.

## Supplementary Material E: Validation of inference approach

We created five different simulated datasets, using values shown in Supplementary table 7. These datasets were created from the blackbird-only model (model A). A total of five parameters were estimated: 1) re-emergence rate, the fraction of the USUV prevalence in mosquitoes in September that persists to the subsequent spring, 2) transmission probability bird-to-mosquito, 3) abundance scaling parameter, this converts the host and vector abundance to be on the same scale, meaning that their ratio reflects the vector-to-host ratio, 4) infection mortality rate in blackbirds, 5) historical FOI, to quantify transmission prior to first detection leading to non-zero immunity levels in 2016.

The ABC-SMC algorithm, hyperparameters, and summary statistics were identical to those used for fitting to the real surveillance data, see Supplementary Material C.

**Supplementary table 7:** Parameter values used to create simulated datasets for inference validation.

| Set | Re-emergence rate | Transmission probability bird to mosquito | Abundance scaling parameter | Infection mortality rate | Historical FOI |
| --- | --- | --- | --- | --- | --- |
| A | 0.25 | 0.7 | 0.20 | 0.2 | 0.25 |
| B | 0.25 | 0.8 | 0.15 | 0.3 | 0.08 |
| C | 0.20 | 0.9 | 0.40 | 0.2 | 0.20 |
| D | 0.20 | 0.5 | 0.15 | 0.1 | 0.04 |
| E | 0.10 | 0.6 | 0.10 | 0.4 | 0.01 |

### Inference validation results

The re-emergence rate, abundance scaling parameter, transmission probability, and infection mortality rate were well estimated with 18/20 (90%) true values falling within the estimated 95% highest density intervals (HDI) of the posterior distributions across the five datasets (*Supplementary figure 10*). We calculated concordance coefficients to quantify the agreement between parameter’s true values and the estimated median values from the posterior (*Supplementary figure 11*). We found that the abundance scaling parameter and infection mortality rate were best approximated by the posterior median. The inference framework was unable to estimate the values of the historical FOI within the range of the chosen prior. This was likely due to the limited number of data points available for seroprevalence in 2016 (no data before September). The inference framework was therefore not able to distinguish between models with seroprevalence levels due to transmission prior to 2016 or due to transmission in 2016. This parameter was still kept to ensure that the uncertainty around historical transmission is reflected in model results.

**Supplementary figure 10:**
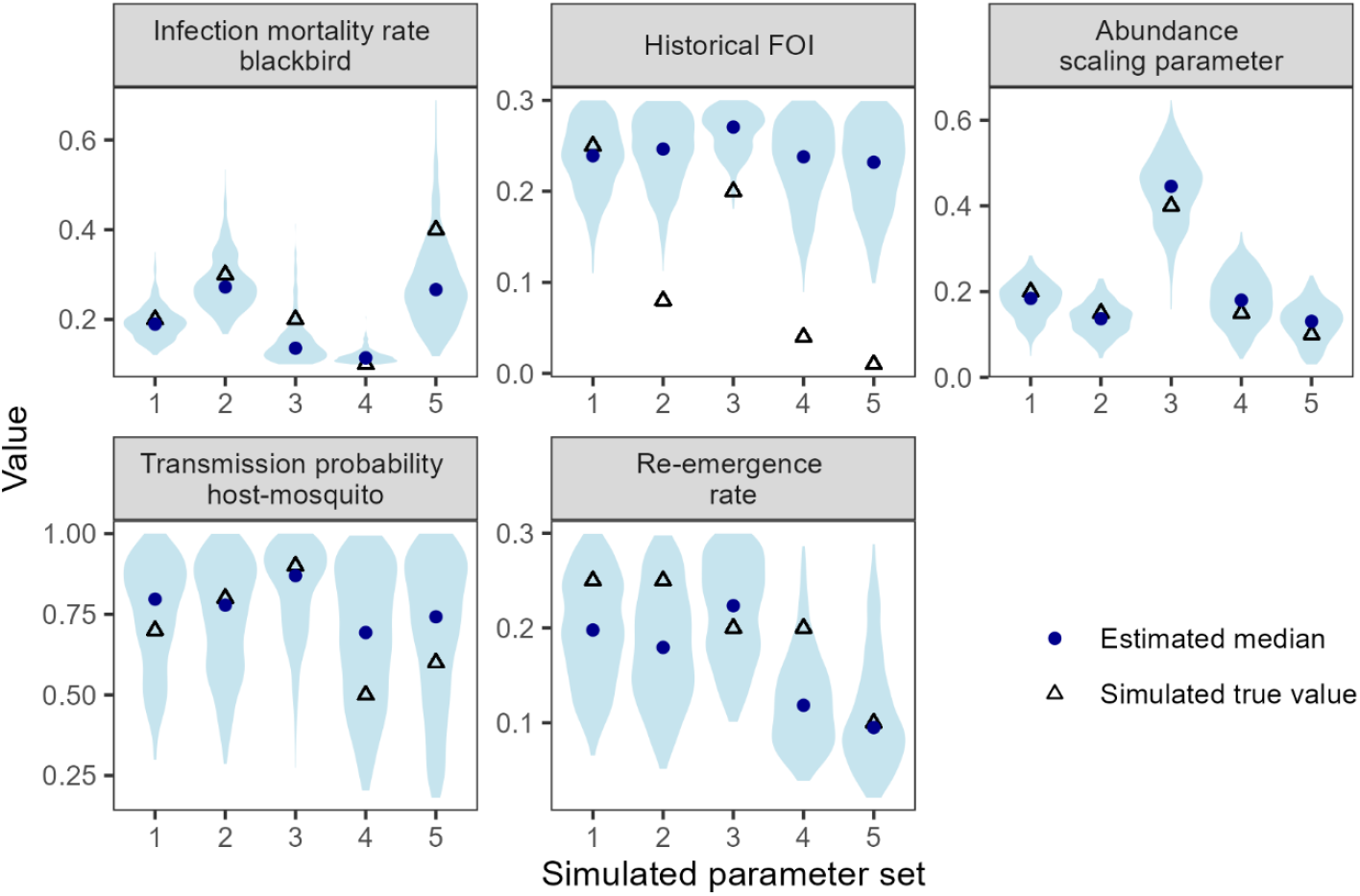
Estimated parameters from five simulated datasets (1-5) with the true value used in the simulation (blue triangle), the estimated median (blue circles) and posterior distributions (blue violins). Each estimate was based on two independent chains with 200 accepted particles each.

**Supplementary figure 11:**
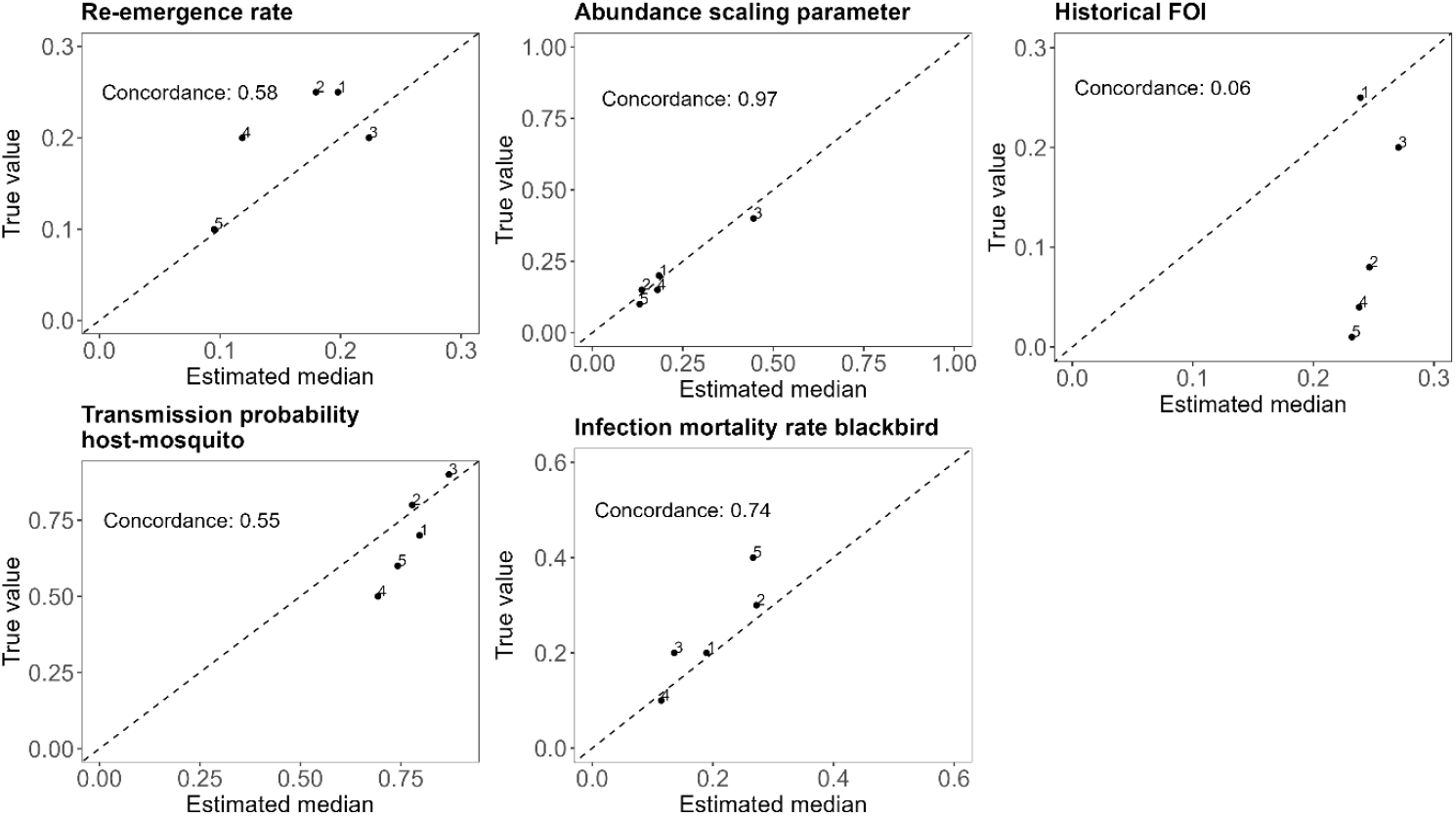
Concordance between parameter’s true values and the estimated posterior median values across five simulated datasets (1-5). Concordance represents the strength of the relationship y=x. A value of 1 indicates perfect agreement, while 0 indicates poor agreement.

## Supplementary Material F: Additional results

### Seroprevalence

**Supplementary figure 12:**
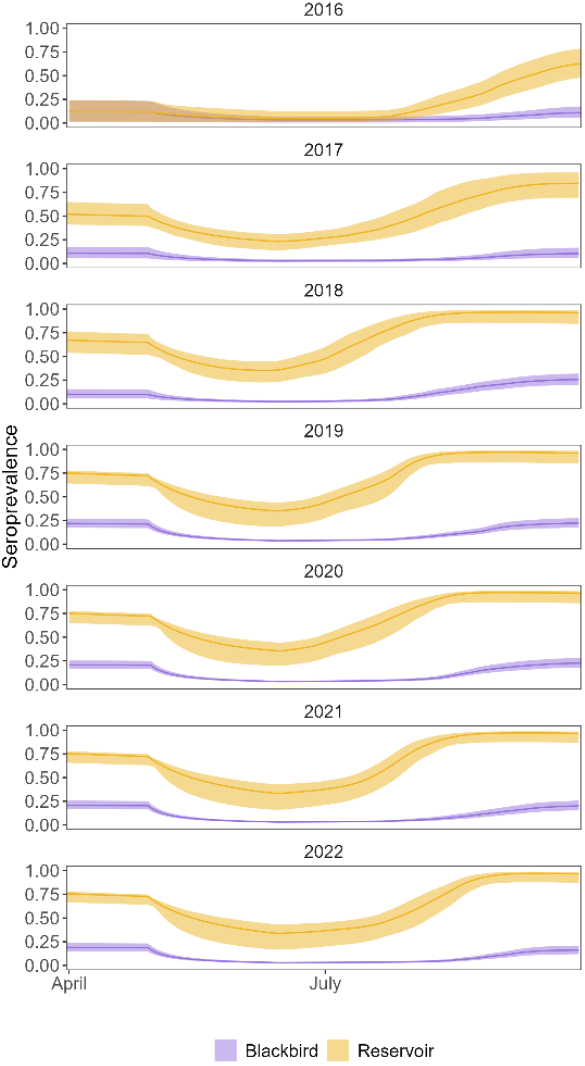
Predicted seroprevalence in blackbird and reservoir population. Model predictions are based on 100 simulations using random draws from parameter sets from posterior distributions. Data are presented as mean (solid line) with 95% prediction intervals.

### Model comparison for sensitivity analyses

**Supplementary figure 13:**
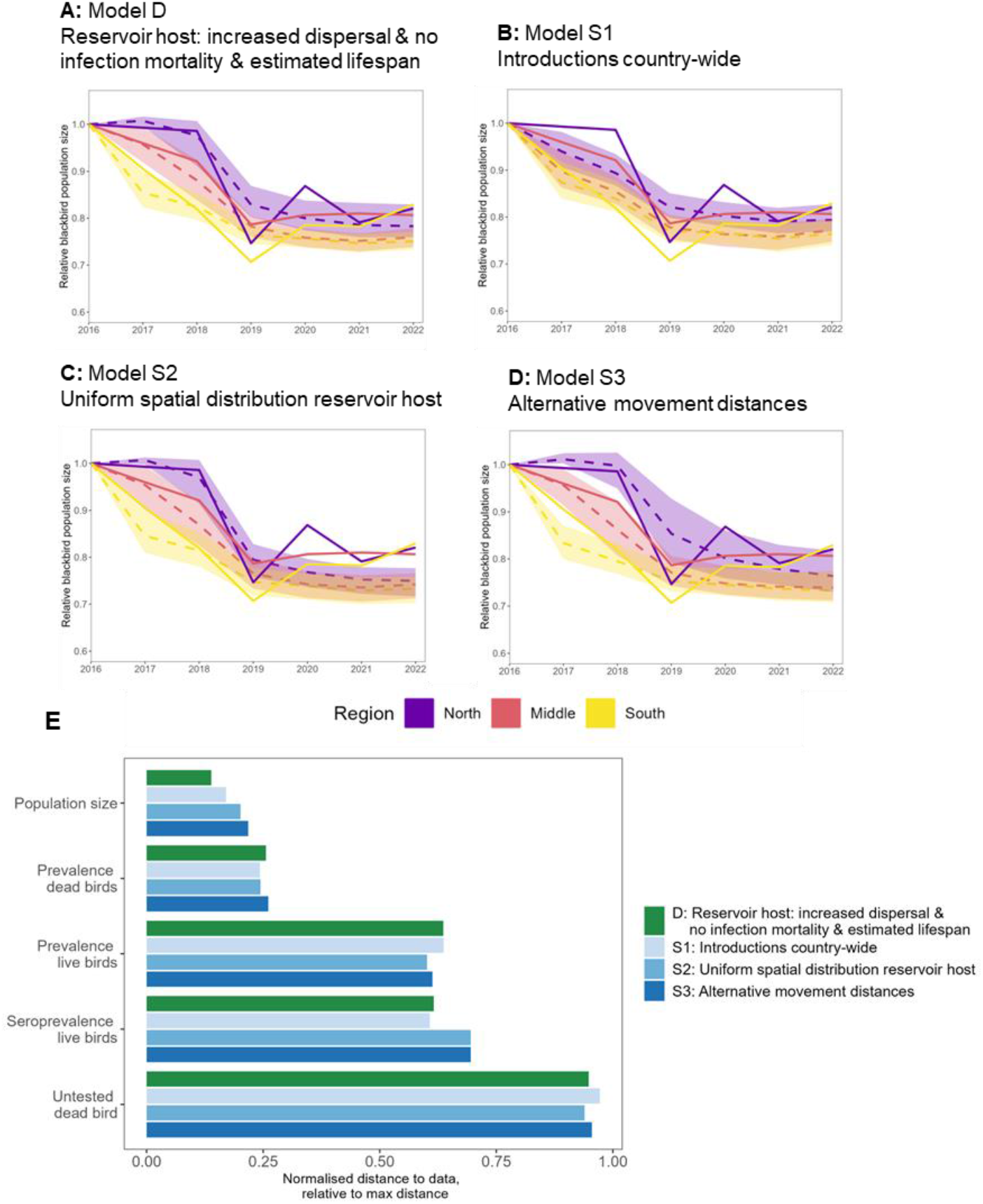
Comparison of model versions for sensitivity analyses. A-D) Visual fit of simulated data (dashed lines) to observed data (solid lines) for the blackbird population size. Simulated data are presented as mean values with 95% prediction intervals. Results are shown for the best-fitting model (main model D), and the three sensitivity analysis models S1-S3. E) Comparison of model fits between best-fitting model and sensitivity analysis models. Comparisons were based on distances between observed and simulated data for each data set. For each data set, distances were normalised by dividing each model’s distance by the maximum across all model versions. A normalised distance of 1 thus indicates that this model showed the largest distance between observed and simulated data (i.e. worst fit). Lower normalised distances indicate a better fit to the data.

### Fits of main model (model D)

**Supplementary figure 14:**
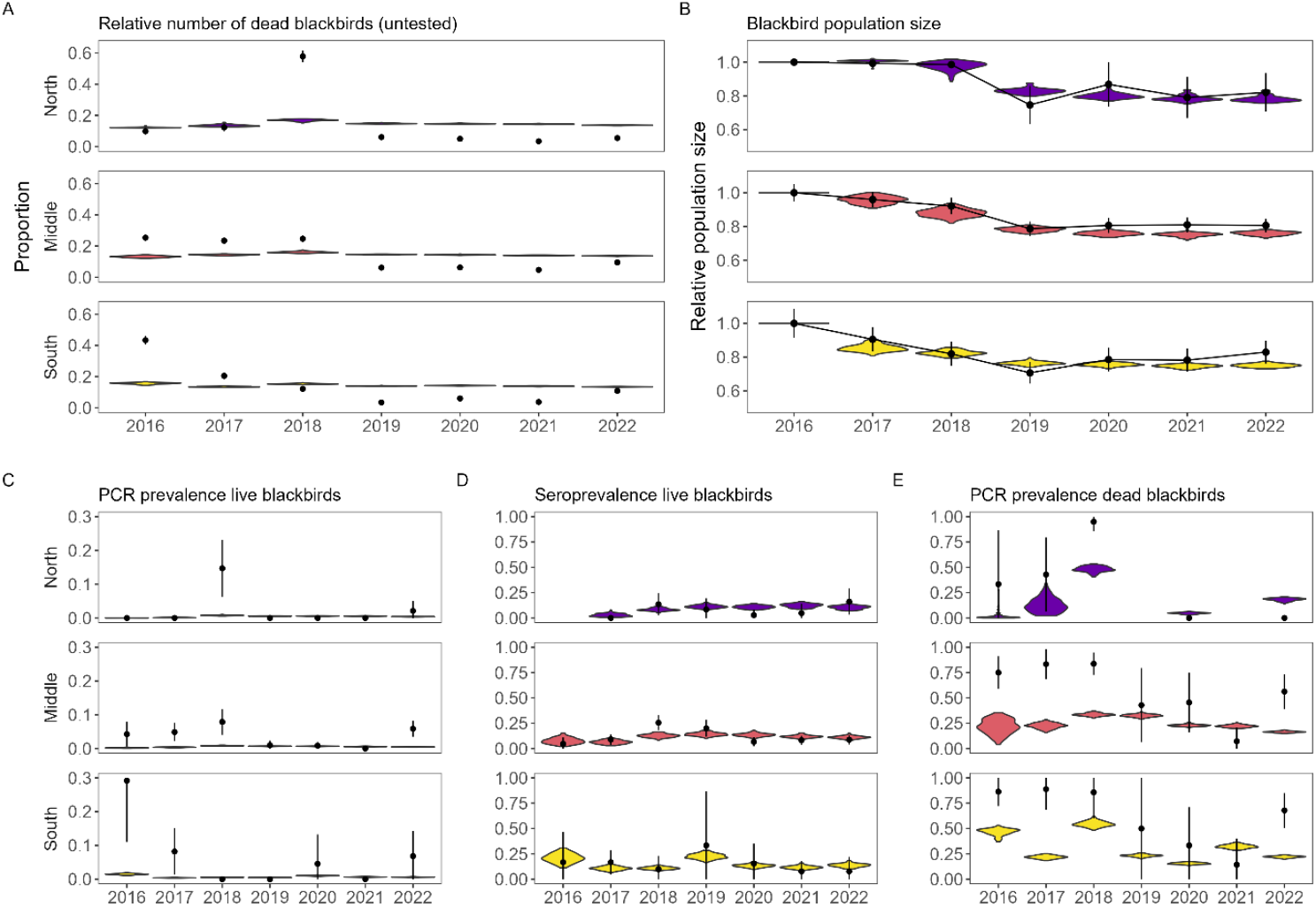
Resulting model fits from best-fitting model (model D) obtained by comparing model predictions (violins) to observed data (in black, with dots representing the mean and vertical lines representing binomial 95% confidence intervals). Model predictions are based on 100 simulations using random draws from parameter sets from posterior distributions. Results are aggregated across time to annual level. Results on within-year level are available in Supplementary figure 15. For figures C-E, predictions are only calculated for those months and regions when surveillance data was available. The period for which model predictions were calculated therefore varies between years and regions to ensure alignment with the observation process.

### Model fits over time for model D

In addition to the regional disaggregation of model fits presented in Supplementary figure 14, the prevalence-based patterns could also be disaggregated to monthly estimates. Fits to within-year trends in the data are shown for these three summary statistics (Supplementary figure 15).

**Supplementary figure 15:**
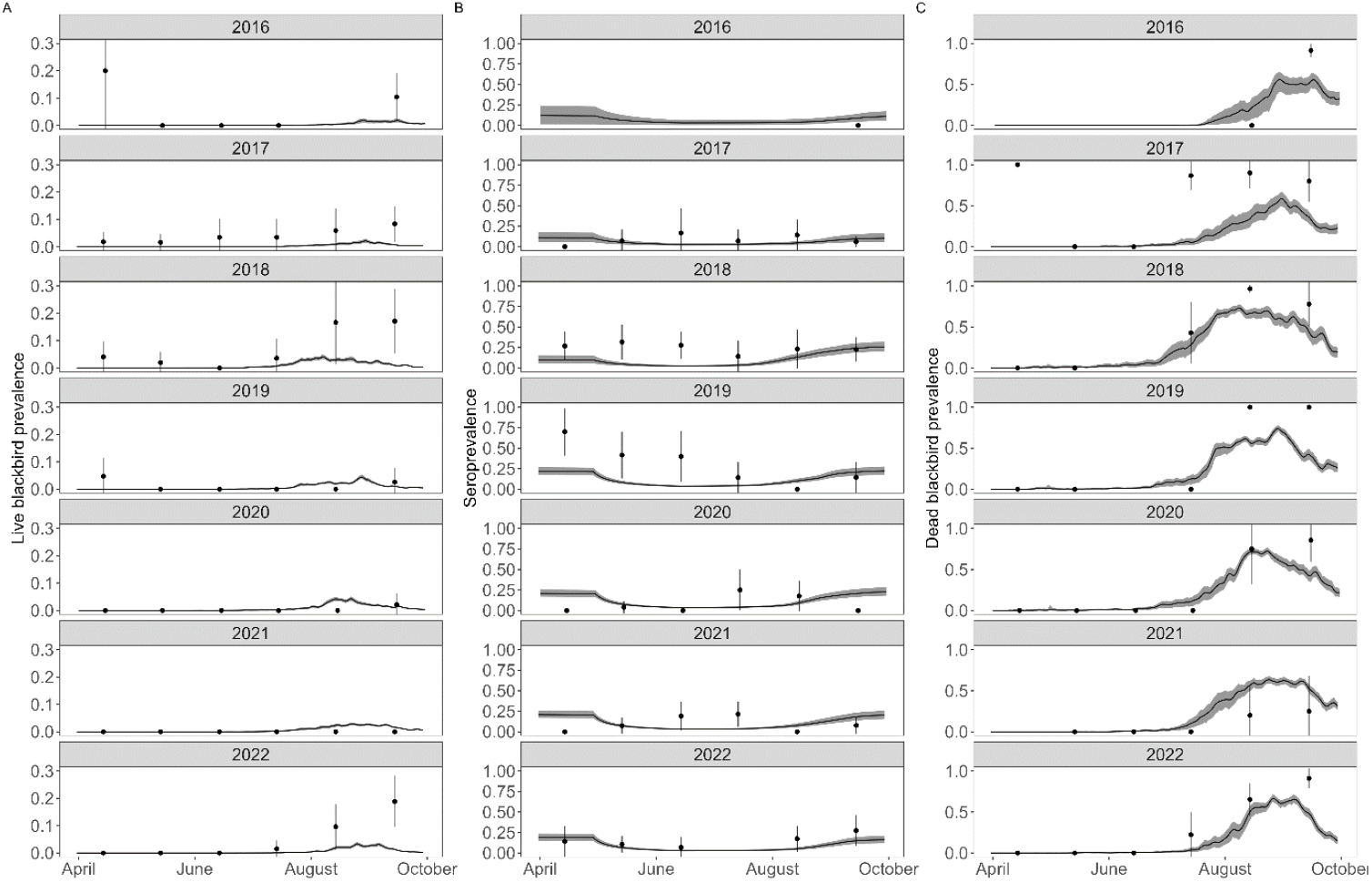
Comparison of observed (dots with vertical lines representing 95% confidence intervals) and simulated data from best-fitting model at monthly level for A) live blackbird infection prevalence, B) live blackbird seroprevalence, and C) dead blackbird infection prevalence. Model predictions are based on 100 simulations using random draws from parameter sets from posterior distributions. Observed data are aggregated to monthly level. Missing dots indicate no data were available for this month. When data were available but no positive samples were detected during a specific month, no confidence intervals are shown.

### Model fits of alternative models

#### Below we show the model fits for all alternative models (models A-C and S1-S3)

**Supplementary Figure 16:**
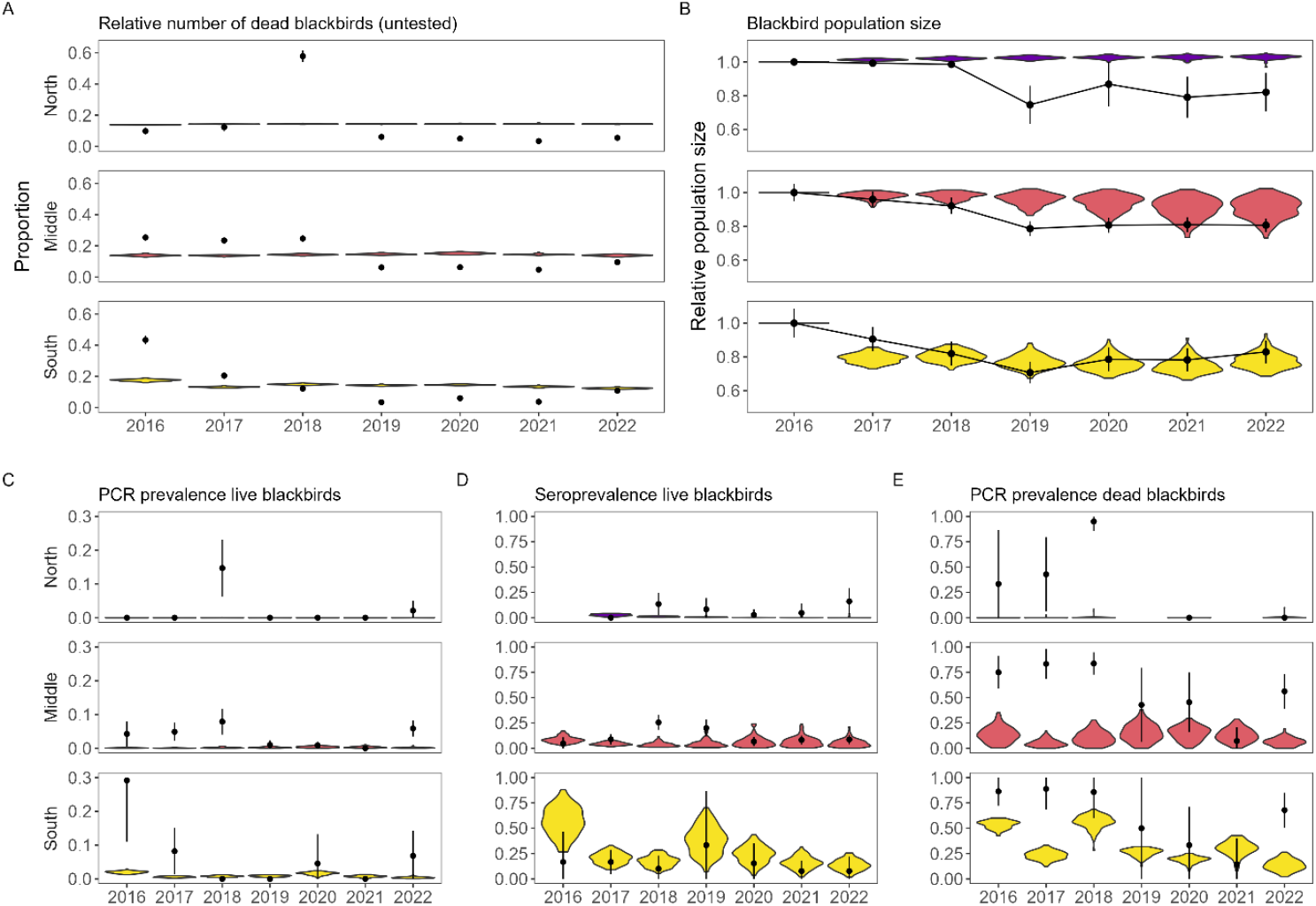
Resulting model fits from model A obtained by comparing model predictions (violins) to observed data (in black, with dots representing the mean and vertical lines representing binomial 95% confidence intervals). Model predictions are based on 100 simulations using random draws from parameter sets from posterior distributions. Results are aggregated across time to annual level. For figures C-E, predictions are only calculated for those months and regions when surveillance data was available. The period for which model predictions were calculated therefore varies between years and regions to ensure alignment with the observation process.

**Supplementary figure 17:**
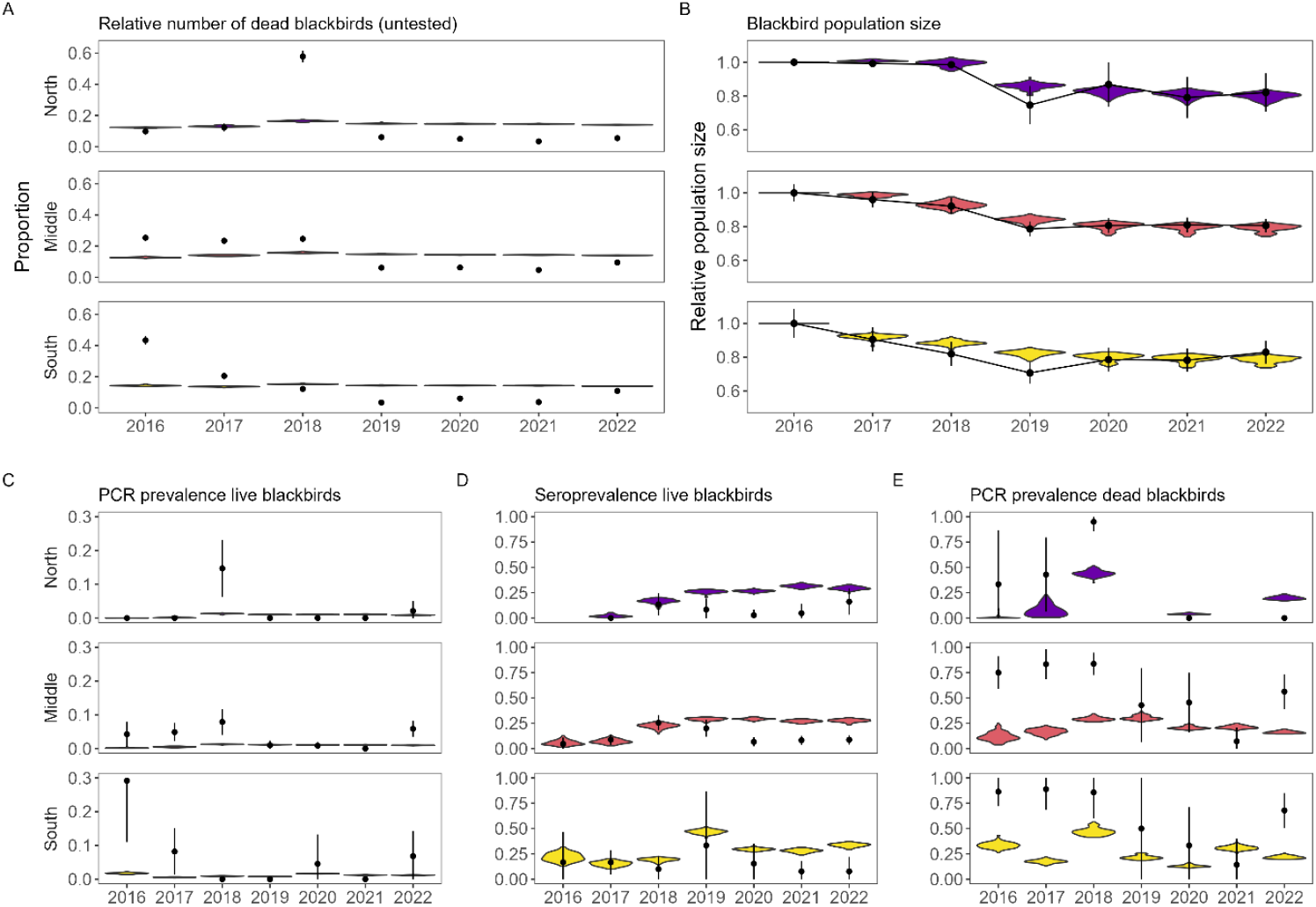
Resulting model fits from model B obtained by comparing model predictions (violins) to observed data (in black, with dots representing the mean and vertical lines representing binomial 95% confidence intervals). Model predictions are based on 100 simulations using random draws from parameter sets from posterior distributions. Results are aggregated across time to annual level. For figures C-E, predictions are only calculated for those months and regions when surveillance data was available. The period for which model predictions were calculated therefore varies between years and regions to ensure alignment with the observation process.

**Supplementary figure 18:**
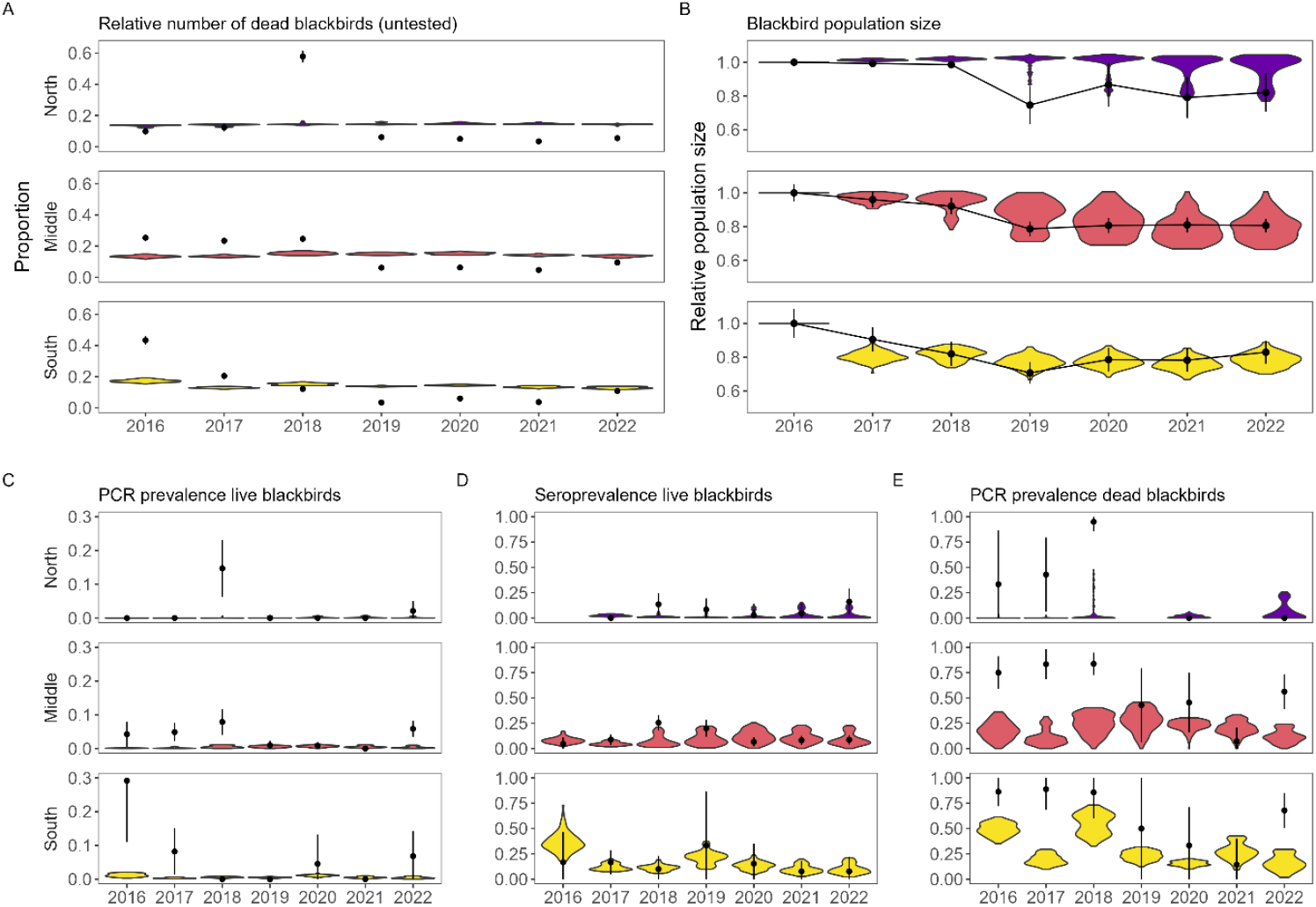
Resulting model fits from model C obtained by comparing model predictions (violins) to observed data (in black, with dots representing the mean and vertical lines representing binomial 95% confidence intervals). Model predictions are based on 100 simulations using random draws from parameter sets from posterior distributions. Results are aggregated across time to annual level. For figures C-E, predictions are only calculated for those months and regions when surveillance data was available. The period for which model predictions were calculated therefore varies between years and regions to ensure alignment with the observation process.

**Supplementary figure 19:**
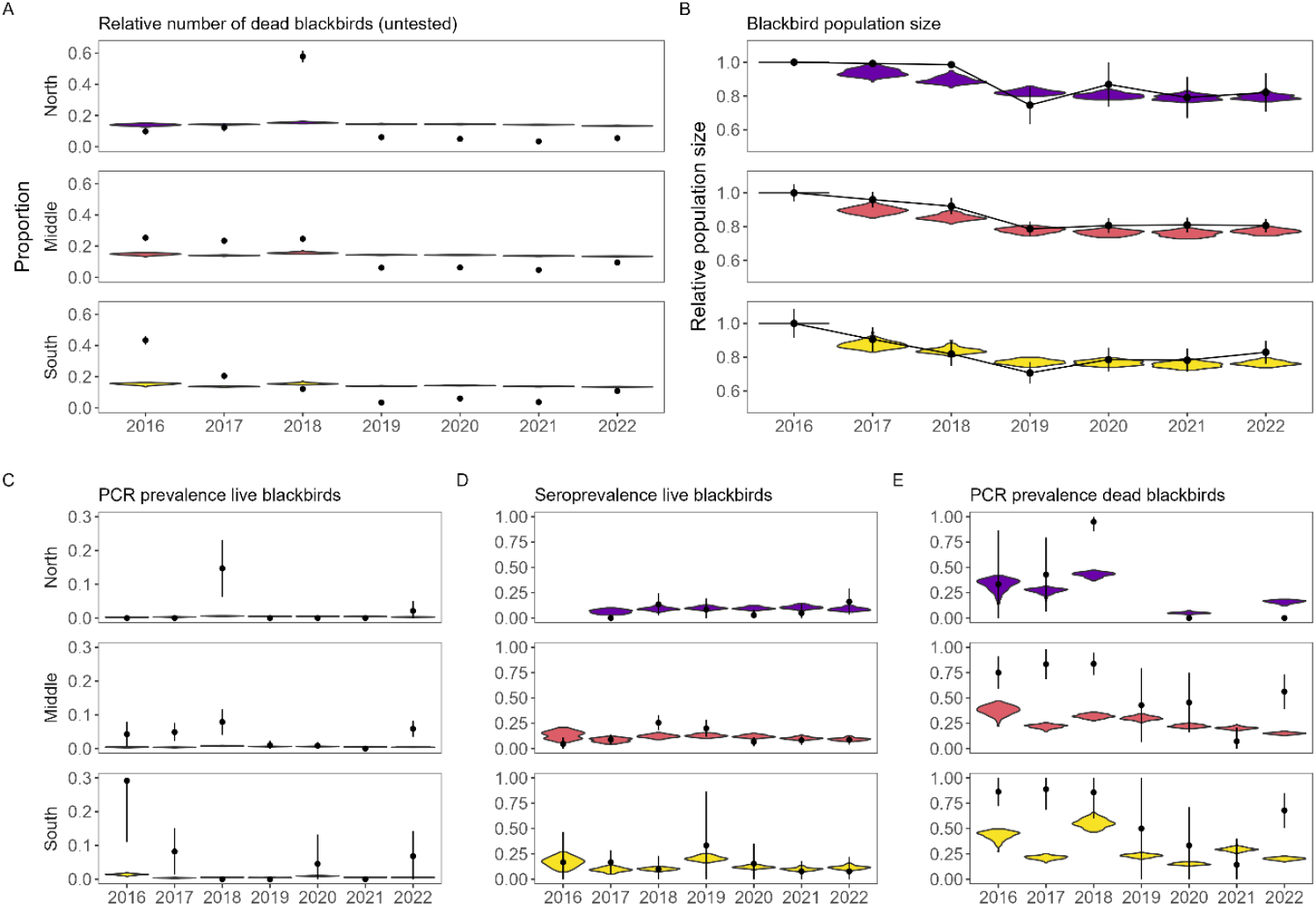
Resulting model fits from model S1 obtained by comparing model predictions (violins) to observed data (in black, with dots representing the mean and vertical lines representing binomial 95% confidence intervals). Model predictions are based on 100 simulations using random draws from parameter sets from posterior distributions. Results are aggregated across time to annual level. For figures C-E, predictions are only calculated for those months and regions when surveillance data was available. The period for which model predictions were calculated therefore varies between years and regions to ensure alignment with the observation process.

**Supplementary figure 20:**
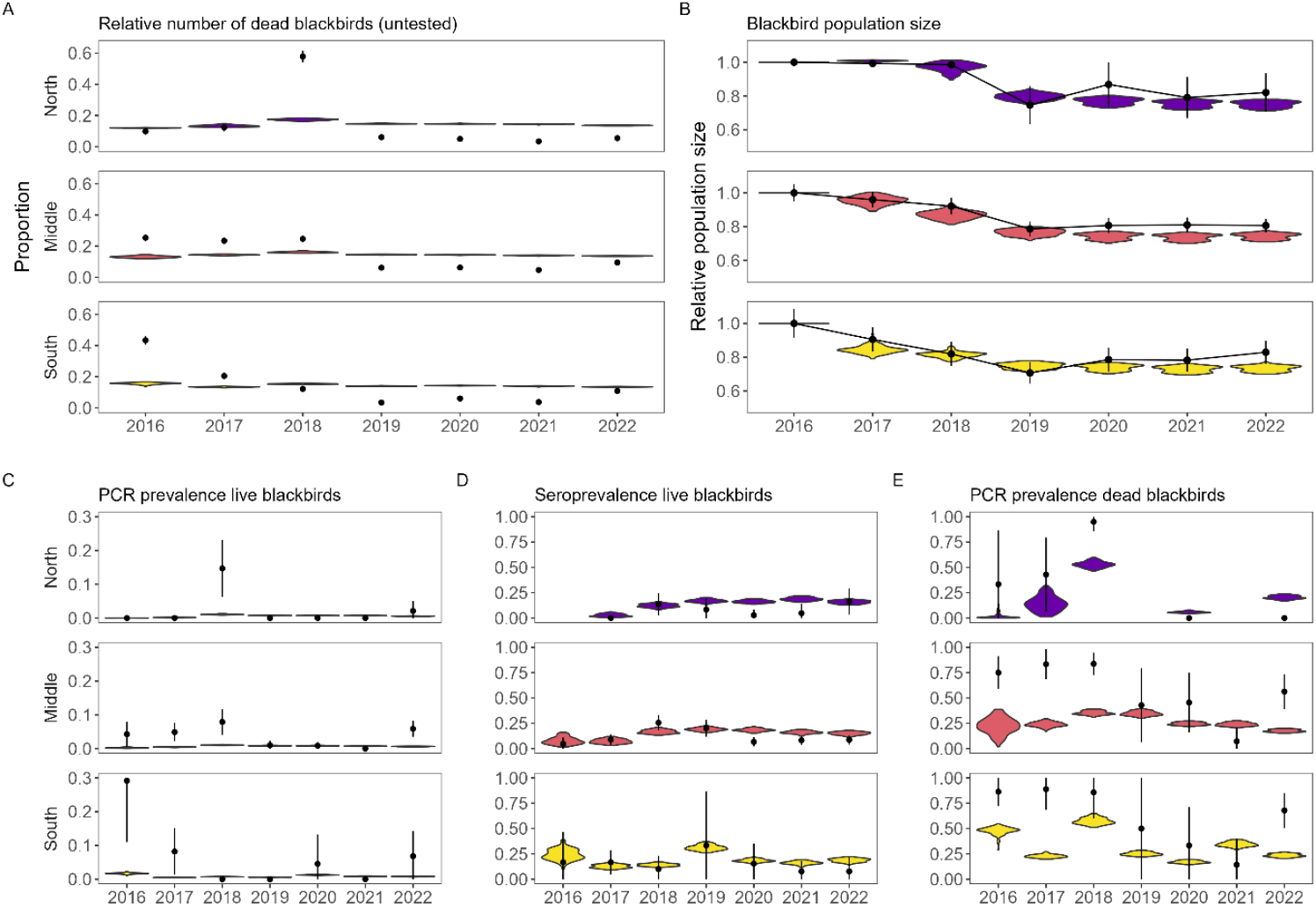
Resulting model fits from model S2 obtained by comparing model predictions (violins) to observed data (in black, with dots representing the mean and vertical lines representing binomial 95% confidence intervals). Model predictions are based on 100 simulations using random draws from parameter sets from posterior distributions. Results are aggregated across time to annual level. For figures C-E, predictions are only calculated for those months and regions when surveillance data was available. The period for which model predictions were calculated therefore varies between years and regions to ensure alignment with the observation process.

**Supplementary figure 21:**
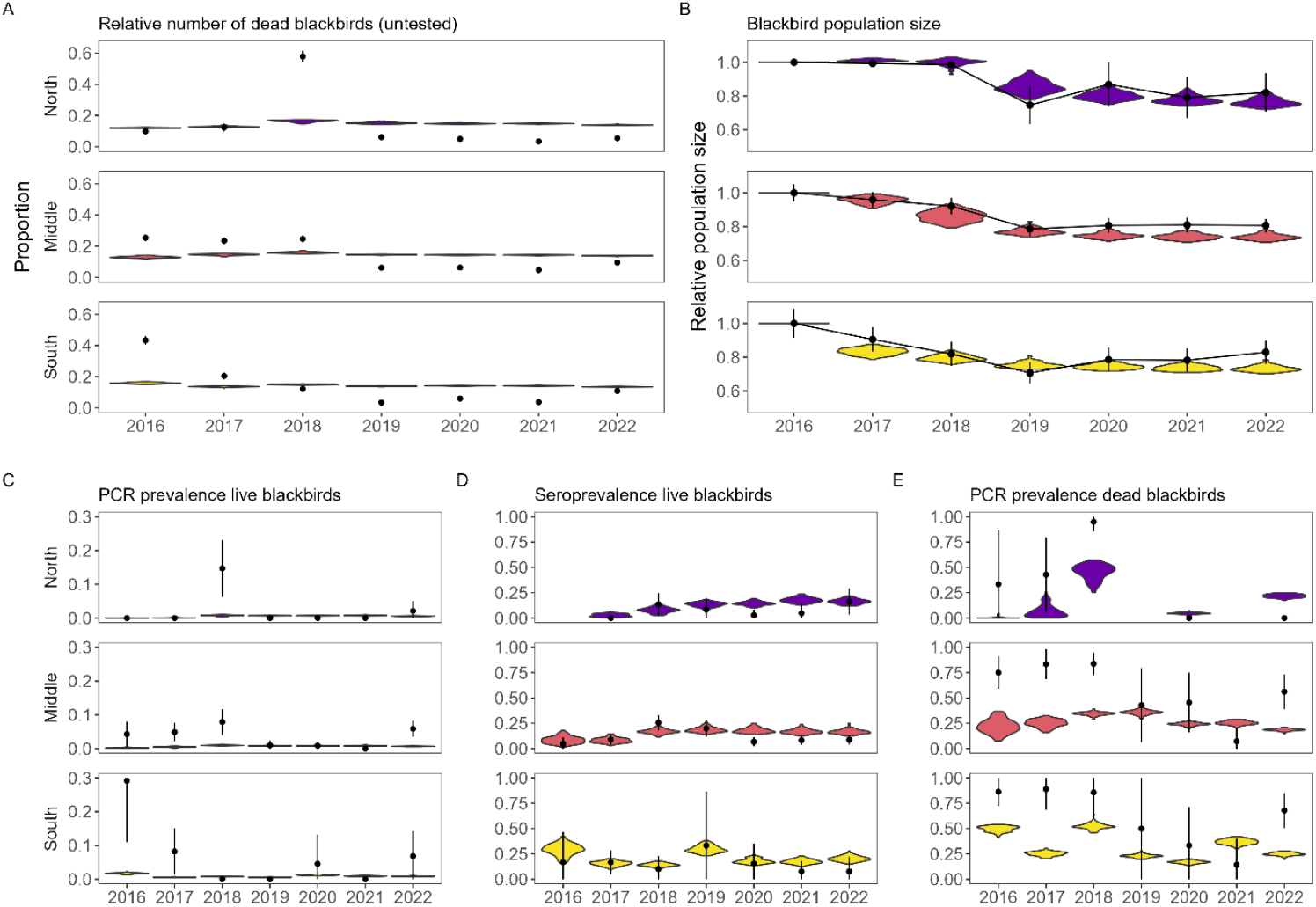
Resulting model fits from model S3 obtained by comparing model predictions (violins) to observed data (in black, with dots representing the mean and vertical lines representing binomial 95% confidence intervals). Model predictions are based on 100 simulations using random draws from parameter sets from posterior distributions. Results are aggregated across time to annual level. For figures C-E, predictions are only calculated for those months and regions when surveillance data was available. The period for which model predictions were calculated therefore varies between years and regions to ensure alignment with the observation process.

### Posterior distributions of all models

**Supplementary table 8:**
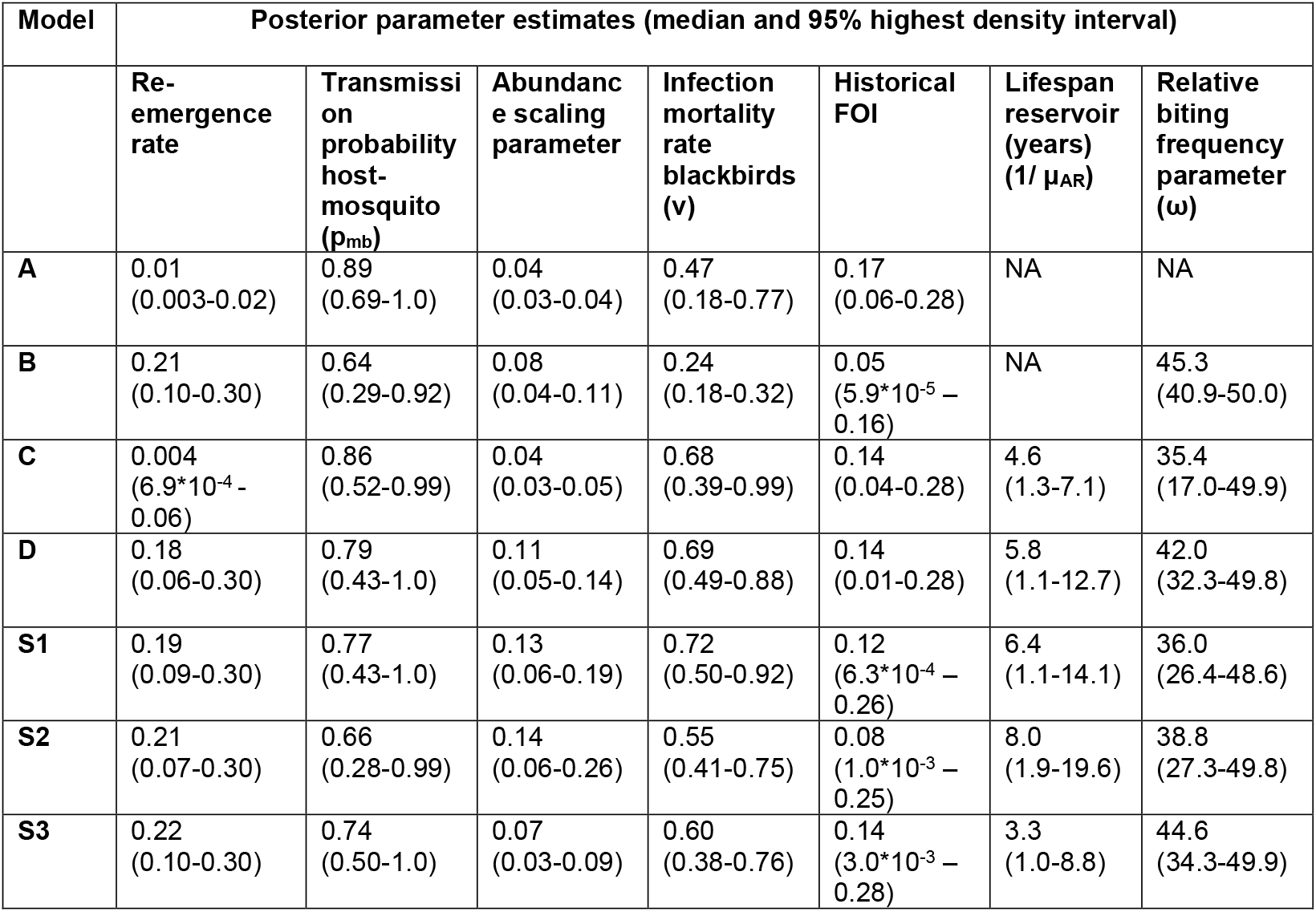
Posterior estimates for each model version. Values shown are median estimates with 95% highest density intervals.

| Model | Posterior parameter estimates (median and 95% highest density interval) |  |  |  |  |  |  |
| --- | --- | --- | --- | --- | --- | --- | --- |
| | Re-emergence rate | Transmission probability host-mosquito ($p_{mb}$) | Abundance scaling parameter | Infection mortality rate blackbirds ($v$) | Historical FOI | Lifespan reservoir (years) ($1/\mu_{AR}$) | Relative biting frequency parameter ($\omega$) |
| **A** | 0.01<br>(0.003-0.02) | 0.89<br>(0.69-1.0) | 0.04<br>(0.03-0.04) | 0.47<br>(0.18-0.77) | 0.17<br>(0.06-0.28) | NA | NA |
| **B** | 0.21<br>(0.10-0.30) | 0.64<br>(0.29-0.92) | 0.08<br>(0.04-0.11) | 0.24<br>(0.18-0.32) | 0.05<br>($5.9 \times 10^{-5}$ – 0.16) | NA | 45.3<br>(40.9-50.0) |
| **C** | 0.004<br>($6.9 \times 10^{-4}$ - 0.06) | 0.86<br>(0.52-0.99) | 0.04<br>(0.03-0.05) | 0.68<br>(0.39-0.99) | 0.14<br>(0.04-0.28) | 4.6<br>(1.3-7.1) | 35.4<br>(17.0-49.9) |
| **D** | 0.18<br>(0.06-0.30) | 0.79<br>(0.43-1.0) | 0.11<br>(0.05-0.14) | 0.69<br>(0.49-0.88) | 0.14<br>(0.01-0.28) | 5.8<br>(1.1-12.7) | 42.0<br>(32.3-49.8) |
| **S1** | 0.19<br>(0.09-0.30) | 0.77<br>(0.43-1.0) | 0.13<br>(0.06-0.19) | 0.72<br>(0.50-0.92) | 0.12<br>($6.3 \times 10^{-4}$ – 0.26) | 6.4<br>(1.1-14.1) | 36.0<br>(26.4-48.6) |
| **S2** | 0.21<br>(0.07-0.30) | 0.66<br>(0.28-0.99) | 0.14<br>(0.06-0.26) | 0.55<br>(0.41-0.75) | 0.08<br>($1.0 \times 10^{-3}$ – 0.25) | 8.0<br>(1.9-19.6) | 38.8<br>(27.3-49.8) |
| **S3** | 0.22<br>(0.10-0.30) | 0.74<br>(0.50-1.0) | 0.07<br>(0.03-0.09) | 0.60<br>(0.38-0.76) | 0.14<br>($3.0 \times 10^{-3}$ – 0.28) | 3.3<br>(1.0-8.8) | 44.6<br>(34.3-49.9) |

## Supplementary Material G: Inference diagnostics

A range of ABC diagnostics are shown in figures below. These are all based on the best-fitting model, model D.

**Supplementary Figure 22:**
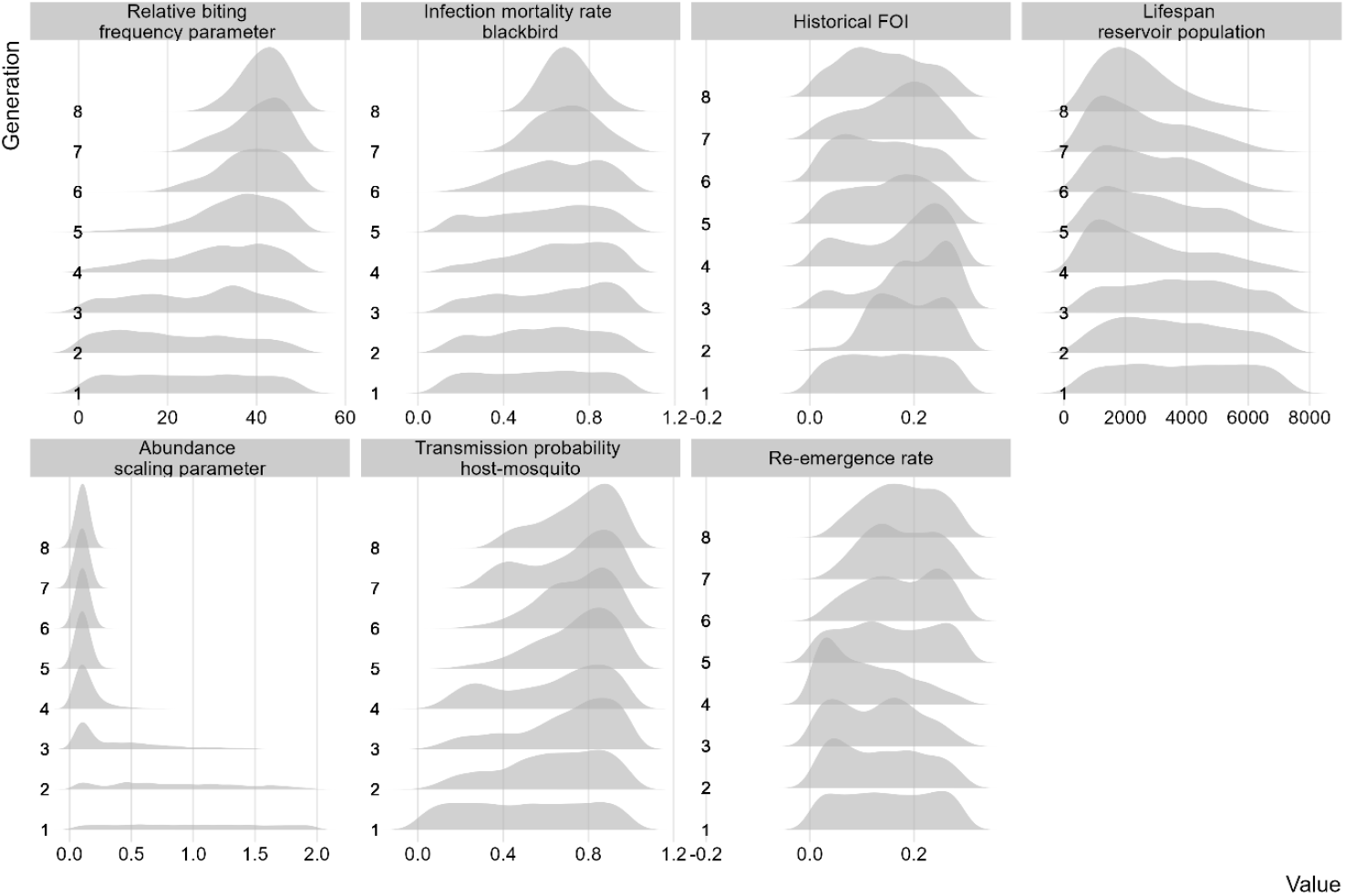
Convergence of intermediate distributions towards the posterior distribution per generation for model D.

**Supplementary figure 23:**
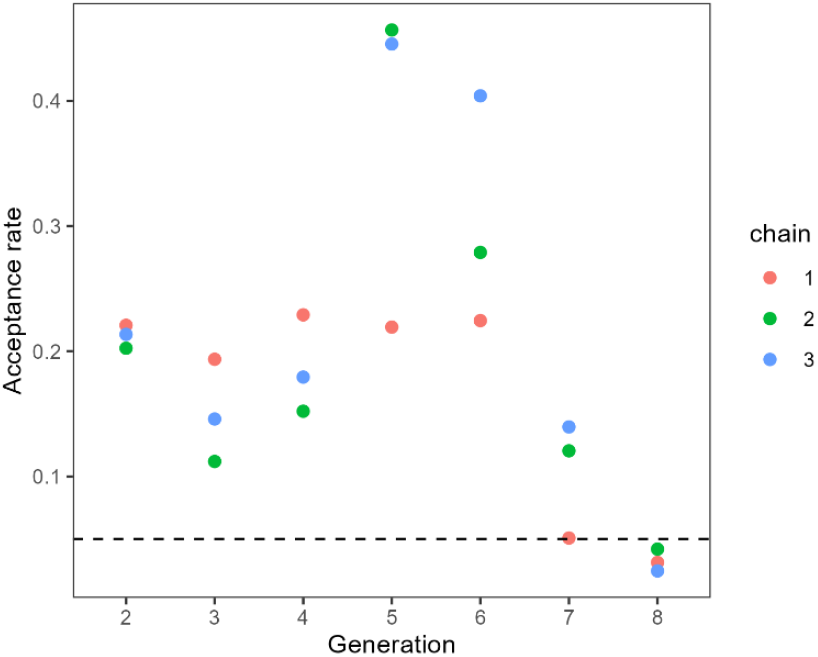
Acceptance rate of each chain per generation for model D. The horizontal dashed line represents the stopping criterion of a 5% acceptance rate.

**Supplementary figure 24:**
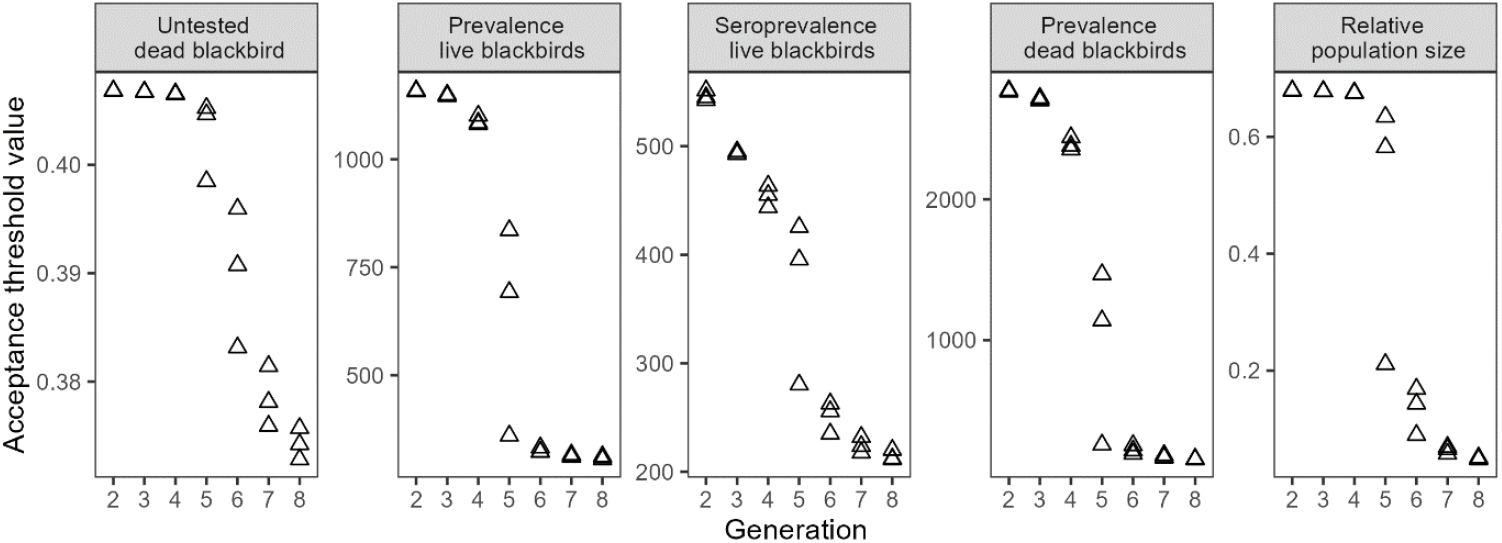
Development of increasingly strict threshold across generations for each of the summary statistics.

**Supplementary figure 25:**
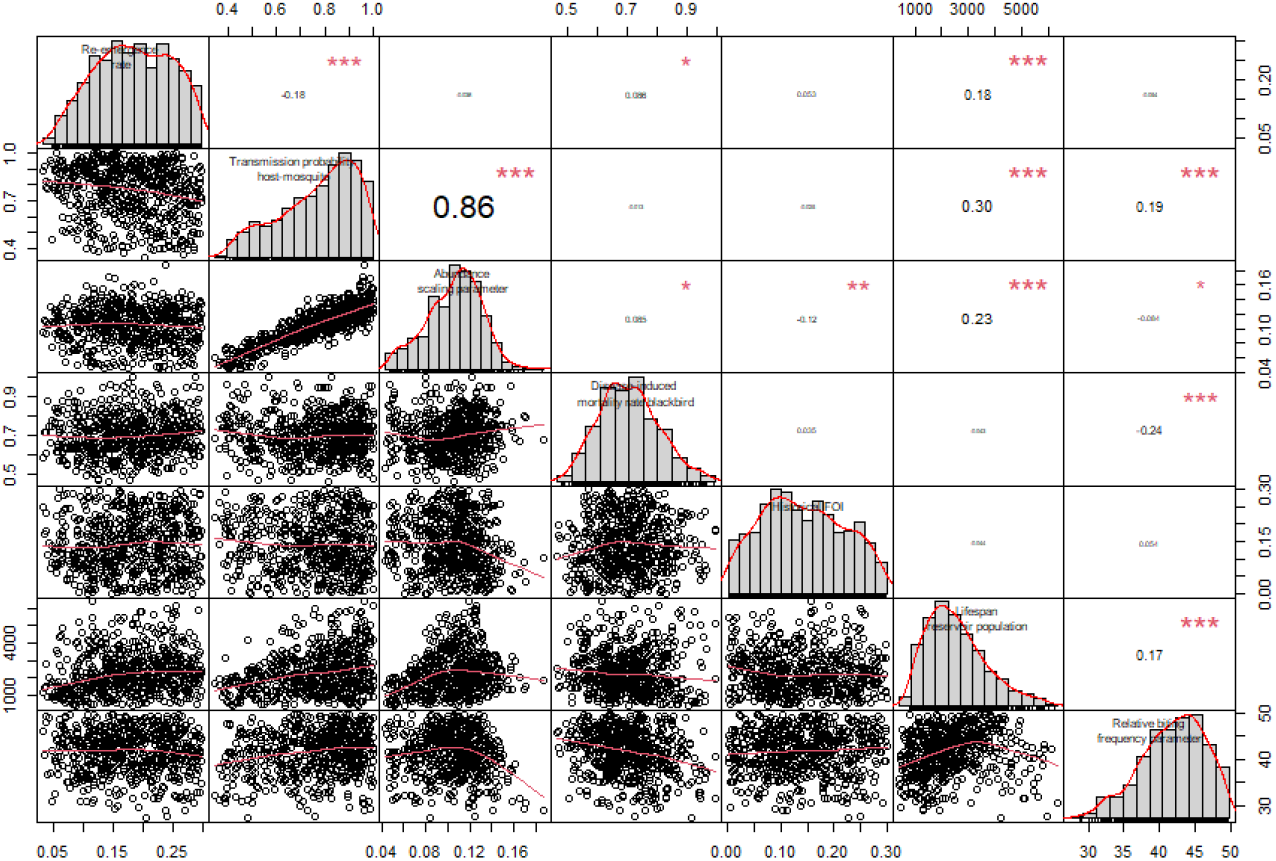
Correlation plot of posterior distributions for model D.

## Supplementary Material H: Reproduction numbers

### Derivation of reproduction numbers

Reproduction numbers were calculated using the Next Generation Matrix (NGM) approach. The reproduction number was obtained by calculating the dominant eigenvalue of the NGM. The dimensions of the NGM were 5×5 representing the number of species capable of transmitting the infection (mosquitoes (M), juvenile blackbirds (J), adult blackbirds (A), juvenile reservoir population (R), adult reservoir population (P)). Each element in the NGM, *kij*, represents the expected number of cases of type *i* caused by an individual of type *j*:

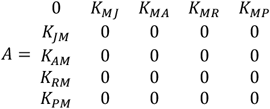

All mosquito-to-mosquito and bird-to-bird elements in the NGM are set to zero as transmission only occurred between mosquitoes and birds and vice versa. The value of each of the elements in the NGM was derived as follows:

K_JM_: The number of infected juvenile blackbirds caused by an infectious mosquito is determined by

- The fraction of mosquitoes transitioning from E to I
- The duration of infectiousness in mosquitoes
- The mosquito-to-juvenile blackbird transmission rate

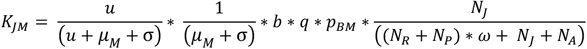

K_;AM_;: The number of infected adult blackbirds caused by an infectious mosquito is determined by

- The fraction of mosquitoes transitioning from E to I
- The duration of infectiousness in mosquitoes
- The mosquito-to-adult blackbird transmission rate

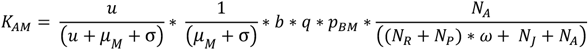

K_;RM_;: The number of infected juvenile reservoir hosts caused by an infectious mosquito is determined by

- The fraction of mosquitoes transitioning from E to I
- The duration of infectiousness in mosquitoes
- The mosquito-to-juvenile reservoir transmission rate

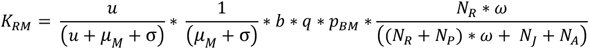

K_;PM_;: The number of infected adult reservoir hosts caused by an infectious mosquito is determined by

- The fraction of mosquitoes transitioning from E to I
- The duration of infectiousness in mosquitoes
- The mosquito-to-adult reservoir transmission rate

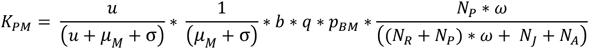

K_;MJ_;: The number of infected mosquitoes caused by an infectious juvenile blackbird is determined by

- The fraction of juvenile blackbirds transitioning from E to I
- The duration of infectiousness in juvenile blackbirds
- The juvenile blackbird-to-mosquito transmission rate

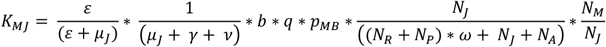

K_;MA_;: The number of infected mosquitoes caused by an infectious adult blackbird is determined by

- The fraction of adult blackbirds transitioning from E to I
- The duration of infectiousness in adults
- The adult blackbird-to-mosquito transmission rate

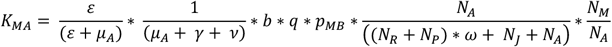

K_;MR_;: The number of infected mosquitoes caused by an infectious juvenile reservoir host is determined by

- The fraction of juvenile reservoir birds transitioning from E to I
- The duration of infectiousness in juvenile reservoir birds
- The juvenile reservoir-to-mosquito transmission rate

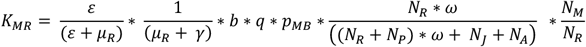

K_;MP_;: The number of infected mosquitoes caused by an infectious adult reservoir host is determined by

- The fraction of adult reservoir birds transitioning from E to I
- The duration of infectiousness in adult reservoir birds
- The adult reservoir-to-mosquito transmission rate

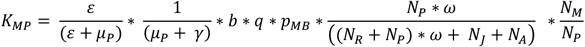

The elements in the NGM represent partial reproduction numbers, i.e. the reproduction number for a specific transmission route. The effective reproduction number was calculated by multiplying each of the partial reproduction numbers by the proportion of the population that is susceptible to infection. The effective reproduction number was therefore the dominant eigenvalue of the following matrix:

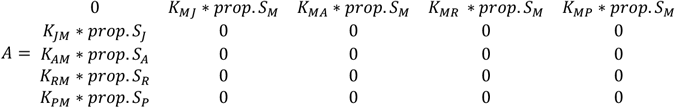

**Supplementary figure 26:**
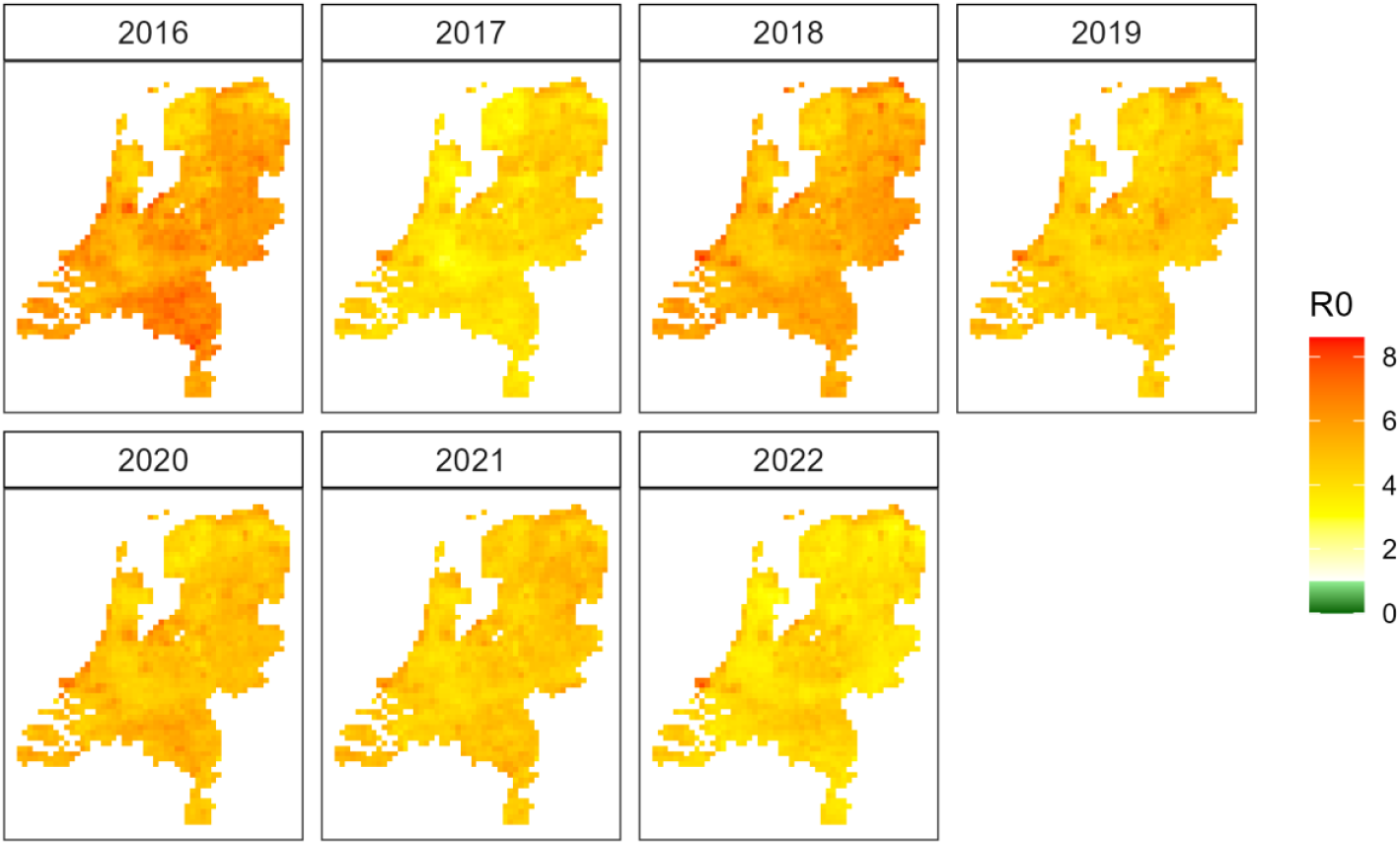
Map of mean basic reproduction number during the transmission season (April to October)

**Supplementary figure 27:**
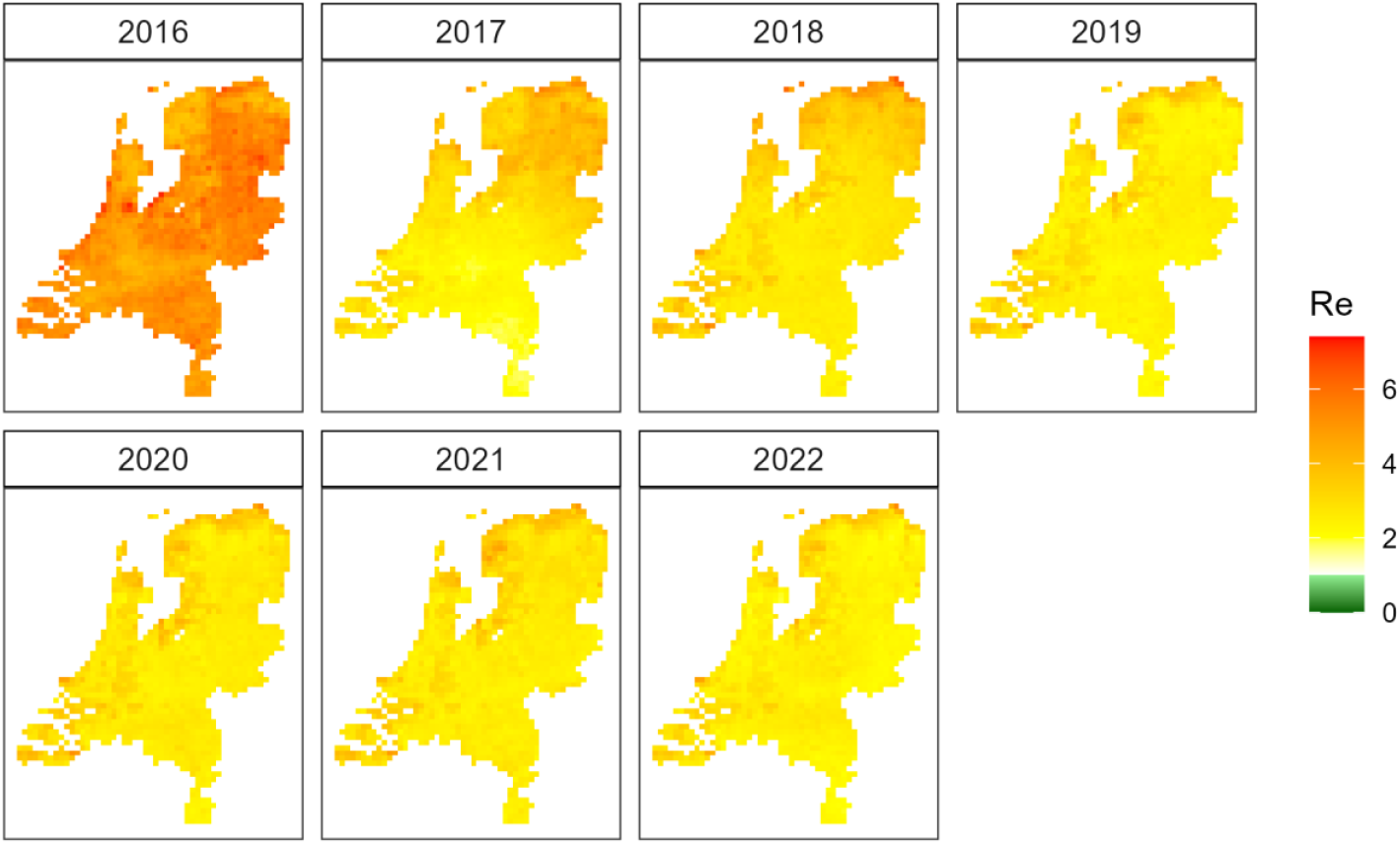
Map of mean effective reproduction number during the transmission season (April to October)

### Results related to reproduction number

**Supplementary figure 28:**
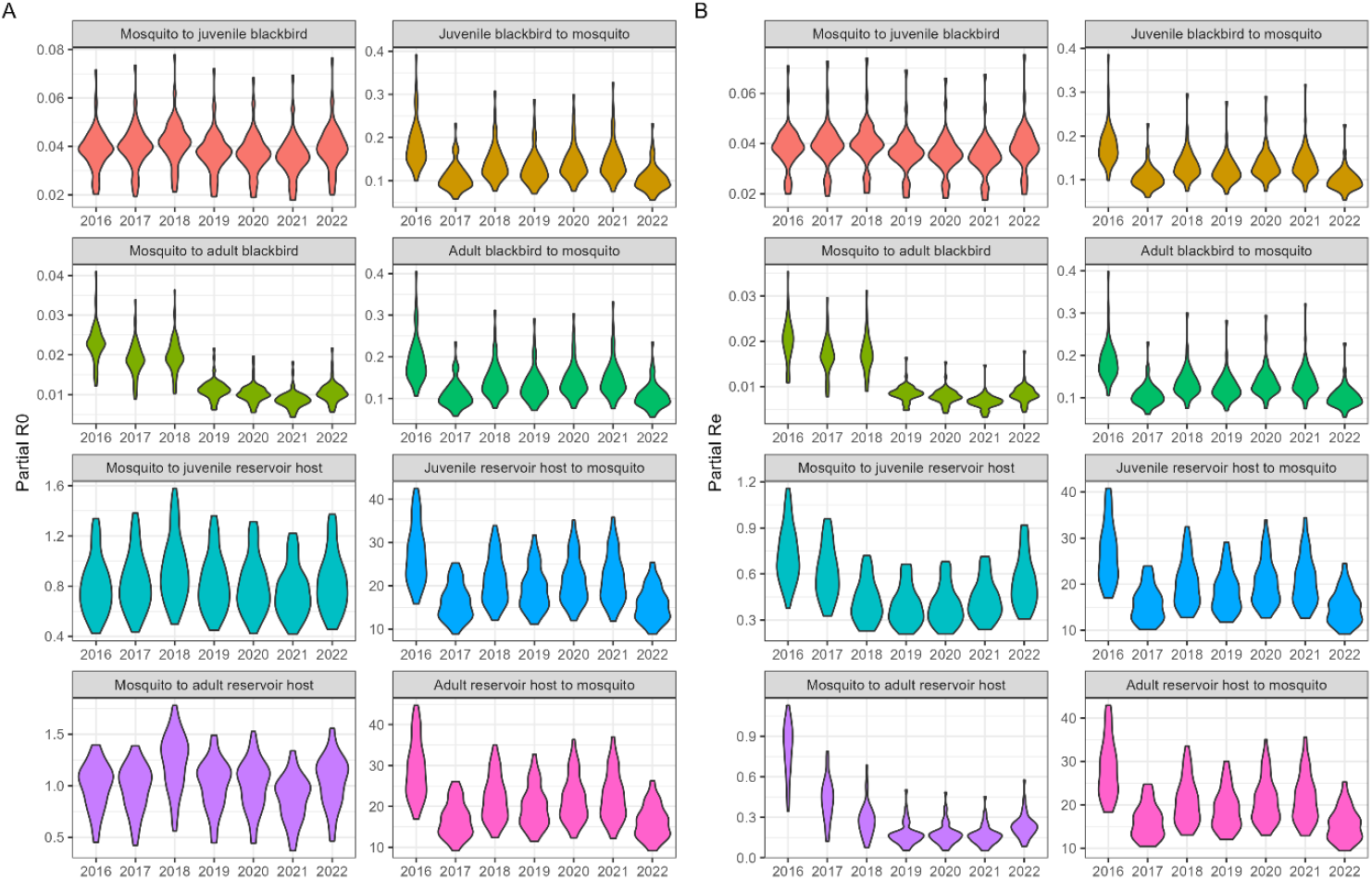
Annual partial reproduction numbers (basic (A) and effective (B)) during transmission season from April to October.

## Supplementary Material I: Model calibration without PCR datasets

The models were fitted to five sources of surveillance data (see Supplementary figure 9). To explore how much information was taken from the PCR data we fitted the main model (model D) to the three surveillance datasets that did not use PCR testing: seroprevalence in live blackbirds, reported dead blackbirds, and annual blackbird population size.

We explored its impact on model convergence and on posterior distributions. Convergence was slower in the model fitted to fewer datasets. The threshold used for acceptance of particles was consistently higher compared to the model fitted to all datasets (indicating a larger difference between observed and simulated data) and more generations were needed to reach the desired acceptance rate (Supplementary figure 29). While convergence was slower, the final threshold values were similar between both models, indicating that final model fits were similar. The resulting posterior distributions were also similar (Supplementary table 9). Median estimates of the model without PCR data all fell within the 95% highest density interval of the model fitted to all datasets.

**Supplementary figure 29:**
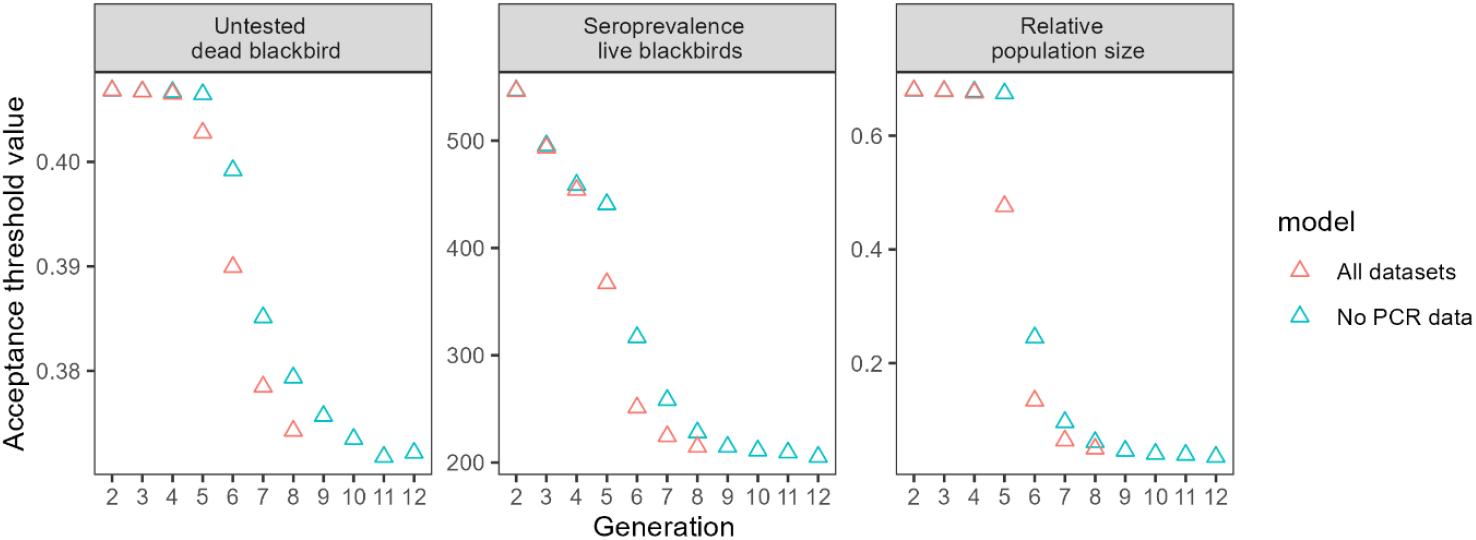
Comparison of acceptance thresholds across generations between the model fitted to all datasets (red) and the model fitted to only three datasets (blue). Values shown are the mean threshold values across all chains.

**Supplementary table 9:** Comparison of posterior estimates for model D fitted to all surveillance datasets and for model D fitted to only three datasets (PCR data removed). Values shown are median estimates with 95% highest density intervals.

| Model | Posterior parameter estimates (median and 95% highest density interval) |  |  |  |  |  |  |
| --- | --- | --- | --- | --- | --- | --- | --- |
| | Re-emergence rate | Transmission probability host-mosquito ($p_{mb}$) | Abundance scaling parameter | Infection mortality rate blackbirds ($v$) | Historical FOI | Lifespan reservoir (years) ($1/\mu_{AR}$) | Relative biting frequency parameter ($\omega$) |
| D: all data | 0.18<br>(0.06-0.30) | 0.79<br>(0.43-1.0) | 0.11<br>(0.05-0.14) | 0.69<br>(0.49-0.88) | 0.14<br>(0.01-0.28) | 5.8<br>(1.1-12.7) | 42.0<br>(32.3-49.8) |
| D: no PCR data | 0.22<br>(0.08-0.30) | 0.78<br>(0.37-1.0) | 0.11<br>(0.04-0.14) | 0.77<br>(0.59-0.96) | 0.10<br>($5.0 \times 10^{-4}$ - 0.26) | 8.9<br>(1.6-15.0) | 46.0<br>(39.0-49.9) |

## Notes

### Competing Interest Statement

The authors have declared no competing interest.

### Summary of Updates

An updated version of the blackbird population data has become available. We have reran all analyses using this updated dataset. While some figures and numbers have slightly changed, the main findings and conclusions remain unchanged.

## References

[1] Paull SH, Song S, McClure KM, Sackett LC, Kilpatrick AM, Johnson PTJ. From superspreaders to disease hotspots: linking transmission across hosts and space. Front Ecol Environ 2011;10:75. 10.1890/110111.

[2] Weaver SC, Barrett ADT. Transmission cycles, host range, evolution and emergence of arboviral disease. Nat Rev Microbiol 2004;2:789–801. 10.1038/nrmicro1006.

[3] Munnink BBO, Sikkema RS, Nieuwenhuijse DF, Molenaar RJ, Munger E, Molenkamp R, et al. Transmission of SARS-CoV-2 on mink farms between humans and mink and back to humans. Science (1979) 2021;371:172–7. 10.1126/SCIENCE.ABE5901/SUPPL_FILE/ABE5901_OUDE_MUNNINK_TABLE_S1.PDF.

[4] Brown VL, Drake JM, Stallknecht DE, Brown JD, Pedersen K, Rohani P. Dissecting a wildlife disease hotspot: the impact of multiple host species, environmental transmission and seasonality in migration, breeding and mortality. J R Soc Interface 2013;10. 10.1098/RSIF.2012.0804.

[5] Crispell J, Zadoks RN, Harris SR, Paterson B, Collins DM, de-Lisle GW, et al. Using whole genome sequencing to investigate transmission in a multi-host system: Bovine tuberculosis in New Zealand. BMC Genomics 2017;18:1–12. 10.1186/S12864-017-3569-X/TABLES/1.

[6] Webb CT, Brooks CP, Gage KL, Antolin MF. Classic flea-borne transmission does not drive plague epizootics in prairie dogs. Proc Natl Acad Sci U S A 2006;103:6236–41. 10.1073/PNAS.0510090103/SUPPL_FILE/INDEX.HTML.

[7] ten Bosch QA, Clapham HE, Lambrechts L, Duong V, Buchy P, Althouse BM, et al. Contributions from the silent majority dominate dengue virus transmission. PLoS Pathog 2018;14. 10.1371/JOURNAL.PPAT.1006965.

[8] Colman E, Holme P, Sayama H, Gershenson C. Efficient sentinel surveillance strategies for preventing epidemics on networks. PLoS Comput Biol 2019;15:e1007517. 10.1371/JOURNAL.PCBI.1007517.

[9] Powers KA, Ghani AC, Miller WC, Hoffman IF, Pettifor AE, Kamanga G, et al. The role of acute and early HIV infection in the spread of HIV and implications for transmission prevention strategies in Lilongwe, Malawi: a modelling study. The Lancet 2011;378:256–68. 10.1016/S0140-6736(11)60842-8.

[10] Streicker DG, Fenton A, Pedersen AB. Differential sources of host species heterogeneity influence the transmission and control of multihost parasites. Ecol Lett 2013;16:975–84. 10.1111/ELE.12122.

[11] Flasche S, Lipsitch M, Ojal J, Pinsent A. Estimating the contribution of different age strata to vaccine serotype pneumococcal transmission in the pre vaccine era: A modelling study. BMC Med 2020;18:1–12. 10.1186/S12916-020-01601-1/FIGURES/3.

[12] Beauvais W, Musallam I, Guitian J. Vaccination control programs for multiple livestock host species: An age-stratified, seasonal transmission model for brucellosis control in endemic settings the LCNTDR Collection: Advances in scientific research for NTD control. Parasit Vectors 2016;9:1–10. 10.1186/S13071-016-1327-6/FIGURES/4.

[13] Fitzpatrick MC, Hampson K, Cleaveland S, Meyers LA, Townsend JP, Galvani AP. Potential for Rabies Control through Dog Vaccination in Wildlife-Abundant Communities of Tanzania. PLoS Negl Trop Dis 2012;6:e1796. 10.1371/JOURNAL.PNTD.0001796.

[14] Buhnerkempe MG, Roberts MG, Dobson AP, Heesterbeek H, Hudson PJ, Lloyd-Smith JO. Eight challenges in modelling disease ecology in multi-host, multi-agent systems. Epidemics 2015;10:26–30. 10.1016/J.EPIDEM.2014.10.001.

[15] Orton RJ, Wright CF, Morelli MJ, Juleff N, Thébaud G, Knowles NJ, et al. Observing micro-evolutionary processes of viral populations at multiple scales. Philosophical Transactions of the Royal Society B: Biological Sciences 2013;368. 10.1098/RSTB.2012.0203.

[16] Rijks JM, Kik M, Slaterus R, Foppen R, Stroo A, Ijzer J, et al. Widespread Usutu virus outbreak in birds in The Netherlands, 2016. Eurosurveillance 2016;21. 10.2807/1560-7917.ES.2016.21.45.30391.

[17] Weissenböck H, Kolodziejek J, Url A, Lussy H, Rebel-Bauder B, Nowotny N. Emergence of Usutu virus, an African mosquito-borne flavivirus of the Japanese encephalitis virus group, central Europe. Emerg Infect Dis 2002;8:652–6. 10.3201/EID0807.020094.

[18] Angeloni G, Bertola M, Lazzaro E, Morini M, Masi G, Sinigaglia A, et al. Epidemiology, surveillance and diagnosis of Usutu virus infection in the EU/EEA, 2012 to 2021. Eurosurveillance 2023;28:2200929. 10.2807/1560-7917.ES.2023.28.33.2200929/CITE/REFWORKS.

[19] Montizaan M, Kik M, Rijks J, DWHC, Slaterus R, Schoppers J, et al. Nature Today | Opnieuw dode vogels door usutuvirus in Nederland; nog onverklaarde daling meldingen dode merels 2019. https://www.naturetoday.com/intl/nl/nature-reports/message/?msg=25571 (accessed August 24, 2020).

[20] Münger E, Atama NC, van Irsel J, Blom R, Krol L, van Mastrigt T, et al. One Health approach uncovers emergence and dynamics of Usutu and West Nile viruses in the Netherlands. Nature Communications 2025 16:1 2025;16:1–14. 10.1038/s41467-025-63122-w.

[21] Van Irsel J, van der Jeugd HP, de Boer WF, Matson KD, van den Brand JMA, Sikkema R, et al. Spatio-temporal Usutu virus model explains Eurasian blackbird Turdus merula population trends. Ecography 2025;2025:e07759. 10.1111/ECOG.07759.

[22] Nikolay B. A review of West Nile and Usutu virus co-circulation in Europe: how much do transmission cycles overlap? Trans R Soc Trop Med Hyg 2015;109:609–18. 10.1093/TRSTMH/TRV066.

[23] Becker N, Jöst H, Ziegler U, Eiden M, Höper D, Emmerich P, et al. Epizootic Emergence of Usutu Virus in Wild and Captive Birds in Germany. PLoS One 2012;7:e32604. 10.1371/JOURNAL.PONE.0032604.

[24] Agliani G, Visser I, Marshall EM, Giglia G, de Bruin E, Verstappen R, et al. Experimental Usutu virus infection in Eurasian blackbirds (Turdus merula). Npj Viruses 2025;3:51. 10.1038/S44298-025-00133-W.

[25] Krol L, Dellar M, Ibáñez-Justicia A, Schrier G van der, Schrama M, Geerling GW, et al. Combined effects of future climate and land use change on mosquitoes: The distribution of Culex pipiens under One Health scenarios in the Netherlands 2024. 10.21203/RS.3.RS-5298493/V1.

[26] Dellar M, Sierdsema H, Schrama M, Geerling G, van Bodegom PM. The Future Abundance of Key Bird Species for Pathogen Transmission in the Netherlands. EcoHealth 2025 2025:1–17. 10.1007/S10393-025-01727-9.

[27] Temperature - gridded daily mean temperature in the Netherlands. KNMI 2015. https://dataplatform.knmi.nl/dataset/tg1-5 (accessed September 26, 2024).

[28] Marini G, Rosá R, Pugliese A, Heesterbeek H. Exploring vector-borne infection ecology in multi-host communities: A case study of West Nile virus. J Theor Biol 2017;415:58–69. 10.1016/J.JTBI.2016.12.009.

[29] Simpson JE, Hurtado PJ, Medlock J, Molaei G, Andreadis TG, Galvani AP, et al. Vector host-feeding preferences drive transmission of multi-host pathogens: West Nile virus as a model system. Proc Biol Sci 2012;279:925–33. 10.1098/RSPB.2011.1282.

[30] Cruz-Pacheco G, Esteva L, Vargas C. Multi-species interactions in West Nile virus infection. J Biol Dyn 2012;6:281–98. 10.1080/17513758.2011.571721.

[31] Robertson SL, Caillouët KA. A host stage-structured model of enzootic West Nile virus transmission to explore the effect of avian stage-dependent exposure to vectors. J Theor Biol 2016;399:33–42. 10.1016/J.JTBI.2016.03.031.

[32] Maidana NA, Yang HM. Dynamic of West Nile Virus transmission considering several coexisting avian populations. Math Comput Model 2011;53:1247–60. 10.1016/J.MCM.2010.12.008.

[33] Beebe TA, Robertson SL. A two-species stage-structured model for West Nile virus transmission. Lett Biomath 2017;4:112–32. 10.30707/LIB4.1BEEBE.

[34] Blom R, Krol L, Langezaal M, Schrama M, Trimbos KB, Wassenaar D, et al. Blood-feeding patterns of Culex pipiens biotype pipiens and pipiens/molestus hybrids in relation to avian community composition in urban habitats. Parasit Vectors 2024;17:1–12. 10.1186/S13071-024-06186-9/FIGURES/4.

[35] Rizzoli A, Bolzoni L, Chadwick EA, Capelli G, Montarsi F, Grisenti M, et al. Understanding West Nile virus ecology in Europe: Culex pipiens host feeding preference in a hotspot of virus emergence. Parasit Vectors 2015;8. 10.1186/s13071-015-0831-4.

[36] Fine P, Eames K, Heymann DL. “Herd Immunity”: A Rough Guide. Clinical Infectious Diseases 2011;52:911–6. 10.1093/CID/CIR007.

[37] Measles vaccines: WHO position paper – April 2017 n.d. https://www.who.int/publications/i/item/who-wer9217-205-227 (accessed September 8, 2025).

[38] Sæther BE. Pattern of covariation between life-history traits of European birds. Nature 1988;331:616–7. 10.1038/331616A0;KWRD=SCIENCE.

[39] Meister T, Lussy H, Bakonyi T, Šikutová S, Rudolf I, Vogl W, et al. Serological evidence of continuing high Usutu virus (Flaviviridae) activity and establishment of herd immunity in wild birds in Austria. Vet Microbiol 2008;127:237–48. 10.1016/J.VETMIC.2007.08.023.

[40] Magallanes S, Llorente F, Ruiz-López MJ, Puente JM de la, Ferraguti M, Gutiérrez-López R, et al. Warm winters are associated to more intense West Nile virus circulation in southern Spain. Emerg Microbes Infect 2024;13. 10.1080/22221751.2024.2348510.

[41] De Nardi A, Marini G, Dorigatti I, Rosà R, Tamba M, Gelmini L, et al. Quantifying West Nile virus circulation in the avian host population in Northern Italy. Infect Dis Model 2025;10:375–86. 10.1016/J.IDM.2024.12.009.

[42] Streng K, Atama N, Chandler F, Blom R, van der Jeugd H, Schrama M, et al. Sentinel chicken surveillance reveals previously undetected circulation of West Nile virus in the Netherlands. Emerg Microbes Infect 2024;13. 10.1080/22221751.2024.2406278.

[43] de Bruin E, van Irsel J, Chandler F, Kohl R, van de Voorde T, van der Linden A, et al. Usutu Virus Antibody Dynamics in Naturally Infected Blackbirds, the Netherlands, 2016–2018 - Volume 31, Number 6—June 2025 - Emerging Infectious Diseases journal - CDC. Emerg Infect Dis 2025;31:1244–6. 10.3201/EID3106.241744.

[44] Fink D, Auer T, Johnston A, Strimas-Mackey M, Ligocki S, Robinson O, et al. eBird Status and Trends, Data Version 2022. EBird 2023. 10.2173/EBIRDST.2022.

[45] Escribano-Romero E, Jiménez de Oya N, Camacho MC, Blázquez AB, Martín-Acebes MA, Risalde MA, et al. Previous Usutu Virus Exposure Partially Protects Magpies (Pica pica) against West Nile Virus Disease But Does Not Prevent Horizontal Transmission. Viruses 2021;13. 10.3390/V13071409.

[46] Magpie | BTO - British Trust for Ornithology n.d. https://www.bto.org/understanding-birds/birdfacts/magpie (accessed May 8, 2024).

[47] Coombs CFB, Isaacson AJ, Murton RK, Thearle RJP, Westwood NJ. Collared Doves (Streptopelia decaocto) in Urban Habitats. J Appl Ecol 1981;18:41. 10.2307/2402478.

[48] Yaremych SA, Novak RJ, Raim AJ, Mankin PC, Warner RE. HOME RANGE AND HABITAT USE BY AMERICAN CROWS IN RELATION TO TRANSMISSION OF WEST NILE VIRUS. Https://DoiOrg/101676/03-104 2004;116:232–9. 10.1676/03-104.

[49] Kuchinsky SC, Marano J, Hawks SA, Loessberg E, Honaker CF, Siegel PB, et al. North American House Sparrows Are Competent for Usutu Virus Transmission. MSphere 2022. 10.1128/MSPHERE.00295-22.

[50] Benzarti E, Rivas J, Sarlet M, Franssen M, Desmecht D, Schmidt-Chanasit J, et al. Experimental usutu virus infection in domestic canaries serinus canaria. Viruses 2020;12. 10.3390/v12020164.

[51] Cadar D, Lühken R, van der Jeugd H, Garigliany M, Ziegler U, Keller M, et al. Widespread activity of multiple lineages of Usutu virus, Western Europe, 2016. Eurosurveillance 2017;22. 10.2807/1560-7917.ES.2017.22.4.30452.

[52] Sieg M, Schmidt V, Ziegler U, Keller M, Höper D, Heenemann K, et al. Outbreak and Cocirculation of Three Different Usutu Virus Strains in Eastern Germany. Vector-Borne and Zoonotic Diseases 2017;17:662–4. 10.1089/VBZ.2016.2096/ASSET/IMAGES/LARGE/FIGURE1.JPEG.

[53] Young JJ, Haussig JM, Aberle SW, Pervanidou D, Riccardo F, Sekulić N, et al. Epidemiology of human West Nile virus infections in the European Union and European Union enlargement countries, 2010 to 2018. Eurosurveillance 2021;26:2001095. 10.2807/1560-7917.ES.2021.26.19.2001095/CITE/REFWORKS.

[54] Visser ME, Both C, Lambrechts MM. Global Climate Change Leads to Mistimed Avian Reproduction. Adv Ecol Res 2004;35:89–110. 10.1016/S0065-2504(04)35005-1.

[55] Sauer FG, Timmermann E, Lange U, Lühken R, Kiel E. Effects of Hibernation Site, Temperature, and Humidity on the Abundance and Survival of Overwintering Culex pipiens pipiens and Anopheles messeae (Diptera: Culicidae). J Med Entomol 2022;59:2013–21. 10.1093/JME/TJAC139.

[56] De Angelis D, Presanis AM, Birrell PJ, Tomba GS, House T. Four key challenges in infectious disease modelling using data from multiple sources. Epidemics 2015;10:83–7. 10.1016/J.EPIDEM.2014.09.004.

[57] Wonham MJ, De-Camino-Beck T, Lewis MA. An epidemiological model for West Nile virus: Invasion analysis and control applications. Proceedings of the Royal Society B: Biological Sciences 2004;271:501–7. 10.1098/rspb.2003.2608.

[58] Anderson RM, May RM. Infectious Diseases of Humans: Dynamics and Control. OUP Oxford; 1991.

[59] Paradis E, Baillie SR, Sutherland WJ, Gregory RD. Patterns of natal and breeding dispersal in birds. Journal of Animal Ecology 1998;67:518–36. 10.1046/J.1365-2656.1998.00215.X.

[60] Boulinier T, Kada S, Ponchon A, Dupraz M, Dietrich M, Gamble A, et al. Migration, Prospecting, Dispersal? What Host Movement Matters for Infectious Agent Circulation? Integr Comp Biol 2016;56:330–42. 10.1093/ICB/ICW015.

[61] Van Vliet J, Musters CJM, Ter Keurs WJ. Changes in migration behaviour of Blackbirds Turdus merula from the Netherlands. Bird Study 2009;56:276–81. 10.1080/00063650902792148.

[62] Hamer GL, Anderson TK, Donovan DJ, Brawn JD, Krebs BL, Gardner AM, et al. Dispersal of Adult Culex Mosquitoes in an Urban West Nile Virus Hotspot: A Mark-Capture Study Incorporating Stable Isotope Enrichment of Natural Larval Habitats. PLoS Negl Trop Dis 2014;8:e2768. 10.1371/JOURNAL.PNTD.0002768.

[63] Schoppers J, van Turnhout C, van Diek H. Handleiding Meetnet Urbane Soorten (MUS).. 2020.

[64] Teunissen WA, Wiersma P, de Jong A, Kleyheeg E, Vergeer J-W. Handleiding voor het Meetnet Agrarische Soorten.. 2019.

[65] Vergeer JW, Boele A, van Bruggen J, van Turnhout C. Handleiding Sovon Broedvogelmonitoring: Broedvogel Monitoring Project en kolonievogels. Sovon Vogelonderzoek Nederland, Nijmegen: 2023.

[66] Vigie-Nature. Base de données du Suivi Temporel des Oiseaux Communs / Common Bird Monitoring Scheme database for France. 2021.

[67] Ibañez-Justicia A, Stroo A, Dik M, Beeuwkes J, Scholte EJ. National mosquito (Diptera: Culicidae) survey in the Netherlands 2010-2013. J Med Entomol 2015;52:185–98. 10.1093/jme/tju058.

[68] Bortel W Van, Versteirt V, Dekoninck W, Hance T, Brosens D, Hendrickx G. MODIRISK: Mosquito vectors of disease, collection, monitoring and longitudinal data from Belgium. GigaByte 2022;2022:gigabyte58. 10.46471/GIGABYTE.58.

[69] Beaunée G. Batched Resilient and Rapid Estimation Workflow through Approximate Bayesian Computation. R package version 1.0.0. 2024. https://github.com/GaelBn/BRREWABC/.

[70] Sisson SA, Fan Y, Tanaka MM. Sequential Monte Carlo without likelihoods. Proc Natl Acad Sci U S A 2007;104:1760–5. 10.1073/PNAS.0607208104/ASSET/92D3CBC5-E270-4860-A408-2A7840ED8049/ASSETS/GRAPHIC/ZPQ00607-4886-M09.JPEG.

[71] Toni T, Welch D, Strelkowa N, Ipsen A, Stumpf MPH. Approximate Bayesian computation scheme for parameter inference and model selection in dynamical systems. J R Soc Interface 2008;6:187–202. 10.1098/RSIF.2008.0172.

[72] Diekmann O, Heesterbeek JAP, Roberts MG. The construction of next-generation matrices for compartmental epidemic models. J R Soc Interface 2010;7:873–85. 10.1098/RSIF.2009.0386.

[73] Boele A, Vergeer J, Van Bruggen J, Goffin B, Kavelaars M, Louwe Kooijmans J, et al. Broedvogels in Nederland in 2022. Sovon-rapport 2023/40. 2023.

[74] Birdfact. Blackbird Nesting: A Complete Guide | Birdfact. BirdfactCom 2023. https://birdfact.com/articles/blackbird-nesting (accessed December 7, 2023).

[75] van den Bremer L, van Turnhout C. Voorstudie Jaar van de Merel 2022. Sovon-rapport 2021/56. Nijmegen: 2021.

[76] Robinson RA, Kew JJ, Kew AJ. Survival of suburban blackbirds Turdus merula varies seasonally but not by sex. J Avian Biol 2010;41:83–7. 10.1111/J.1600-048X.2009.04789.X.

[77] Ewing DA, Cobbold CA, Purse B V., Nunn MA, White SM. Modelling the effect of temperature on the seasonal population dynamics of temperate mosquitoes. J Theor Biol 2016;400:65–79. 10.1016/J.JTBI.2016.04.008.

[78] Denlinger DL, Armbruster PA. Mosquito diapause. Annu Rev Entomol 2014;59:73–93. 10.1146/ANNUREV-ENTO-011613-162023/1.

[79] UK: average temperature by month 2023 | Statista n.d. https://www.statista.com/statistics/322658/monthly-average-daily-temperatures-in-the-united-kingdom-uk/ (accessed May 2, 2024).

[80] KNMI - Maandgemiddelde temperaturen, normalen, anomalieën n.d. https://www.knmi.nl/nederland-nu/klimatologie/geografische-overzichten/archief/maand/tg (accessed May 2, 2024).

[81] Perkins TA, Reiner RC, España G, Ten Bosch QA, Verma A, Liebman KA, et al. An agent-based model of dengue virus transmission shows how uncertainty about breakthrough infections influences vaccination impact projections. PLoS Comput Biol 2019;15:e1006710. 10.1371/journal.pcbi.1006710.

[82] Rubel F, Brugger K, Hantel M, Chvala-Mannsberger S, Bakonyi T, Weissenböck H, et al. Explaining Usutu virus dynamics in Austria: Model development and calibration. Prev Vet Med 2008;85:166–86. 10.1016/j.prevetmed.2008.01.006.

[83] Nicholas Komar. West Nile Virus: Epidemiology and Ecology in North America, 2003, p. 185–234. 10.1016/S0065-3527(03)61005-5.

[84] McKee EM, Walker ED, Anderson TK, Kitron UD, Brawn JD, Krebs BL, et al. WEST NILE VIRUS ANTIBODY DECAY RATE IN FREE-RANGING BIRDS. J Wildl Dis 2015;51:601–8. 10.7589/2014-07-175.

[85] Shocket MS, Verwillow AB, Numazu MG, Slamani H, Cohen JM, El Moustaid F, et al. Transmission of west nile and five other temperate mosquito-borne viruses peaks at temperatures between 23°C and 26°C. Elife 2020;9:1–67. 10.7554/ELIFE.58511.

[86] Ciota AT, Matacchiero AC, Kilpatrick AM, Kramer LD. The Effect of Temperature on Life History Traits of Culex Mosquitoes. J Med Entomol 2014;51:55–62. 10.1603/ME13003.

[87] Ruybal JE, Kramer LD, Kilpatrick AM. Geographic variation in the response of Culex pipiens life history traits to temperature. Parasit Vectors 2016;9:116. 10.1186/s13071-016-1402-z.

[88] Andreadis SS, Dimotsiou OC, Savopoulou-Soultani M. Variation in adult longevity of Culex pipiens f. pipiens, vector of the West Nile Virus. Parasitol Res 2014;113:4315–9. 10.1007/S00436-014-4152-X/TABLES/1.

[89] Reisen WK, Fang Y, Martinez VM. Effects of Temperature on the Transmission of West Nile Virus by Culex tarsalis (Diptera: Culicidae). vol. 43. 2006.

[90] Dohm DJ, Sardelis MR, Turell MJ. Experimental Vertical Transmission of West Nile Virus by Culex pipiens (Diptera: Culicidae). J Med Entomol 2002;39:640–4. 10.1603/0022-2585-39.4.640.

[91] Kilpatrick AM, Meola MA, Moudy RM, Kramer LD. Temperature, viral genetics, and the transmission of West Nile virus by Culex pipiens mosquitoes. PLoS Pathog 2008;4. 10.1371/JOURNAL.PPAT.1000092.

[92] Field EN, Shepard JJ, Clifton ME, Price KJ, Witmier BJ, Johnson K, et al. Semi-field and surveillance data define the natural diapause timeline for Culex pipiens across the United States. Communications Biology 2022 5:1 2022;5:1–12. 10.1038/s42003-022-04276-x.

[93] Widgren S, Bauer P, Eriksson R, Engblom S. Siminf: An R package for data-driven stochastic disease spread simulations. J Stat Softw 2019;91:1–42. 10.18637/jss.v091.i12.

[94] Da Re D, Marini G, Bonannella C, Laurini F, Manica M, Anicic N, et al. Modelling the seasonal dynamics of Aedes albopictus populations using a spatio-temporal stacked machine learning model. Sci Rep 2025;15:3750. 10.1038/S41598-025-87554-Y;SUBJMETA=1144,1469,158,704,852;KWRD=BIOGEOGRAPHY,ECOLOGICAL+EPIDEMIOLOGY,ECOLOGICAL+MODELLING.

[95] Davis BE. Habitat use, movements, and survival of radio-marked female mallards in the lower Mississippi Alluvial Valley. LSU Master’s Theses 2007. 10.31390/gradschool_theses.215.

[96] de Bellegarde de Saint Lary C, Kasbergen LMR, Bruijning-Verhagen PCJL, van der Jeugd H, Chandler F, Hogema BM, et al. Assessing West Nile virus (WNV) and Usutu virus (USUV) exposure in bird ringers in the Netherlands: a high-risk group for WNV and USUV infection? One Health 2023;16:100533. 10.1016/J.ONEHLT.2023.100533.

[97] Cleton NB, Godeke GJ, Reimerink J, Beersma MF, Doorn HR van, Franco L, et al. Spot the Difference—Development of a Syndrome Based Protein Microarray for Specific Serological Detection of Multiple Flavivirus Infections in Travelers. PLoS Negl Trop Dis 2015;9:e0003580. 10.1371/JOURNAL.PNTD.0003580.

[98] Dutch Wildlife Health Centre (DWHC) & Sovon. Dode vogels melden. Https://DwhcNl/Publicaties/#informatie-En-Downloads 2023.

[99] Del Moral P, Doucet A, Jasra A. Sequential Monte Carlo samplers. J R Stat Soc Series B Stat Methodol 2006;68:411–36. 10.1111/J.1467-9868.2006.00553.X.

[100] Del Moral P, Doucet A, Jasra A. An adaptive sequential Monte Carlo method for approximate Bayesian computation. Stat Comput 2012;22:1009–20. 10.1007/S11222-011-9271-Y.

[101] Drovandi CC, Pettitt AN. Estimation of Parameters for Macroparasite Population Evolution Using Approximate Bayesian Computation. Biometrics 2011;67:225–33. 10.1111/J.1541-0420.2010.01410.X.

[102] Beaumont MA, Cornuet JM, Marin JM, Robert CP. Adaptive approximate Bayesian computation. Biometrika 2009;96:983–90. 10.1093/BIOMET/ASP052.

[103] Adams B, Boots M. How important is vertical transmission in mosquitoes for the persistence of dengue? Insights from a mathematical model. Epidemics 2010;2:1–10. 10.1016/j.epidem.2010.01.001.

[104] Turell MJ, O’Guinn ML, Dohm DJ, Jones JW. Vector competence of North American mosquitoes (Diptera: Culicidae) for West Nile virus. J Med Entomol 2001;38:130–4. 10.1603/0022-2585-38.2.130.

